# Value and choice as separable, stable representations in orbitofrontal cortex

**DOI:** 10.1101/2019.12.31.892109

**Authors:** Daniel L. Kimmel, Gamaleldin F. Elsayed, John P. Cunningham, William T. Newsome

## Abstract

Value-based decision-making operates on multiple variables—including offer value, choice, expected outcome, and recent history—each functioning at different times in the decision process. Orbitofrontal cortex (OFC) has long been implicated in value-based decision-making, but it is unclear how downstream circuits might read out complex OFC responses into separate representations of the relevant variables to support different cognitive functions at specific times. We recorded from single neurons in OFC while macaque monkeys made cost-benefit decisions to juice offers. Using a novel analysis—optimal targeted dimensionality reduction—we discovered orthogonal, static dimensions (i.e. linear combinations of neurons) that selectively represented the value, choice, and expected reward of the present and, separately, previous offers. The neural composition of most representations was stable over discrete time periods that aligned to concurrent cognitive demands. We applied a new set of statistical methods to determine that the sensitivity, specificity and stability of the representations were greater than expected from the low-level features—dimensionality and temporal smoothness—of the responses alone. The separability and stability of OFC representations suggest a mechanism by which downstream circuits can read out specific task-relevant variables at appropriate times.

## Introduction

Value-based decision-making is an essential component of behavior, and its neural basis has been an area of intense scientific inquiry^1–3^. Economists, psychologists and neuroscientists have long-posited a need for separable representations of key decision variables—including 1) initial valuation of the offer, 2) chosen action, and 3) expected outcome given the choice—that in turn support respective steps in the decision process: 1) committing to an initial behavioral policy, 2) computing expected outcome while executing the policy and then later assigning credit to the chosen action, and 3) reevaluating the policy mid-execution and then later comparing expected and received outcomes to update future expectations^4^.

However, study of the neural basis of this process has faced four significant limitations. First, reports differ on how neural populations represent the key decision variables in a separable format. On one hand, some reports suggest that single neurons, particularly in orbitofrontal (OFC), represent task-relevant variables *categorically*, i.e., specializing for one variable over others^5–8^. In contrast, recordings during value-based tasks from an array of cortical areas—including OFC, dorsolateral prefrontal (PFC), anterior cingulate (ACC), and posterior cingulate cortices—observe that single neurons encode multiple task-relevant variables^9–11^. This phenomenon of *mixed selectivity* appears to be a general encoding phenomenon throughout the brain^12,13^. The question of categorical vs. mixed selectivity has profound implications for how downstream circuits read out population activity into separate decision variables that can drive distinct functions.

Second, the challenge of readout is compounded by the neural dynamics: the task-relevant variable(s) encoded by a given neuron or neural population often change over the course of the decision process^14,15^. Thus a readout selective for one variable at one point in time, may in fact select for an entirely different variable 100’s of milliseconds later. This is particularly true in OFC, which over the course of a single trial, appears to represent multiple decision variables, thus contributing to a plethora of behavioral functions ascribed to OFC^16^ and raising the critical question of how a single population supports multiple downstream functions.

Third, existing analyses of OFC obscure any mixed or dynamic selectivity. For instance, most studies report the percentages of neurons representing variables X or Y at a given time *t*_1_ and, separately, the percentages representing X or Y at a later time, *t*_2_. This approach implicitly assumes categorical selectivity—neurons represent *either* X *or* Y. Moreover, by analyzing each time point independently, this approach obscures how the contribution of individual neurons changes from *t*_1_ to *t*_2_. That is, even if the percentage remained constant, were the same neurons contributing at *t*_1_ and *t*_2_, and if so, to the same extent? Again, the implications for readout are profound. On one extreme, if each neuron’s contribution changed at each time point—as for the activity ‘sequences’ reported in rodent cortex^14,17^—then the readout must also adapt continuously. Conversely, if each neuron’s contribution were stable, then a static readout could capture the representation, but it would do so constitutively, unable to gate when a variable influenced downstream computation.

Though not previously applied to OFC, population-level analyses offer a means to demix single-neuron responses into weighted combinations of neurons—or low-dimensional representations—selective for single task-relevant variables^18–20^. However, these analyses must be coupled with a statistical framework to distinguish meaningful structure from mere epiphenomena, which often emerge in large populations of correlated neurons^21^. For instance, how sensitive would we expect a random dimension to be for a given variable? How much would a random dimension change over the course of a trial?

Fourth, despite the robust value signals observed in OFC, it remains unclear how and when these signals support value-based choice. As discussed above, value signals are likely important at different phases of the decision process. In most value-based studies, subjects render choices with brief, all-or-nothing responses (e.g., a reach, lever press, nose poke, or saccadic eye movement). In these tasks, value informs the decision, but because the operant response is brief and low-cost, has little influence on whether the choice is rendered. In contrast, most ethological decisions require an agent to execute a behavioral policy over time, often with sustained effort, until a desired outcome is reached (e.g., deciding to forage from a given fruit tree, then sustaining that policy while navigating to, climbing and competing with other animals in the tree)^22^. Agents may execute the policy adaptively—applying variable effort or even reversing the policy midway—depending on the value of the expected outcome. Doing so requires a sustained neural representation of value (in OFC or elsewhere), which may not be elicited—and its necessity cannot be tested—by the brief, all-or-nothing responses of most tasks.

Here, we present a novel behavioral task in which macaque monkeys traded sustained effort for an offered juice reward. As expected, animals were more likely to accept larger offers. However, animals did not commit to accepting an offer once per trial, but instead continued to reevaluate, and at times reverse, their initial choice mid-trial. We simultaneously recorded from single neurons in macaque OFC and found heterogeneous responses exhibiting mixed and dynamic selectivity. To capture this complexity, we applied a new dimensionality reduction technique and statistical model that computed separable population representations of the key task-relevant variables—offer size, choice, expected reward—for which the magnitude and specificity exceeded that expected by chance. The contribution of individual units to a given representation were stable throughout the task period when information was behaviorally relevant. However, between trials, task-relevant information abruptly transferred to a new set of dimensions, maintaining previous-trial information while distinguishing it from incoming present-trial information. The dynamics of the representations—abrupt transitions at key task events followed by stability during periods of behavioral relevance—suggested that OFC organized dynamically (i.e., forming and disbanding coordinated combinations of neurons) to represent task-relevant information at specific times. The low-dimensional, stable nature of the representations suggested a neurobiologically plausible mechanism—linear summation over static synaptic weights—by which downstream circuits could read out mixed, heterogeneous responses into separable representations for selectively driving specific behavioral functions at specific times.

## Methods

### Cost-benefit decision-making task

Two adult male macaque monkeys, N (*Macacca fascicularis*) and K (*M. mulatta*), served as subjects in this study. Prior to experimental use, each animal was prepared surgically with a head-holding device consisting of either a plastic cylinder embedded in acrylic^23^ (monkey N) or a titanium post secured directly to the skull^24^ (monkey K). During training and while engaged in experiments, daily fluid intake was restricted to maintain adequate levels of motivation; food was freely available. All surgical, behavioral, and animal care procedures complied with National Institutes of Health guidelines and were approved by the Stanford Institutional Animal Care and Use Committee.

Animals were seated in a primate chair in a sound-insulated and dimly lit chamber at a viewing distance of 43 cm (monkey N) or 55 cm (monkey K) from a 20” CRT computer monitor (ViewSonic G22fb, Walnut, CA) displaying 800 x 600 pixels at 96 Hz. Head position was stabilized using the head-holding device. Eye position was monitored at 1000 Hz with an infrared video tracker (EyeLink 1000, SR Research, Ontario, Canada) mounted on the primate chair with custom hardware; the real-time eye position signal was calibrated periodically^25^.

Behavioral control and stimulus presentation were managed by Apple Macintosh G5-based computers (Cupertino, CA) running Expo software written by Peter Lennie (University of Rochester, NY) with modifications by Julian Brown (Stanford University, CA). Behavioral and stimulus event data were acquired by the Plexon MAP System (Dallas, TX), while digital eye position samples were recorded natively on the EyeLink system.

We trained the animals on a novel cost-benefit decision-making task (Figure 1a). A trial began with the appearance of a fixation point (FP; white annulus, inner/outer diameter 0.3°/0.6°) against a dim background. The animal acquired the FP by directing its gaze within an invisible, circular *fixation window* around the FP (radius 3° or 1.8°, monkey N or K, respectively). If the animal failed to acquire the FP within 2 s, the FP was extinguished followed by a 1 s delay before reappearance of the FP. The animal was required to maintain fixation for a brief, variable period of time (*fixation* period; 0.5 - 1 s, uniformly distributed). Fixation breaks during the fixation period terminated the sequence (which was not scored as a “trial” in future analyses), and the task entered an *inter-trial interval* (ITI; see below).

**Figure 1.**
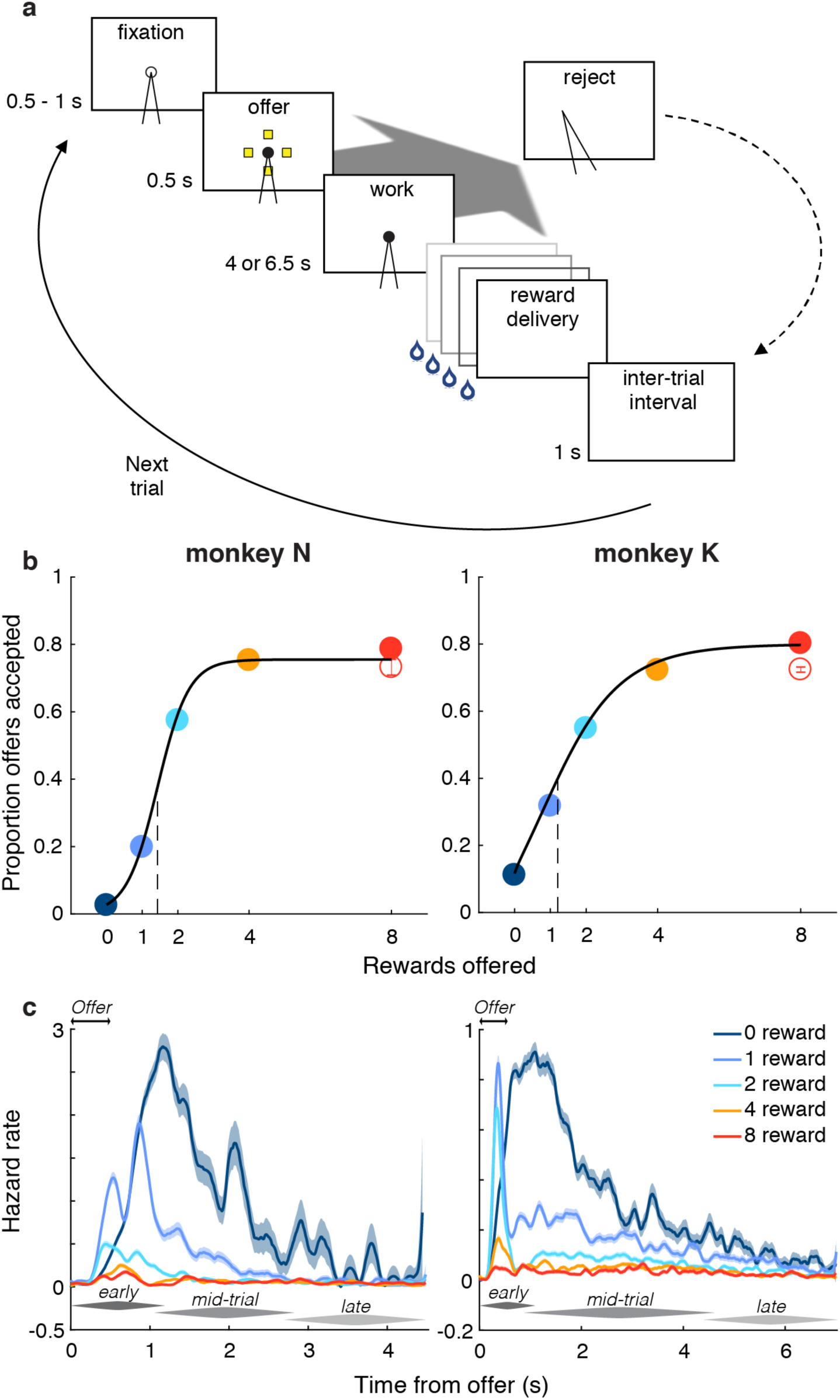
Behavioral design and results. **(a)** The time course of a trial is shown proceeding from top-left to bottom-right. The animal initiated the trial by directing its gaze toward an open circle, or fixation point (FP), and held fixation for a variable interval. An offer was presented as an array of icons (0, 1, 2, 4 or 8 yellow icons, selected randomly across trials) with each icon indicating a drop of juice in exchange for effortful work; simultaneously, the FP filled-in to a solid circle (providing a timing cue on 0-reward offers). To accept the offer, the animal maintained fixation through the work period (4 or 6.5 s, monkey N or K, respectively), after which the promised drops of juice were delivered in rapid succession. To reject the offer (large oblique gray arrow), the animal averted its gaze during the offer or work period. The trial then entered a time-out period equivalent in duration to the time that would have elapsed had the animal accepted the offer. The trial was followed by an inter-trial interval before advancing to the next trial (solid curve). **(b)** The proportion of offers accepted is shown as a function of offer size, which was conveyed to the animal either by the number of yellow icons (non-singleton offers; filled circles) or as single purple icon worth 8 drops (singleton offer; open circle). Error bars (generally smaller than circle) show variance of the proportion. Logistic function was fit to aggregated data for display purposes only (black curve); all statistical tests were performed on individual sessions. **(c)** Hazard rate of fixation breaks are shown for non-singleton offers (colored curves) with standard deviations (shading) as function of time from onset of the offer period (double arrows). Early, mid-trial, and late phases of rejections referred to in main text are indicated by dark, middle, and light tapered bars, respectively.

Following the fixation period, we presented the offer to the animal (*offer* period; 0.5 s) as a set of 0, 1, 2, 4 or 8 square icons (0.5° x 0.5°), evenly spaced along an invisible circular ring (6° radius) centered on the FP. The number and color of the icons indicated the *offer size*, or number of drops of juice the animal would receive as reward for accepting the offer, which varied pseudorandomly from trial to trial, with the constraint that all 5 offer sizes were presented twice every 10 trials. Most offers were presented with yellow icons, each of which represented a single drop of juice. However, to control for the correlation between visual stimulus properties and offer size, half of the 8-reward offers were presented with 8 yellow icons (*non*-*singleton* offer) and half with a single purple icon (*singleton* offer) for half or all experiments with monkey N or K, respectively. The animals learned the association between icons and rewards during training that preceded the current experiments. To control for particular orientations of the icons having a disproportionate effect on behavior or neural responses, we randomly rotated the array of icons around the invisible ring from trial-to-trial, always maintaining an equal angle between icons for a given offer. To provide a temporal cue for 0-reward offers (in which no icons were presented), the center of the FP was filled-in at the onset of the offer period for all offer sizes. (For approximately half of experiments for monkey N, the FP remained an annulus through the offer period, filling-in at the onset of the work period. The delayed FP transition did not affect the timing of responses; Supplementary Figure 3).

The offer period was followed by the *work* period. The offer icons were extinguished and the animal was required to maintain fixation for a constant period of time (4 or 6.5 s for monkey N or K, respectively) that was selected during training such that the animal rarely to maximally accepted the smallest to largest offers, respectively. If the animal maintained fixation until the end of the work period (i.e., *accepted* the offer), we extinguished the FP and delivered sequentially the drops of juice offered (*reward* period), with 0.4 s between each drop. (Drop size was regulated by the opening and closing of an electronic solenoid valve, which was calibrated regularly to achieve a constant drop size). To *reject* the offer, the animal simply averted its gaze any time during the offer or work periods. The trial then entered a timeout period whose duration was equal to that had the animal accepted the offer. During this timeout period, the screen went blank, and the animal was free to move its eyes, but no reward was delivered.

Following the period when rewards were or would have been delivered, the ITI was imposed. To prevent the animal from “zoning-out” and reflexively fixating continuously across a string of sequential trials, we required the animal to break fixation at some point during or after the current trial, i.e., any time during the offer or work periods (while the FP was present), or during the reward or ITI periods (during which the FP was absent, but the invisible fixation window was maintained). The ITI period was repeated until the required break occurred. In practice, the animals made numerous saccades during the ITI, and thus this task feature was not engaged outside of initial training. Following the ITI, the FP was presented for the next trial.

We defined the behavioral *conditions* as the unique combinations of five standard offers (i.e., 0, 1, 2, 4, or 8 rewards) and two choices (i.e., accept or reject the offer), plus the two choices in response to the 8-reward singleton offer, resulting in 12 possible conditions, though some conditions were more frequent than others (e.g., the animal rarely rejected 8-reward offers and rarely accepted 0-reward offers).

### Analysis of behavioral choice

All analyses described here and below were performed with custom scripts written in MATLAB (Mathworks, Natick, MA).

We modeled binary choice behavior to accept or reject the offer on trial *t* as the logistic function:

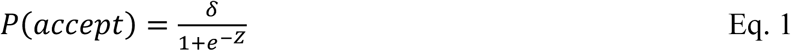

where 0 ≤ *δ* ≤ 1 specified the maximal accept rate, or saturation point of the psychometric curve. This parameter has previously been used to model the lapse rate, or intrinsic failure rate, of behavior^26^. The exponent *Z* took the form:

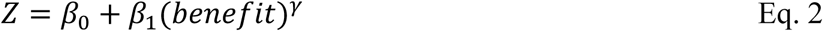

where *β*_0_ was a constant and *β*_1_ determined the influence of the offered benefit [0, 1, 2, 4, 8] on trial *t* raised to *γ*, an exponent typical in economic models to implement a non-linear utility function^27^ that was either fixed (*γ* = 1) or allowed to vary as a free parameter. The benefit predictor was scaled to the range [0, 1].

Maximum likelihood estimates for the free parameters *β*_0_, *β*_1_, *δ*, and *γ* were obtained independently for each experimental session (i.e., set of trials collected at a given recording site). Maximization was performed by the MATLAB function *fmincon* with constraint 0 ≤ *δ* ≤ 1. We considered choices in the normal stimulus condition (in which the number of yellow icons indicated the number of rewards) to represent most closely the animal’s cost-benefit function, and therefore excluded the singleton condition (in which 1 purple icon indicated 8 rewards) from all model fits. We took the similarity between the model estimate for the 8-reward normal stimulus and the behavior observed on singleton trials as an indicator of how well the animals had learned the value of the singleton stimulus.

We solved for two variations of the model, *a* or *b*, with *γ* as a free parameter or with *γ* = 1, respectively, obtaining the likelihood *L* of the data given the model. For each experimental session, we computed the log likelihood ratio (LR) for models *a* and *b*:

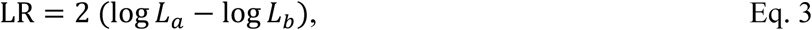

which, under the null hypothesis that the two models were equally likely, was *χ*^2^ distributed with df_*a*_ – df_*b*_ degrees of freedom, where df was the model’s number of free parameters. Finally, we computed the probability p(LR) of falsely rejecting the alternative hypothesis that the empirical cumulative distribution function of LR across sessions was right-shifted relative to the predicted *χ*^2^ cumulative distribution (i.e., 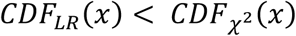 for all experimental sessions *x*) by the Kolmogorov-Smirnov test. We rejected the null hypothesis and selected the more complex model *a* to serve for all sessions when p(LR) was less than 0.05. For monkey N or K, LR = 0.53 or 0.63 and p(LR) = 0.36 or 0.037, respectively. Thus, we included *γ* as a free parameter for monkey K only and fixed *γ* = 1 for monkey N.

### Electrophysiological recording of OFC

In preparation for physiological recordings, each animal underwent anatomical magnetic resonance imaging (MRI) of the brain. During imaging, a rigid, MRI-visible fiducial marker was attached to the animal’s head and could be repositioned identically during subsequent surgery. Using the anatomical images, we identified ideal placement of the recording cylinder that accessed OFC in stereotactic planes. Each animal then underwent a surgical procedure in which a craniotomy was placed and a plastic recording cylinder (Crist Instrument, Hagerstown, MD) was positioned according to the MRI-guided coordinates and relative to the extracranial fiducial marker. The cylinders were centered at stereotaxic coordinates of 36.1 or 35.6 mm anterior to the interaural line, 5.2 or 8.9 mm left of midline, and angled 10.5° or 0.0° anterior to the coronal plane for monkey N or K, respectively. (Note that the reported cylinder location was based on intra-surgical stereotaxic measurements, while the medial-lateral position of individual recording sites, reported in Supplementary Figure 7, was measured from the MR images and accounted for the angle of the electrode trajectory.)

After the procedure, a small, cylindrical recording grid (Crist Instrument; 1 mm spacing between holes) was placed within the chamber and filled with salinized agarose solution that provided contrast for a second anatomical MRI prescribed such that the imaging planes were parallel (for coronal and sagittal sequences) or orthogonal (for axial sequences) to the recording grid holes. Thus, the final set of MR images shared the same planes as our eventual electrode penetrations, allowing us to visualize the electrode trajectory during each experiment and facilitating precise electrode placement within OFC (Supplementary Figure 4).

On the basis of previously reported value-related responses^15,28^, we concentrated our physiological recordings around the medial orbitofrontal sulcus (mOFS), including the medial and lateral banks and fundus, corresponding to Brodmann areas 11 and 13^29^. We employed standard methods to record the discharge of single units and ensembles of multiple units using extracellular tungsten microelectrodes (FHC Inc., Bowdoin, ME). For each experiment, we advanced the electrode with precision motors that were calibrated to provide a precise (< 10 μm) estimate of electrode position (NAN Instruments, Nazareth, Israel), while simultaneously advancing a virtual electrode in the co-registered MR images (Supplementary Figure 4). We found the correspondence between virtual and actual electrode position to be highly consistent (within 100 μm), as corroborated by the gray-white matter transitions. After waiting 10-20 minutes for the electrode to stabilize just superior to the mOFS, we slowly advanced the electrode while the animal performed the cost-benefit task, increasing the likelihood that relevant neurons would be active during the selection process. We selected for recording sites where we could isolate at least one single unit waveform and otherwise applied no additional criteria (taking “all comers”) so as to collect as representative a dataset of medial OFC as possible (26 or 86 sites, monkey N or K, respectively).

Once selecting a site for recording, we captured and stored all amplified waveforms—discrete excerpts of the continuous voltage time series—that exceeded a predetermined voltage threshold (set such that the event rate in white matter was 1 - 5 Hz) as digitized samples using the MAP data acquisition system (Plexon Inc., Dallas, TX), which simultaneously captured behavioral events relayed by the Expo software.

### Offline data processing and selection

All analyses were performed on waveforms sorted offline using specialized software (Offline Sorter, Plexon, Inc.) into single units (i.e., single neurons) and multi-units to which we collectively referred as “units”. The term “individual unit” referred to either one single unit or one multi-unit. We identified single units as waveforms whose morphology was stereotypical of single units, consistent over time, and easily distinguishable from other concurrent waveforms, and whose timing was separated by at least 1 ms (i.e., outside the minimum refractory period) and was at most weakly correlated with the timing of other waveforms (i.e., unlikely resulting from multiple threshold crossings from a single polyphasic waveform), as done typically in the literature^30^. The remaining, multi-unit waveforms were presumed to originate from two or more neurons. Waveforms deemed to be either electrical artifact or time-locked continuations of a previously counted waveform were excluded.

The final dataset included 26 and 131 single units and 42 and 211 multi-units from monkeys N and K, respectively. As reflected in the above counts, we excluded units that were recorded for fewer than 60 trials total (44 and 33 units, monkey N and K, respectively), had fewer than 5 trials in any of the 10 conditions (i.e., intersection of offer size and choice; 71 and 133 units), or for which the accompanying behavior was grossly aberrant (monkey K only: 1 session with accept rate of ∼55% for 0-reward offers and 1 session with ∼10% accept rate for all offers). When a unit was not present during a trial (i.e., mean spike rate across trial < 0.1 Hz) for 5 or more consecutive trials, we excluded those trials for that unit; this excluded both units with very low firing rates, as well as trials during which the unit may have been lost to recording. We computed the trial-average firing rate as the mean firing rate across trials in 100 ms time bins for each condition. Because our analysis would ultimately normalize a unit’s firing rate by its variability, we excluded 23 units (monkey K only) with extremely low variability (s.d. < 0.5 Hz) as measured across conditions and time bins so that imprecision in estimating variability (to which low-variability units were particularly susceptible) would not result in spurious over-weighting of these units. (Coincidentally, the mean firing rate of the low-variability units was also low, ranging from 0.072 to 0.76 Hz and with a population mean of 0.27 Hz compared to 29 Hz for the included population for monkey K.) Finally, because our analysis required that all included conditions were represented by all units, we excluded conditions for which less than 40% of units met the trial count threshold (i.e., 60 trials), including “0-reward, accept” for both animals and additionally “8-reward, reject” for monkey K.

To compute the trial-average response across *N* units, *C* conditions, and *T* times, we extracted the firing rate *R*_*n*_(*r, t*) for each unit *n* on trial *r* in the *t*^*th*^ non-overlapping, 100 ms time bin aligned to the time of the offer. We then computed the mean firing rate 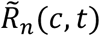, or trial-average response, across the trials of condition *c* (see next paragraph for condition definitions). To standardize the neural response, 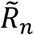 was z-transformed to 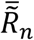 such that, for each unit, the mean and standard deviation were 0 and 1, respectively, across all times. Unless otherwise noted, we subtracted the mean response across conditions at each time *t* (i.e., common-condition response, CC_*n*_(*t*)) from 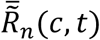, giving 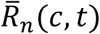. Finally, we compiled the responses 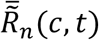 and 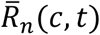 across units into tensors 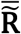 and 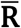, respectively, with dimensions *N* x *C* x *T*. As such, tensors 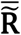 and 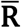 contained the standardized population response, with the responses in tensor 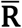 also being mean-subtracted.

We computed two sets of trial-average responses depending on whether the conditions *c* were defined by the present or previous trial. *Present-trial responses* were computed by sorting trials according to the present trial’s condition (i.e., 5 present offers * 2 present choices = 10 possible present-trial conditions; Figures 1-5 and right panels of 6a-d), while *previous-trial responses* were computed based on the previous trial’s condition (i.e., 5 previous offers * 2 previous choices; left panels of Figure 6a-d and all panels of 6e-f). (Note: the 8-reward singleton condition was excluded when computing population-level dimensions, see below; however, its trial-average response was computed and normalized identically as for the other conditions and then projected onto the dimensions for certain analyses, e.g., Figure 4b.) To understand the rationale for the two sorting schemes, consider the trial-average responses of a given unit computed according to the present conditions, such as the example unit in Figure 2a that strongly differentiated sizes of the present offer. However, if one followed these trial-average responses back in time, they would presumably overlap entirely during the previous trial because, given the random trial order, included in any one present-trial response were all 10 previous-trial conditions! However, if the same unit’s trial-level responses were resorted according to the conditions on the *previous* trial, we would again observe differential responses to the different levels of the previous offer *during the previous trial*.

**Figure 2.**
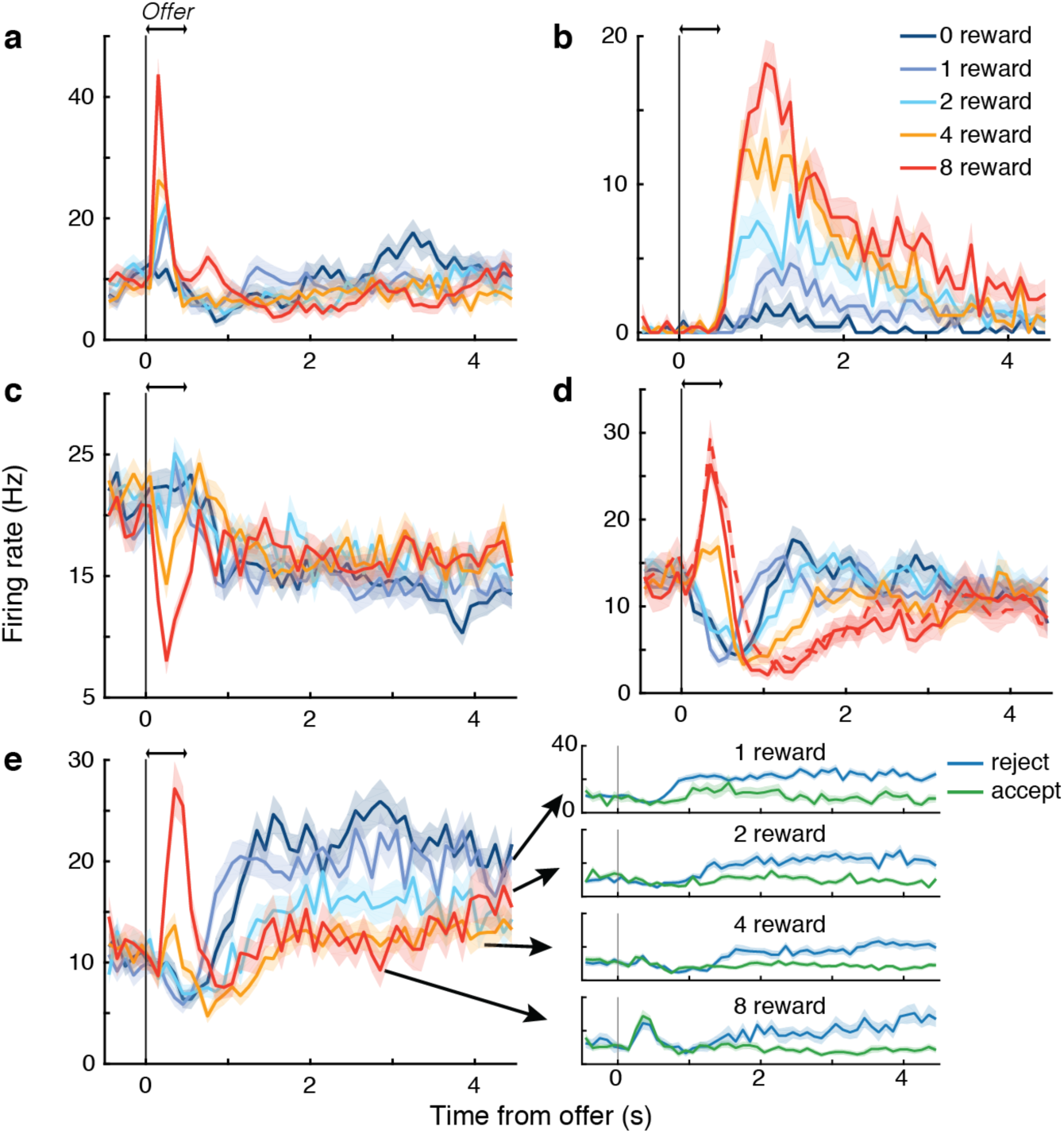
Dynamic encoding and mixed selectivity in individual units. **(a-e)** Mean response to the non-singleton (solid curves) and singleton (dashed curve) offers, with associated s.e.m. (shading), is shown as a function of time from onset of the offer period (double arrows) for five example units. (Singleton condition was not present in all sessions. Averages include all choices. Further stratifying by accept (green) and reject (blue) choices (e) showed selectivity for both benefit and choice. While the encoding of offer size appeared to reverse for unit (e), this was an artifact of the greater response for reject than accept choices later in the trial combined with more frequent rejections for smaller offers. In contrast, the reversal observed for unit (d) did not depend on choice (not shown).

**Figure 3.**
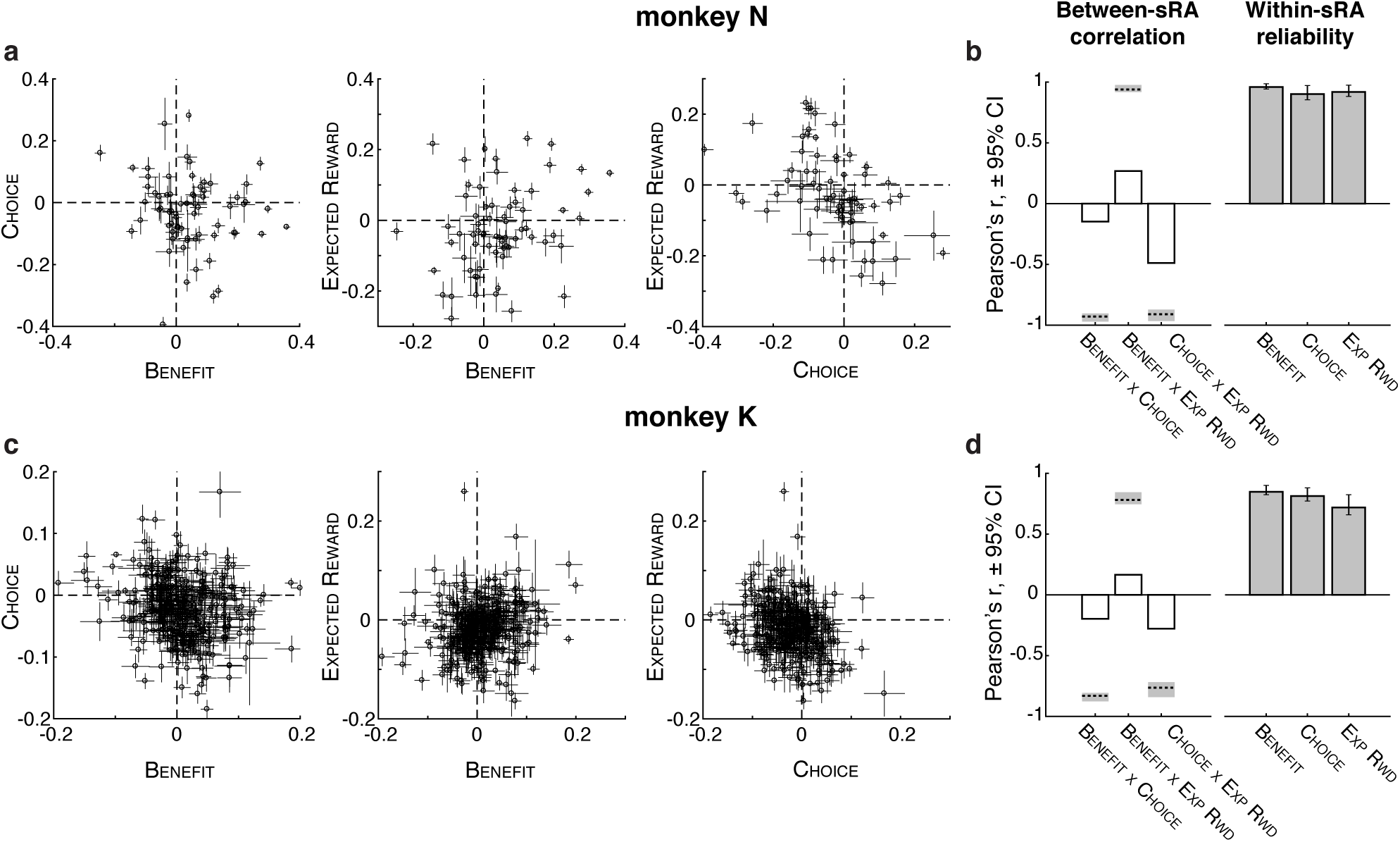
Contribution of individual units to low-dimensional representations. **(a**,**c)** The pairwise relationship between regression coefficients for task-relevant variables (abscissa and ordinate labels) are shown for each unit (open circles) with standard deviations (horizontal and vertical error bars; computed by resampling trial-level responses, see Methods), for monkeys N (a) and K (c). If units generally encoded a solitary variable, circles would cluster on the horizontal and vertical meridians (dashed lines), which was not observed, consistent with mixed selectivity. If estimates of the regression coefficients were unreliable, then apparent mixed selectivity could arise spuriously from imprecise estimates and error bars would consistently overlap the meridians, which was not observed. **(b**,**d)** The Pearson’s correlation coefficient between pairs of representations (i.e., sRAs) of different (open bars) or same (filled bars) variable(s) are shown for monkeys N (b) and K (d). For between-sRA correlations, the coefficient was measured from the full dataset; dashed horizontal line and shading indicate the mean and 95% confidence interval (CI), respectively, of the hypothetical correlation between two perfectly correlated (or anti-correlated) representations corrupted by independent noise (i.e., low precision of the individual-unit coefficients), as measured across resampled datasets. Observed correlations were significantly closer to zero than these hypothetical values, defining the representations as separable (Supplementary Figure 8 and Supplementary Table 2). For within-sRA correlations, the coefficient was computed as the mean (bar height) and 95% CI (error bar) across resampled datasets. For present figure, all regression coefficients pertain to non-orthogonalized sRAs, which most closely reflected the population encoding and permitted meaningful comparison between representations (in contrast, the correlation between orthogonalized sRAs would necessarily be zero).

**Figure 4.**
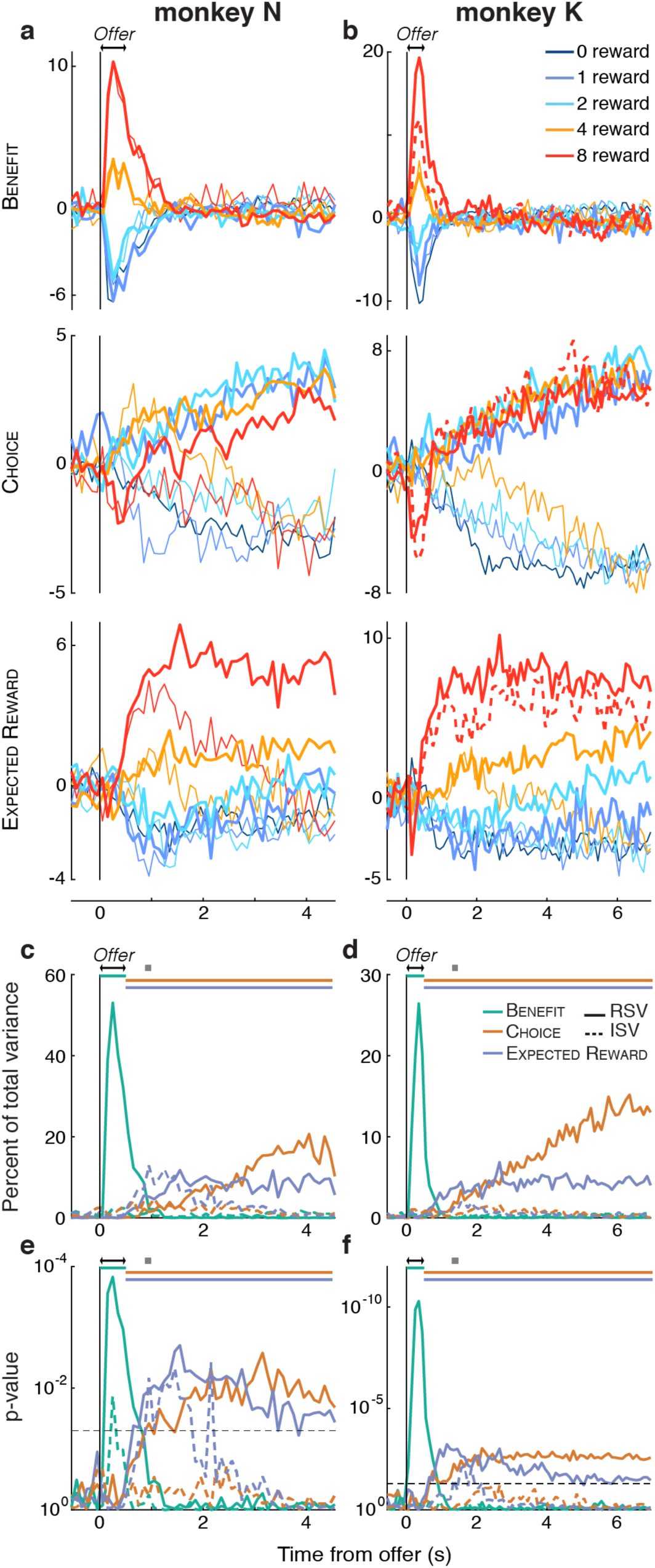
Activity of and variance explained by low-dimensional representations. **(a**,**b)** The time course of the high-dimensional neural activity projected onto the low-dimensional representations (i.e., “activity” of the sRA) of BENEFIT (top), CHOICE (middle), and EXPECTED REWARD (bottom) is shown in arbitrary units for each offer size (colors) and accept and reject choices (thick and thin curves, respectively) as a function of time from the onset of the offer period (double arrows) for monkeys N (a) and K (b). The singleton condition (dashed curve) is shown here for monkey K and Supplementary Figure 11 for monkey N, but was not used to discover the representations. Note that some combinations of offer and choice (e.g., zero-reward, accept choices) had too few trials per unit to accurately estimate trial-average responses and were excluded (see Methods). **(c**,**d)** The relevant and irrelevant signal variance (solid and dashed curves, respectively) for BENEFIT (green), CHOICE (orange), and EXPECTED REWARD (blue) are shown as a percentage of total cross-condition variance as a function of time from the onset of the offer period (double arrows) for monkeys N (c) and K (d). **(e**,**f)** Log_10_ probability of data in (c) and (d), respectively. Horizontal dashed line corresponds to p = 0.05. In (c-f), gray squares indicate the median rejection time and colored horizontal bars span the temporal epoch in which the color-matched sRAs were computed.

**Figure 5.**
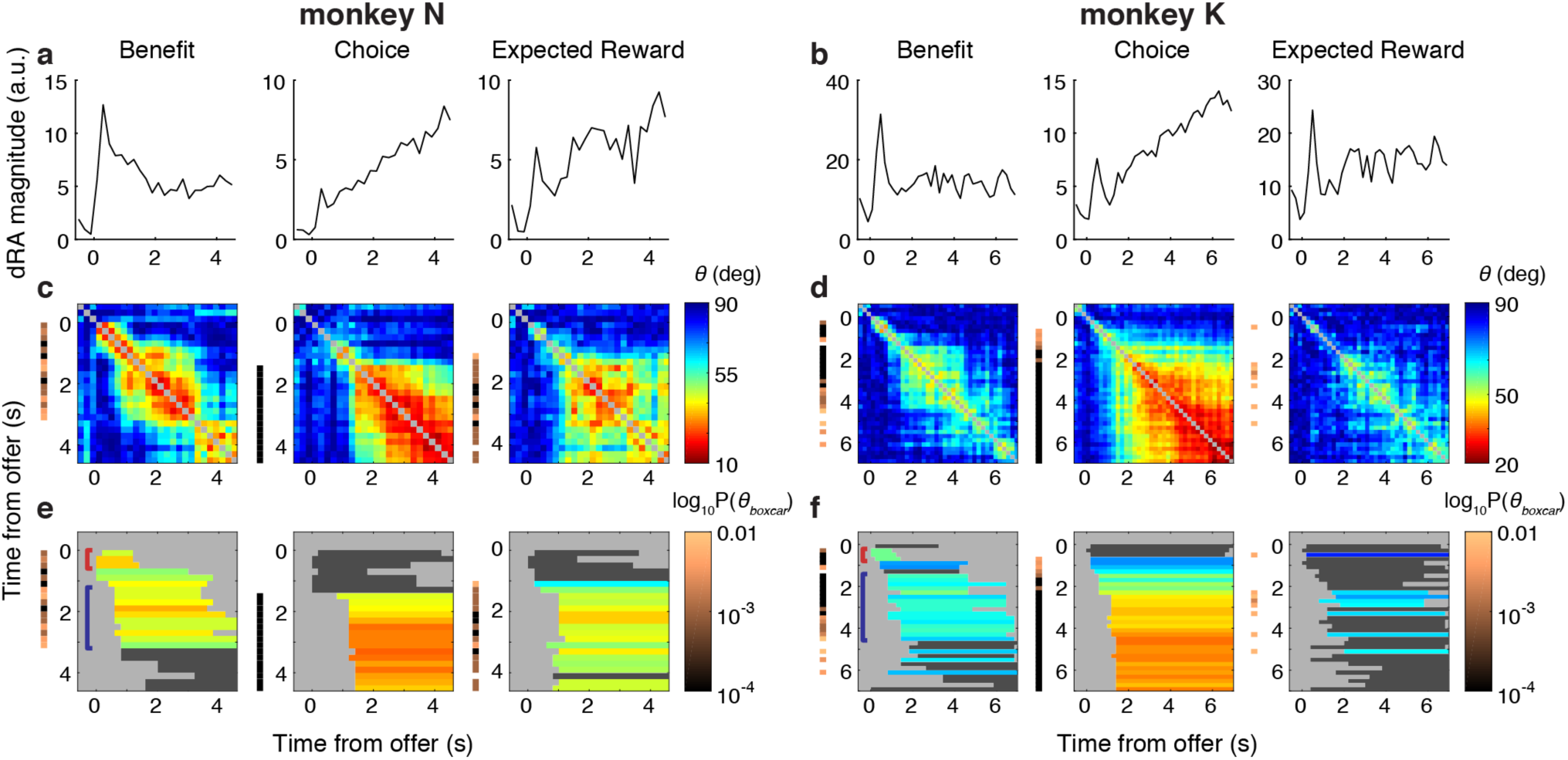
Stability of dynamic representations. For monkeys N (a,c,e) and K (b,d,f), features of the dynamic low-dimensional representations (dRAs) of benefit, choice, and expected reward are shown (left, middle, and right columns, respectively, for a given animal). **(a**,**b)** The vector magnitude of each dRA is shown as a function of time from the onset of the offer period and refers to the extent of population encoding at a given time, which was useful for interpretation: e.g., high similarity between weak representations may be less meaningful than when between robust, higher magnitude dRAs. **(c**,**d)** Angle *θ* in degrees between pairs of dRAs for the same variable is given by the color of each pixel (referencing right-hand color scale in (c,d)), while the times of the dRA pair (relative to the onset of the offer period) are given by the pixel’s row and column positions. Smaller angles (warmer colors) correspond to greater similarity between representations. The diagonal compares identical dRAs (*θ* = 0°) and is colored gray. **(e**,**f)** A boxcar function was fit to each row of angles in (c,d), with the boxcar height (*θ*_boxcar_) indicating the average similarity. The period of non-zero boxcar height is shown by a colored or dark gray horizontal bar in the corresponding row of (e,f). For periods of significant similarity (i.e., p(*θ*_boxcar_) < 0.01), bar color indicates boxcar height, referencing the right-hand color scale in (c,d), whereas non-significant periods are shaded dark gray. (For visual consistency, boxcar height was inverted such that higher boxcars, i.e., greater similarity, corresponded to smaller angles.) The thin vertical bands to the left of each panel in (c-f) show the log_10_ probability of observing *θ*_boxcar_ for the corresponding row, with colors referencing the right-hand color scale in (e,f); no color is shown when p(*θ*_boxcar_) ≥ 0.01. Colored brackets in left-most panel highlight specific periods referenced in main text. (Note: The boxcar fits and p-values captured periods of contiguous similarity. For a more granular account of observing the pairwise angles *θ*_*ij*_ by chance, see Supplementary Figure 20.)

**Figure 6.**
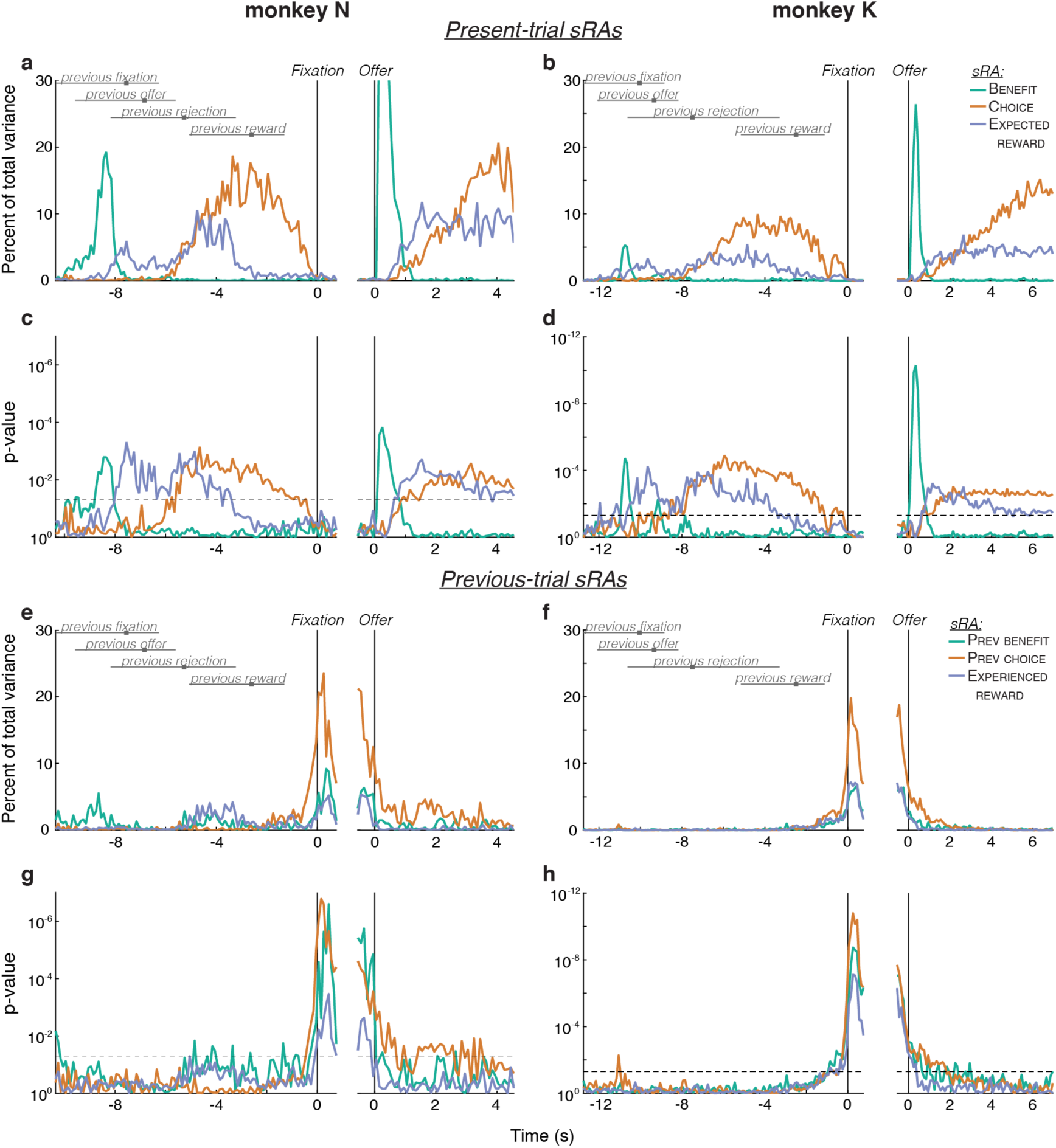
Cross-trial representations of task-relevant variables. **(a**,**b)** The relevant signal variance (RSV) for the present-trial sRAs of BENEFIT (green), CHOICE (orange), and EXPECTED REWARD (blue) was computed with respect to the previous-trial (left panels) or present-trial (right panels) conditions and plotted as a function of time from fixation (left panels) or offer onset (right panels) on the present trial for monkeys N (a) and K (b). Peak RSV for BENEFIT (a, right panel) is not shown so as to conserve the vertical scale across axes. Horizontal gray bars and solid gray squares indicate the 2.5-to-97.5 percentile range and median, respectively, of previous trial events, as labeled. RSV during the previous trial epoch (left panels) was smaller than during the present-trial epoch (right panels) likely due to the wide temporal variability between trials—reward delivery of 0 ∼ 3 s depending on reward number and ITI of 1 ∼ 2 s depending on when animal initiated fixation—leading to temporal “smearing” of otherwise robust, temporally aligned responses. **(c**,**d)** Log_10_ probability of data in (a) and (b), respectively, is plotted. Horizontal dashed line corresponds to p = 0.05. Note: Offer-aligned axes (right panels) of a-d recapitulate Figure 4c-f and are provided for reference. **(e**,**f)** The RSV for the previous-trial sRAs of PREVIOUS BENEFIT (green), PREVIOUS CHOICE (orange), and EXPERIENCED REWARD (blue) was computed with respect to the previous trial conditions for *both* temporal alignments (unlike for a,b, where offer-aligned RSV was with respect to present-trial conditions). Other conventions as in (a,b). **(g**,**h)** Log_10_ probability of data in (e) and (f), respectively, is plotted. Horizontal dashed line corresponds to p = 0.05.

### Optimal Targeted Dimensionality Reduction

To discover low-dimensional representations of the task-relevant variables, we developed *optimal targeted dimensionality reduction* (oTDR), an extension of the earlier TDR technique^19^. The oTDR method discovered linear combinations of neurons, or low-dimensional representations, that linearly encoded the task-relevant variables, i.e., were *targeted* to variables identified *a priori* as relevant to the behavioral task. We referred to the low-dimensional representations as *regression axes* (RAs), as they took the form of linear dimensions in the high-dimensional neural space and were defined by the coefficients for individual units from regression analysis. We referred to the variable for which a given RA was defined as *on-target*, while referring the other variables as *off-target*.

As an intuition, it is useful to consider our core assumption that for a given trial *r* and time *t*, each task-relevant variable *k* was represented linearly by a single dimension *β*_*k*_ as derived from the individual-unit, single-trial regression model:

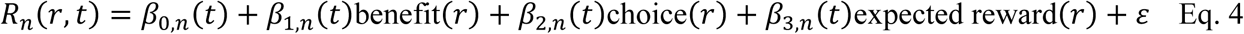

where *R*_*n*_(*r, t*) was the firing rate of unit *n, ε* was independent Gaussian noise, and the predictors were the task-relevant variables: benefit (encoded as {0, 1, 2, 4, 8}), choice (encoded as {0, 1} for rejects and accepts, respectively), and expected reward (given by benefit * choice). By compiling the regression coefficients across units, the vector *β*_*k*_(*t*) defined an *N*-dimensional vector, or RA, in the high-dimensional neural space. The projection of the population response onto this vector corresponded to the “activity” of the RA, as could be read out by a downstream circuit. Because this vector was defined by coefficients derived from linear regression, the variance across conditions in the RA activity was linearly proportional to variable *k*.

In practice, we operated on the trial-average, mean-subtracted, z-normalized population response captured in tensor 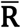 (see above). Likewise, we referred to the *K* = 3 task-relevant variables, or predictors, as P_*k*_, where P_1_ = benefit, P_2_ = choice, and P_3_ = expected reward. When computing previous-trial representations, P was replaced with the previous-trial variables {previous benefit, previous choice, experienced reward}, which referred to the condition in the previous trial, but were otherwise identical to their present-trial counterparts. All variables were scaled to the range [0, 1].

We applied oTDR for two, related goals: 1) to compute a single set of *static* RAs (sRAs) that represented the task-relevant variables across all times in the trial; and 2) to measure how the representations changed across the trial, as described by a set of *dynamic* RAs (dRAs) computed independently at each time in the trial.

#### Static low-dimensional representations

In the first application of oTDR, we sought a single set of sRAs to serve across all times in the trial. We reasoned that a static set of dimensions would both provide a more compact visualization of the data and suggest a means by which downstream neurons could read out distinct task-relevant representations via a static set of weights. Of course, if the dimensions representing the task-relevant variables changed markedly over the trial, then a set of static dimensions would fail to capture some portion of the available signal, which we quantified explicitly (see Supplementary Figure 16).

Toward discovering a static set of RAs, the model given by Eq. 4 posed several limitations, which we addressed via the following modifications. First, Eq. 4 solved for a given task-relevant representation *β*_*k*_ at each time *t*, generating multiple, time-varying RAs. This conflicted with our goal of a static representation, i.e., a single dimension that would represent variable *k* across multiple times. Therefore, we summarized the time-varying firing rate of Eq. 4 using the mean response 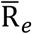 (dimensions *N* x *C*) taken over the subset of time bins 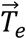. While this potentially limited the generalizability of the resulting representation, we reasoned that the sRA should capture the available signal during the epoch when the variable was most behaviorally relevant (which defined 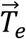), and we would assess generalizability subsequently as an empirical question.

In the present study, benefit information was relevant during the first 0.5 s after the offer, during which the animal encoded the incoming sensory information. After the offer period, the valuation and decision process proceeded without external information and thus depended on internal representations of the task variables (namely choice and expected reward), which we computed during the post-offer, “work” period. In addition to these *a priori* hypotheses, our selection of the temporal epochs was supported by the dRA analysis (Figure 5), which showed stable representations of benefit during the offer period and of choice and expected reward during the post-offer period.

Following the above rationale, we segregated the neural data into two epochs defined *a priori* by the sets of time bins 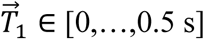 and 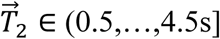 (monkey N) or (0.5,…,7s] (monkey K), and, within an epoch, averaged across the time bins to produce the *N* x *C* response matrices 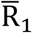 and 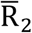, respectively. (Note the subscript and lack of boldface distinguished these response matrices from the time-varying responses given by the tensor 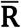, as defined above.) We discovered the sRA for benefit within 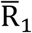 and discovered the sRAs for choice and expected reward within 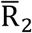. (As discussed below, the method supports an arbitrary number of epochs and the representation(s) of a given variable can be computed in one or more epochs.)

In separate analyses, we tested alternative hypotheses about the relevant epochs in which the variables were represented by varying the duration of 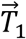 from 0.5 to 2 s (and shortening 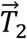 accordingly). We also tested the case of having no *a priori* hypothesis by discovering all three sRAs within a single epoch spanning the entire trial. In all cases, the results did not change qualitatively (Supplementary Figure 13).

Note that by discovering sRAs for choice and expected reward simultaneously in the same epoch, the representations competed to explain shared variance. This competitive process was critical when regressing the same neural responses (i.e., 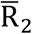) onto partially correlated variables, as was the case for choice and expected reward. The sRA for benefit was discovered in a distinct set of neural responses (i.e., 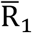) and did not compete for shared variance with the other representations. However, because we required all sRAs to be orthogonal (see below), the discovery of any sRA depended on the other sRAs, regardless of the temporal epoch in which it was discovered.

When discovering the previous-trial representations, we defined a single temporal epoch 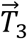 that included the 0.5 s prior to fixation on trial *r*, a time when the animal was under behavioral control but had not yet experienced the offer on trial *r*. Unlike for the present-trial sRAs, we discovered *all three* previous-trial sRAs (i.e., representations of the task-relevant variables in trial *r*-1) in the corresponding responses 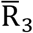. All other steps were identical to those in discovering the present-trial sRAs.

Second, the core regression assumption (Eq. 4) operated at the single-trial level and thus had the advantage of weighting each condition by the corresponding number of observations (i.e., trials), which differed systematically across conditions (e.g., animals rarely accepted 1-reward offers and usually accepted 8-reward offers). Therefore and conveniently, conditions for which our estimate of the true, long-run neural response was more reliable (i.e., we had more observations) exerted greater influence on the regression coefficients. However, the trial-average responses 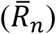, where were needed to combine responses across serially recorded units, did not incorporate the number of observations per condition. Thus, the reliability of the estimates in 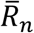 differed across conditions. To control for variable trial counts across conditions, we found that the single-trial model (Eq. 4) was equivalent to minimizing the Mahalanobis distance of the trial-averaged problem scaled by the square root of trial count (Appendix 1), and thus applied this correction when estimating the coefficients (see matrix M in Eq. 5.).

Third, Eq. 4 placed no constraints on the relationship between the sRAs. However, we would ultimately project the neural responses onto each sRA. For the projections to reflect independent portions of the total neural variance—and, per the goal of oTDR, reflect independent readouts of the task-relevant variables—we required orthogonality between the sRAs. As such, we constrained the regression such that all pairs of *β*_*k*_ were orthogonal^31^. We removed the orthogonality constraint when characterizing the intrinsic representation of a given variable or the relationships between representations (e.g., Figure 3). The constant term *β*_0_ was never included in the orthogonality constraint.

In summary, oTDR discovered static, low-dimensional representations (i.e., sRAs) of the task-relevant variables with the following assumptions:

1. At the level of the single trial, the neural response linearly represented task-relevant variable *k* as a single dimension *β*_*k*_ according to the individual-unit model:

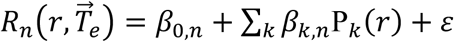

where 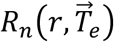 was the mean firing rate of unit *n* on trial *r* over the temporal epoch *e* given by set of time bins 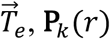, **P**_*k*_(*r*) was the value of variable *k* on trial *r*, subscript *k* indexed an arbitrary subset of task-relevant variables assigned to epoch *e*, and *ε* was independent Gaussian noise.
2. Reliability of the trial-average response 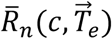 of unit *n* over all trials of condition *c* and time bins 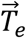 depended on the number of trials observed for condition *c*.
3. (Optional) Representations *β*_*k*_ were orthogonal across all *K* variables and epochs *e*, i.e., *β*_1_ ⊥ … ⊥ *β*_*K*_.

We incorporated all model assumptions into a single objective function, solved using established optimization tools^31,32^:

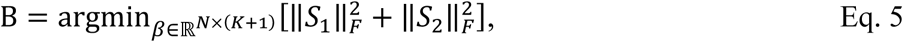

where

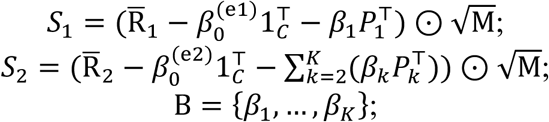

such that *β*_1_ ⊥ … ⊥ *β*_*K*_ (orthogonality assumption); where ‖·‖_F_ was the Frobenius norm; ⊙ indicated element-wise matrix multiplication; 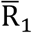 and 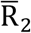 were *N* x *C* matrices specifying the average neural response of unit *n* in condition *c* over the first and second temporal epochs e1 and e2 (given by time bins 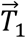 and 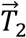), respectively; M was an *N* x *C* matrix specifying the number of trials observed for each unit *n* and condition *c*; 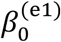 and 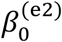 were *N* × 1 vectors specifying the constant term for the first and second temporal epochs, respectively; 1_C_ was a *C* × 1 vector of ones; *β*_*k*_ was an *N* × 1 vector specifying the regression coefficient of each unit *n* as pertaining to the variable *k*; and *P*_*k*_ was a *C* × 1 vector specifying the values of variable *k* for condition *c*, where *k* ∈ {1, 2, 3} pertained to the variables benefit, choice, and expected reward, respectively. (Note that we solved for the benefit coefficient, *β*_1_, exclusively in the first temporal epoch and solved for the choice, *β*_2_, and expected reward, *β*_0_, coefficients exclusively in the second temporal epoch, and thus it was not necessary to further specify the epoch for these coefficients.) The matrix B was of dimensionality *N* x *K*, and the columns contained the vectors *β*_1:*K*_, while the vectors of constant terms, 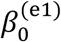 and 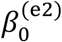, were not stored. The columns of B were then normalized to unit vectors, resulting in the final *K* sRAs.

#### Generalized form of the objective function

We designed oTDR to be a general-purpose algorithm for discovering low-dimensional representations of an arbitrary number of *K* variables and *E* epochs, while computing dimensions for each variable in one or more epochs and assuming orthogonalization between any subset of dimensions. For simplicity, we stated the objective function above (Eq. 5) in its narrow form as applied to the current dataset. Here we restate the objective in a generalizable form that can be applied to any dataset.

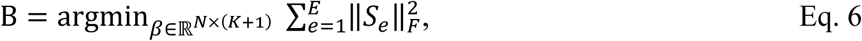

where

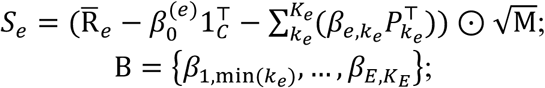

such that 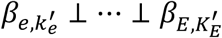 (orthogonality assumption) & 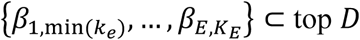 principal components of 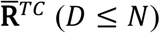; where 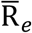 was an *N* x *C* matrix specifying the average neural response of unit *n* in condition *c* in temporal epoch *e* (given by vector of time bins 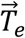); 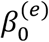 was an *N* × 1 vector specifying the constant term for epoch *e*; *k*_*e*_ ∈ {1, …, *K*_*e*_} referred to task-relevant variable *k*_*e*_ = *k* under the assumption that *k*_*e*_ was represented in epoch *e*; 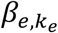 was an *N* × 1 vector specifying the regression coefficient of each unit *n* as pertaining to the variable *k*_*e*_ = *k* computed in epoch *e*; and 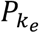 was a *C* × 1 vector specifying the values of variable *k*_*e*_ = *k* for condition *c*. The representation(s) of the variables 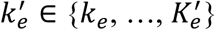 across all epochs *e* were assumed to be orthogonal (i.e., orthogonalization was applied to all of the selected representations regardless of epoch). Note that, for a given epoch, representations for only a subset of variables may be computed (i.e., *K*_*e*_ ≤ *K*) and, across epochs, only a subset of representations may be orthogonalized (i.e.,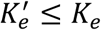). The 3-D tensor 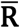 (with data spanning the entire trial) was transformed to the 2-D matrix 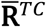 (dimensions *N* x *TC*) for which the principal components were computed. The columns of the matrix B contained the vectors 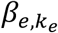 and was of dimensionality 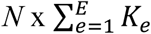, where *K*_*e*_ was the number of task variables computed in epoch *e*. The vectors of constant terms, 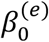, were not stored. The columns of B were normalized to unit vectors, resulting in the final sRAs. The remaining terms and conventions are defined above.

In Supplementary Figure 13e,f, we applied the generalized objective function to the current dataset to show how representations of a single task-relevant variable can be computed in multiple temporal epochs and how orthogonalization can be applied to a subset of representations. We also discuss the rationale for these assumptions.

As stated above, the objective function could accommodate an additional de-noising assumption such that the task-relevant representations were limited to the high-variance subspace, i.e., space spanned by the top *D* principal components (PCs). We did not apply the de-noising assumption in the present study, in part because the noise-reducing effect of averaging over multiple time bins 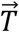 obviated the need for additional noise reduction. More importantly, limiting the data to the high-variance subspace would compromise the subsequent hypothesis testing. Specifically, as described below, we tested the significance of a given representation (i.e., sRA) by comparing it to random vectors biased to the space occupied by the data. By limiting the data to the high-variance subspace, the random vectors would also be limited to this high-variance subspace—a subspace much smaller than that spanned by the original data. As a result, the random vectors would underestimate the full space of possible representations, and our estimates of the probability of obtaining a given sRA by chance, as computed from the random vectors, would be exaggerated.

#### Comparison to prior technique

The present oTDR technique extended the previous TDR method^19^. Both techniques were “targeted” in that variables of interest were defined *a priori* and then dimensions representing those variables were discovered in the high-dimensional neural data. However, the previous technique used separate, ad-hoc algorithms to apply each assumption serially: computing a set of linear representations *β*_*k*_ of variable *k* via Eq. 4, projecting *β*_*k*_ onto the top PCs (also see ref^33^), and then orthogonalizing the set of *β*_*k*_ using a greedy algorithm. Each step distorted *β*_*k*_ from the original vectors, and thus the final vectors were no longer necessarily as close to linear representations of the targeted variables as possible given the model assumptions. In addition, the prior technique relied on solving for *β*_*k*_ at a single moment in time, which compromised the generality of the representation across all relevant times. In contrast, by incorporating all assumptions into a single objective function (Eq. 5), oTDR satisfied the model assumptions simultaneously and thereby discovered targeted dimensions that were optimal given all model assumptions.

Of note, as for oTDR, the prior technique implicitly weighted conditions by the number of observations (assumption 2) through use of the single-trial model (Eq. 4). However, use of this model complicated simultaneous application of the orthogonalization and de-noising steps, since application of these steps would require the objective function to solve for each unit serially and transition back-and-forth between single-trial and trial-average data. Our finding that the single-trial model could be recast as a scaled trial-average problem facilitated the simultaneous application of all assumptions.

#### Dynamic low-dimensional representations

Thus far, we applied oTDR to discover static representations, or sRAs, of the task-relevant variables. In our second application, we measured how the representations changed across the trial. To this end, we discovered a time series of dynamic representations, or dRAs, discovered independently in each time bin. The dynamic and static analyses shared the basic assumptions outlined in the single-trial model (Eq. 4), but the methods differed slightly. In particular, the dynamic analysis operated on shorter time bins, which made the regression coefficients more susceptible to random variation in firing rate. As such, we applied several additional steps to reduce the impact of random variation.

First, prior to solving the regression, we noised-reduced the neural data by eliminating the low-variance dimensions, which contributed less to representations of the task-relevant variables (as confirmed in Supplementary Figure 19). To identify the low-variance dimensions, we transformed the 3-D tensor 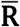 (with data spanning the entire trial) to the 2-D matrix 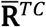 (dimensions *N* x *TC*) and performed principal components analysis (PCA) on 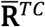. (Note, by transforming 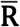 to 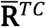, the resulting PCs explained variance related *either* to time *or* task condition.) We projected the trial-average, mean-subtracted response 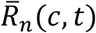 onto the top *D* PCs, as ordered from those explaining the most to least variance, generating the noise-reduced responses *Ř*_*n*_(*c, t*). We selected *D* as the approximate inflection point of the log variance explained as a function of PC. The subspace spanned by the top PCs (*D* = 8 or 20) explained 45% or 32% of total variance across time and conditions for monkey N or K, respectively.

Second, we averaged adjacent 100 ms time bins into single, non-overlapping 200 ms time bins that did not straddle *t* = 0. The variable “*t*” in the dynamic analyses refers to these wider, 200 ms time bins.

Third, we applied *L*_2_ regularization (i.e., ridge regression) to mitigate random variation in the coefficients *β*_*k*.*n*_(*t*). The regularization penalized large values of *β*_*k,n*_(*t*) and assumed the contribution of a given unit *n* was distributed across variables *k.* This assumption was supported by the observed mixed selectivity of individual units, i.e., lack of clustering of sRA coefficients along the horizontal and vertical meridians (Figure 3). Of note, we did not apply *L*_2_ regularization when discovering the sRAs (Eq. 5) for two reasons. First, the noise-reducing effect of regularization was less necessary in the static analysis given the more precise measurements of the neural response, as discussed. Second, in discovering the special case of non-orthogonalized sRAs (e.g., Figure 3), we were interested in accurately measuring the absolute relationships between task-relevant representations, which would be artificially decreased by assuming *a priori* (via regularization) that coefficients were distributed across variables. In contrast, we used the dRAs primarily to understand the change in relationships between representations over time (relative to synthetic data to which regularization was also applied; see below), and thus the absolute relationships were less critical.

We tested the effects of PCA-based noise-reduction, time bin width (100, 200, or 500 ms), and regularization in separate analyses (Supplementary Figure 24).

Finally, unlike the sRAs, we did not orthogonalize the set of dRAs because we were interested in the relationship between the representations—either of different variables at the same time, or of the same variable at different times—and orthogonalization would have distorted these relationships.

In solving the static regression model (Eq. 5), numerical matrix optimization was required to include the orthogonality assumption. However, because orthogonalization was not included in the dynamic analysis, the model for the dRAs had a closed-form solution that could be solved independently for each unit:

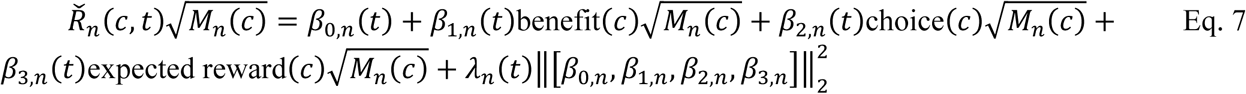

where *Ř*_*n*_(*c, t*) was the noise-reduced, trial-average response of unit *n* for condition *c* at time *t, M*_*n*_(*c*) was the number of trials observed per condition, *λ*_*n*_(*t*) was a scalar parameter governing the impact of the regularization term, and the regularization term, ‖·‖_2_, was the Euclidian norm of the vector of coefficients *β*_*k,n*_ across variables *k* and the constant term *β*_0,*n*_. We found *λ*_*n*_(*t*) empirically and independently for each unit and time bin via leave-one-out cross-validation on conditions and selecting the value of *λ* that minimized the mean squared error for Eq. 7 on the left-out test condition. For certain units and time bins, the error was minimized by *λ* = ∞, implying excessive unexplained variance; in these cases, we set *β*_*k,n*_(*t*) = 0 for all *k*. We compiled coefficients *β*_*k,n*_(*t*) across units to generate *N*-dimensional vectors for each variable *k* that we then normalized to unit vectors, referred to as the “dRA(*t*) for [variable *k*]” in the main text.

### Metrics of regression axes

#### Projection of neural response onto static regression axes

Each sRA defined a linear combination of neurons that represented the targeted task-relevant variable. To read out these representations, we projected the neural responses onto the set of sRAs:

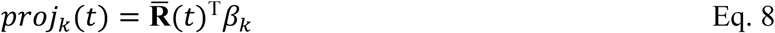

where 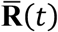 was an *N* x *C* matrix of the population response extracted from tensor 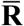 at time *t*, and *β*_*k*_ was the *N*-dimensional sRA corresponding to variable *k*. The *C*-dimensional vector *proj*_*k*_(*t*) gave the “activity” of sRA_*k*_ at time *t* for each of the *C* conditions.

#### Note on limiting projection analysis to the static regression axes

As discussed, a primary aim of the oTDR static analysis was to separate, or de-mix, the population encoding of the task-relevant variables into independent representations. Because the sRAs were orthogonal, the projection onto a given sRA was the optimal readout of the population response (given the model assumptions) as related linearly to the targeted variable of interest and minimally related to the other task-relevant variables. However, if the RAs were non-orthogonal, as was the case for the dRAs (see above), then the resulting projections necessarily would contain representations of multiple task-relevant variables. While these mixed projections may separate the task-relevant variables more than the mixed selectivity at the individual-unit level, they would not offer a maximally independent readout. Therefore, projections onto the dRAs were of limited utility for interpreting the population activity or for positing how downstream circuits may read out the population activity. For this reason, we did not project the population response onto the dRAs, and consequently did not perform the subsequent variance-based analyses (see below) that depended on these projections.

#### Variance explained by static regression axes

We measured the *variance explained* (V_*k*_) by sRA_*k*_ at time *t* with respect to variable *k* as the variance of *proj*_*k*_(*t*) across conditions normalized by the cross-condition variance summed over all dimensions (i.e. units), which we expressed as a percentage:

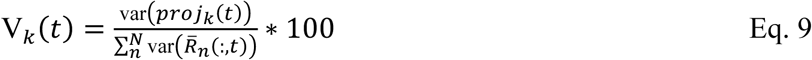

where 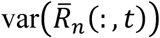 was the variance of neural response across conditions for unit *n* at time *t.*

#### Relevant and irrelevant signal variance

Though each sRA was designed to represent a specific task-relevant variable, some of the variance explained by a given sRA may not have been related to the targeted variable. Moreover, some of the unrelated variance may have been related to an off-target variable. We were interested in measuring the variance explained by a given sRA_*k*_ (designed to target variable *k*) that was related or unrelated to variable *q*, and so developed metrics for *relevant* or *irrelevant signal variance* (RSV or ISV, respectively):

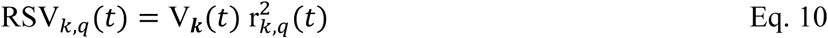

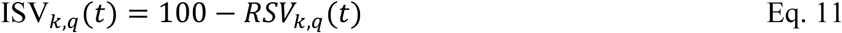

where V_*k*_(*t*) is the variance explained by sRA_*k*_ at time *t* and 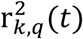 is the squared Pearson’s correlation coefficient (0 ≤ r^2^ ≤ 1) between *proj*_*k*_(*t*) and the condition-matched values of variable *q* (note, by convention, the subscript *k* in 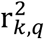 did not refer to variable *k* directly, but rather to the projection onto sRA_*k*_). Thus RSV and ISV were in units of variance, lower-bounded by 0, and together summed to V_*k*_(*t*). By convention, when referring to the on-target variable (i.e., *q* = *k*), we simplified the subscripted indexing: RSV_*k,q*=*k*_ = RSV_*k*_ and ISV_*k,q*=*k*_ = ISV_*k*_.

When we computed RSV and ISV for the on-target variable (i.e., *q* = *k*), the metrics provided a useful account of how well the sRA was sensitive and specific, respectively, to the variable of interest. However, when we computed RSV or ISV for *off-target* variables (i.e., *q* ≠ *k*), the term could misattribute on-target variance as off-target variance when variables *k* and *q* were correlated (e.g., the values of benefit [0, 1, 2, 4, 8, 1, 2, 4, 8] and expected reward [0, 0, 0, 0, 0, 1, 2, 4, 8] were correlated, *r* = 0.54).

To consider the effect of correlated variables, let *a* and *b* be on- and off-target variables, respectively, moderately correlated according to the Pearson’s correlation coefficient r_*a,b*_. Let *p* be the projection of the neural population response onto sRA_*a*_ designed to detect variance related to *a*. To simplify the formulation, assume *a, b*, and *p* are mean-centered, though this is not essential to the argument. Let *a* and *p* be correlated by a relative large coefficient r_*p,a*_.

Consequently, we would compute a relatively high on-target RSV_*a*_. However, given r_*a,b*_, we would also expect a relatively high correlation r_*p,b*_ and thus a high off-target RSV_*a,b*_, giving a false impression that RSV_*a*_ is non-specific to variable *a* and explains substantial variance related to variable *b.* In the extreme case, when r_*a,b*_ = 1, then r_*p,b*_ = r_*p,a*_, and thus RSV_*a,b*_ = RSV_*a*_.

To control for the correlation r_*a,b*_ between variables, we employed a method known as semi-partial correlation^34^ to isolate the relationship between *b* and *p* that was not explained by r_*a,b*_. We replaced the Pearson’s correlation coefficient r_*p,b*_ with the semi-partial correlation coefficient *ρ*_*p,b*|*a*_, as given by

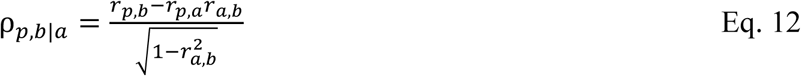

Therefore, to isolate the portion of variance related to off-target variables, while controlling for the correlation between on- and off-target variables, we computed off-target RSV as

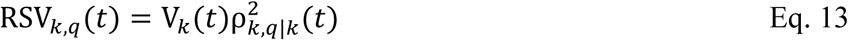

where 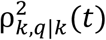 was the squared semi-partial correlation coefficient (Eq. 12) between *proj*_*k*_(*t*) and the component of off-target variable *q* not explained by on-target variable *k*. We continued to computed RSV for on-target variables (i.e., *q = k*) per Eq. 10. Thus,

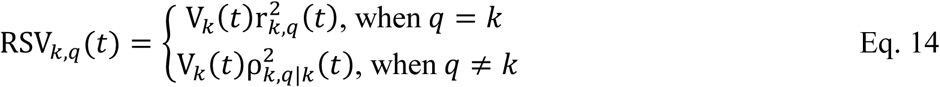

Regarding ISV, on-target ISV_*k*_ indicated the amount of variance explained by sRA_*k*_ that was not correlated with the on-target variable *k* and thus was available for correlation with the off-target variables. The concept of off-target ISV was of limited utility, as one expected for the variance explained by a given sRA to be *un*related to the off-target variables. Indeed, high on-target RSV indicated high off-target ISV. Therefore, we only computed ISV for the on-target variables.

#### RSV and ISV for arbitrary axes

The metrics RSV and ISV were not limited to sRAs computed by oTDR and could be computed for any arbitrary axis. However, for dimensionality reduction techniques agnostic to the variables of interest (e.g., PCA), the notion of on- and off-target variables was not well-defined. In the case of PCA (Supplementary Figure 17), the dimensions (i.e., PC_*d*_) corresponded to vectors in the *N*-dimensional space that explained the greatest cross-condition and temporal variance, from *d* = 1 to *N*. As with the sRAs, we projected the neural population response onto each PC_*d*_ and measured the variance explained (Eq. 9). For a given task-relevant variable *k*, we defined the on-target PC_*d′*_ as that which explained the greatest cumulative RSV_*d,k*_ with respect to variable *k* (as per Eq. 10) across all time bins, and likewise defined *d′* as the number of the corresponding PC. Subsequently, we recomputed off-target RSV_*d,k*_ (as per Eq. 13) for the remaining PCs (i.e., when *d* ≠ *d′*), while maintaining on-target RSV_*d*′,*k*_ per Eq. 10 when *d* = *d′*.

#### Angle between regression axes

To measure the similarity between either the static or dynamic representations, we computed the pairwise angle in degrees between RAs *β*_*i*_ and *β*_*j*_:

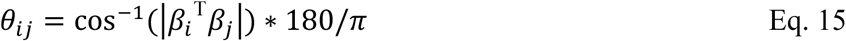

Note that in this general formulation, *β*_*i*_ and *β*_*j*_ were unit vectors and could refer to several types of an RA pair: sRAs for variables *i* and *j*, dRAs for the same variable at times *i* and *j*, or dRAs for different variables *i* and *j* at the same or different times. Angles of 0° indicated the representations were highly similar, whereas angles of 90° indicated the representations were maximally unrelated (i.e., orthogonal). We took the absolute magnitude of the dot product so as treat angles equidistant from 90° as equivalent (e.g., angles of 0° and 180° were both coded as *θ* = 0°), and thus refer to *θ*_*ij*_ as the *folded* angle. In so doing, we emphasized the absolute similarity of the RA pair (e.g., angles of 0° and 180° both indicated that the same units contributed to the same absolute amount to the pairs of RAs). However, this construction of *θ*_*ij*_ was insensitive to differences in sign of representation, as when the neuronal contributions to *β*_*i*_ and *β*_*j*_ remain similar in absolute strength but reverse sign, such as exhibited by the example unit in Figure 2d. Therefore, exclusively to observe changes in sign, we constructed the *unfolded* angle *θ*′_*ij*_ as:

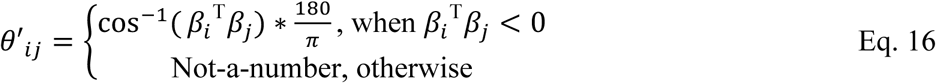

and *θ*′_*ij*_ was therefore limited to the range [90, 180°].

#### Alignment index

In addition to comparing the similarity between 1-dimenional RAs via the angle analysis above, we were interested in assessing the overlap between pairs of subspaces spanned by sets of RAs or other basis vectors. To measure the overlap between subspaces U_1_ and U_2_, we used a custom metric, termed the alignment index *A*, adapted from a recent report^35^:

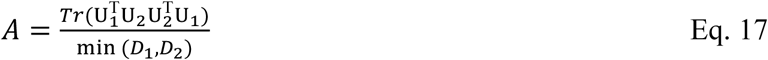

where U_1_ and U_2_ were orthogonal matrices of dimensions *N* x *D*_1_ and *N* x *D*_2_, respectively, with *D* basis vectors arranged in columns, and *Tr*(.) was the matrix trace. The numerator measured the amount of overlap between U_1_ and U_2_, and the denominator normalized the index by the minimum dimensionality of the subspaces (i.e., the maximum possible overlap). Thus, the alignment index ranged from 0 (orthogonal) to 1 (completely aligned) and was invariant to the order of U_1_ and U_2_.

### Hypothesis testing

A major goal of the present study was to develop and apply generalizable, unbiased tools for significance testing of low-dimensional representations of high-dimensional data. We developed two statistical approaches to contextualize the low-dimensional features by generating control datasets of either 1) random dimensions that captured the inter-neuronal correlational structure (i.e., dimensionality) of the data (Random dimensions in neural space), or 2) random synthetic responses that captured both the inter-neuronal and temporal correlational structure (i.e., temporal smoothness) of the data (Neural population control data).

#### Random dimensions in neural space

To model the correlations between neurons (i.e., dimensionality), we generated random dimensions that reflected the high-dimensional space occupied by the neural data^35^. That is, the density of random dimensions in the high-dimensional space was proportional to the frequency that the neural population occupied a given region of the space.

To provide an intuition for the impact of dimensionality, consider a population of two neurons that tended to fire together. The dimension reflecting co-activation would be more likely to contain data than the dimension reflecting exclusive activation of one neuron or the other. This asymmetry should be reflected in the set of random dimensions intended to model the chance probability of a given dimension (or its properties) given the data. Alternatively, if the set of random vectors were evenly distributed throughout two-dimensional space (as in most analyses), then many random dimensions would be over-represented relative to how frequently the neural population actually occupied those dimensions, thereby overestimating the rarity of the commonly occupied dimensions and generating spuriously small p-values. Instead, by generating random dimensions reflecting the correlational structure between neurons, our estimate of the rarity of a given dimension was less biased and the resulting p-values were more conservative.

The details of the random dimension method are described elsewhere^35^. Briefly, we calculated the covariance matrix Σ of the neural responses 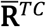 (dimensionality *N* x *TC*) across all times and conditions of the task (as was done for computing the PCs above). We then generated random dimension 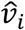aligned to the dimensionality of the neural data as:

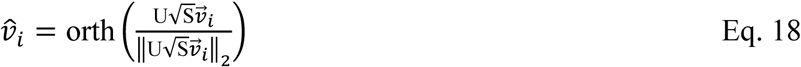

where the eigenvectors of Σ were in the columns of the *N* x *N* matrix U, and the corresponding eigenvalues were on the diagonal of the *N* x *N* diagonal matrix S; 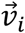 was a random *N*-dimensional vector with each element drawn independently from a normal distribution with zero mean and unity variance; and orth(·) returned the orthonormal basis of an input matrix. We repeated this procedure for *i* = 1 to 10,000, returning a set of random dimensions that had the specified inter-neuronal covariance structure Σ. For statistical analysis, we projected the neural data onto the set of random dimensions, generating 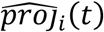 for each random dimension 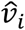. Using 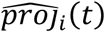, we computed identical metrics as when projecting the data onto a dimension of interest (e.g., when computing variance explained by an sRA). Across random dimensions, this produced a null distribution for a given metric from which we estimated the probability of obtaining a given value of that metric by chance given the inter-neuronal correlational structure of the data.

#### Note on null distribution of off-target RSV

In general, replicating a given metric based on the projection onto a random dimension (i.e., 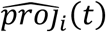) was straightforward. However, computing off-target 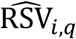 for random dimension *i* and off-target variable *q* required specification of the semi-partial correlation 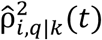 that isolated the portion of *q* that was independent of the Pearson’s correlation 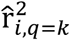 between the specific projection 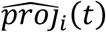 and the on-target variable *k*. Therefore, we first computed the on-target correlation 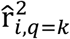 (see Eq. 10) for random dimension *i* and on-target variable *k*, which we then used to compute the semi-partial correlation 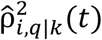 (via Eq. 12) and, in turn, off-target 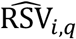 (via Eq. 13).

#### Note on null distribution of angles

We compared the similarity between two dimensions (i.e., representations) as the angle *θ* (Eq. 15; Supplementary Table 2-4). We generated the null distribution of angles 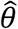 by computing the angle between all pairs of random dimensions 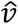.

#### Neural population control data

The random dimensions in the neural space captured the data’s inter-neuronal correlations (i.e., dimensionality), but not the temporal correlation, or smoothness, of firing rates at the level of individual units. While the data’s dimensionality restricted the extent to which a representation could change over time, the temporal smoothness was relevant to how quickly a low-dimensional representation changed over time. We sought to quantify the change expected from these low-level correlations—temporal and inter-neuronal—that was not attributable to the population’s encoding of the task-relevant variables. From this null expectation, we would then quantify the rarity, or significance, of a given representation’s consistency as observed in the data. However, traditional means to model correlational structure, such as shuffling procedures, can capture *either* the data’s dimensionality *or* temporal smoothness, but not both. Instead, we employed a new method, Neural population control^21^, to generate synthetic firing rate data that preserved both the dimensionality and temporal smoothness of the original data but were otherwise random. In brief, the method sampled firing rates at each time bin and for each surrogate neuron from a joint probability distribution that was maximally entropic except for the data’s temporal and inter-neuronal correlational structure, quantified by the covariance of the original data across times and neurons. Given *N* units in the original dataset, we generated 1000 surrogate datasets of *N* neurons each, with the original data’s correlational structure preserved within a given surrogate dataset (i.e., individual datasets were conditionally independent from one another). By computing identical metrics against the surrogate datasets as for the original data, we compared a given value observed in the original data to the distribution of surrogate values. Any population feature, such as stability of representation, that appeared in the surrogate datasets would be considered epiphenomenal, i.e., an expected byproduct of the data’s dimensionality and temporal smoothness. Whereas population features that rarely occurred in the surrogate datasets would be considered statistically significant.

### Separability and reliability of representations

The present study sought to identify the encoding of task-relevant variables at the level of individual units and separate these signals into distinct low-dimensional population representations. Our efforts therefore depended on whether the population representations were indeed separable, a concept for which we develop a definition here. Though prior studies have shared similar goals^19,20,36^, the concept of separability was not formally defined or tested.

A rigorous examination of separability is imperative when making claims about demixing high-dimensional representations since, as we outline below, noisy estimates of the individual representations can contribute to exaggerated estimates of the independence between representations, thus overestimating the very basis for de-mixing. Also at stake are claims of mixed selectivity, since a population of units selective exclusively for variable *A* can appear to also encode variable *B* when estimates of the representations of *A* or *B* are noisy, and thereby generate the spurious impression that selectivity is “mixed”.

In defining separability, let vectors 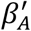 and 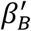 be the true *N*-dimensional linear representations (i.e., RAs) of task-relevant variables *A* and *B* defined for a population of *N* neurons. We will define 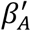 and 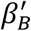 as *separable* when they contain independent information, i.e., when their correlation coefficient is less than unity, |*r*_*A*′ *B*′_ | < 1. To test for separability, we therefore define a null hypothesis that the representations are *not* separable, i.e., perfectly correlated:

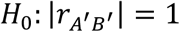

However, we do not have access to the true representations—only to the sample estimates *β*_*A*_ and *β*_*B*_, and their correlation |*r*_*AB*_|, as measured from the data. Following standard hypothesis testing procedure, we calculate the probability *P* that we obtained a particular value of |*r*_*AB*_| under the null hypothesis, i.e., *P*(|*r*_*AB*_| | *H*_0_). Here we develop a sampling distribution for |*r*_*AB*_| given *H*_0_.

The relationship between |*r*_*AB*_| and |*r*_*A*′ *B*′_| is given by Spearman^37^, who observed that |*r*_*AB*_| decreases as independent noise, *ε*_*A*_ and *ε*_*B*_, is added to the true representations, i.e.:

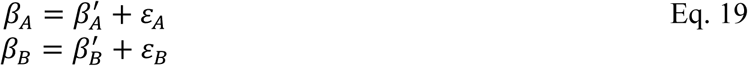

At the limit, the measured correlation between two perfectly correlated variables goes to zero as *ε* ≫ *β*′. We do not directly measure *ε*_*A*_ and *ε*_*B*_, but instead measure the effect of independent noise on the reliability of our measurements of *β*_*A*_ and *β*_*B*_. Our measure of reliability is based on the correlation, *r*_*AA*_ or *r*_*BB*_, between repeated samples of *β*_*A*_ or *β*_*B*_, which we simulate via a bootstrap procedure, detailed below. If reliability were perfect, then *r*_*AA*_ = 1 and *r*_*BB*_ = 1. Spearman offered the following correction for the attenuating effect of independent noise (i.e., imperfect reliability) on the correlation between two variables^37^:

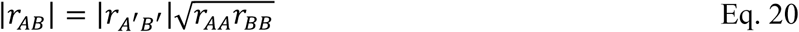

Under the null hypothesis (i.e., |*r*_*A*′ *B*′_| = 1) the above expression reduces to:

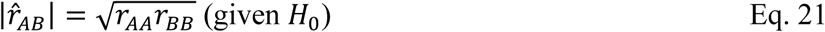

(Note use of the “hat” symbol to indicate the null-hypothetical value.)

In detail, the bootstrap procedure to estimate *r*_*AA*_ or *r*_*BB*_ was as follows. We generated *S* resampled datasets by randomly selecting *Q*_*n,c*_ trials with replacement from each condition *c* and unit *n* given *Q*_*n,c*_ original trials for that condition and unit, such that the number of trials per condition for a given unit was consistent across the original data and resampled datasets (*S* = 700; chosen for the maximum number of trials per unit). Note that we *z*-normalized the resampled data for a given unit via the single mean and standard deviation observed for that unit in the full dataset. This had the effect of decreasing our subsequent reliability measure and making our determination of separability more conservative. The alternative—*z*-normalizing independently within each resampled dataset—would minimize trial-to-trial response fluctuations and thereby overestimate reliability and separability.

Within each resampled dataset, we used to oTDR without orthogonalization to calculate the three sRAs for the task-relevant variables benefit, choice, and expected reward. Here we discuss the representation *β*_*A*_ (i.e., sRA) of variable *A*; an identical procedure was used for the other two variables. We computed all pairwise correlations between *β*_*Ai*_ and *β*_*A,j*_, where *i* and *j* were different resampled datasets (given *S* datasets, we calculated (*S*^2^ − *S*)/2 correlations). We compiled these correlations into the distribution ***r***_*AA*_ (shown in Supplementary Figure 8a,c).

To estimate 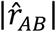 under the null hypothesis, we computed the distribution 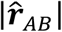 (shown in Supplementary Figure 8b,d) for each pair of variables *A* and *B* (given 3 variables, we computed 3 pairs) according to Eq. 21 using the just-compiled distributions of reliability ***r***_*AA*_ and ***r***_*BB*_:

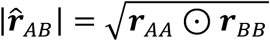

where ⊙ indicated element-wise multiplication.

Finally, we performed a 1-tailed *t*-test to determine the probability *P*(|*r*_*AB*_| | *H*_0_) of falsely rejecting the null hypothesis that the values in 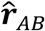 were from a distribution with mean |*r*_*AB*_| (as observed in the data) in favor of the alternative hypothesis that 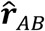 came from a distribution with greater mean (i.e., that the observed |*r*_*AB*_| was less than null-hypothetical value 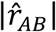. When *P* was sufficiently small, we concluded that the representations *β*_*A*_ and *β*_*B*_ were separable.

Our definition of separability serves as a useful test for whether two representations contain independent information about their respective variables and highlights the confounding influence of noisy representations (a feature in virtually all empirical datasets) on observing separability erroneously. Despite these advantages, we consider the following limitations. Because we used a bootstrap procedure (i.e., resampled datasets shared some trial-level data), the resulting reliability estimates (i.e., *r*_*AA*_ and *r*_*BB*_) were admittedly biased upward, thus making our final test of separability less conservative (via Eq. 21). In addition, because *β*_*A*_ and *β*_*B*_ were derived from the same set of neural responses and because the corresponding variables *A* and *B* may be correlated, we cannot guarantee that the respective noise components, *ε*_*A*_ and *ε*_*B*_, were themselves independent, an assumption of Eq. 20. If *ε*_*A*_ and *ε*_*B*_ were correlated, then the measured correlation |*r*_*AB*_| would increase (via Eq. 19) without a change in the null-hypothetical 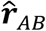 (which depends on *ε*_*A*_ and *ε*_*B*_ separately, not on their interaction), thus making our conclusions regarding separability *more* conservative.

In addition to the separability analysis, we used the resampled datasets to measure the reliability of each unit’s contribution to a given representation. As discussed above, the representation *β*_*A,s*_ of variable *A* was derived for each resampled dataset *s*, and each element of 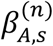 specified the contribution of unit *n* to the representation. We compiled the distribution 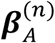 across the *S* datasets. The standard deviation of the distribution was proportional to the reliability of unit *n*’s contribution (shown by error bars in Figure 3a,c). Separately, via a two-tailed *z*-test, we computed the probability of falsely rejecting the null hypothesis that the distribution 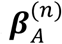 included the value of zero. When this p-value was sufficiently small, we concluded that unit *n* “significantly encoded” variable *A*. Subsequently, we tested whether the population’s selectivity was indeed “mixed” by whether the proportion of units significantly encoding multiple variables was greater than expected by chance (see Supplementary Table 1). This test controlled for the confounding effect of noisy estimates of 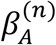 and 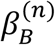 appearing as mixed selectivity for variables A and B, when in fact one or both estimates may have been indistinguishable from zero, and thus the true selectivity was for only one or neither variable.

## Results

### Subjective value increases smoothly with increasing benefit and fixed cost

In a novel cost-benefit task, two macaque monkeys, N and K, decided whether to trade sustained effort for juice reward. The number of juice drops offered was presented briefly as 0, 1, 2, 4, or 8 yellow icons and varied randomly across trials (Figure 1a). To accept an offer, the animal maintained visual fixation for a constant duration (work period) and then received the promised reward. To reject an offer, the animal averted its gaze and waited for the next trial. We analyzed 9,637 and 27,952 trials from 26 and 86 experimental sessions (monkeys N and K, respectively).

As expected, the likelihood of the animal accepting an offer increased smoothly with offer size (Figure 1b), which we confirmed in a logistic model fit to individual sessions (Supplementary Figure 1). To control for the inherent confound between the stimulus’s reward magnitude and the visual impact of more icons, we presented the 8-reward offer either as 8 yellow icons or as a single purple icon (singleton offer), to which the animals responded similarly (Figure 1b, open circles). Taken together, this suggested that animals computed the value of each offer based on its variable benefit (drops of juice) and fixed cost (effort).

### Mid-trial reevaluation of choice

A key task feature was the sustained effort required to accept an offer, allowing the animal to accept an offer initially, but continuously reevaluate its choice and, at times, change its mind, rejecting the offer mid-trial. We leveraged the timing of fixation breaks—measured as the rejection hazard rate (Figure 1c and Supplementary Figure 2)—to infer the dynamics of the underlying decision process. (Here we focus on the non-zero offers, which were qualitatively distinct from 0-reward offers; see Supplementary Figure 3.) For a given offer size, the hazard rate peaked within 1 – 1.5 s of the offer (*early phase*), suggesting most decisions occurred early in the trial, as the task structure incentivized.

However, a second phase of rejections occurred mid-trial—from ∼1 s after the offer until the final 1 ∼ 2 s of the work period (*mid-trial phase*)—with qualitatively distinct magnitude and dynamics from the earlier rejections. The progressively decreasing hazard rate was consistent with volitional rejections and not accidental fixation breaks, unlike the final phase (see below). These features suggested a separate decision process: on a small fraction of trials, the animal initially accepted the offer, but later changed its mind and rejected the offer mid-trial^38–40^. Several value-based variables likely evolved over the trial and may have contributed to these mid-trial reversals, such as an increase (decrease) in expected benefit (cost) as the time-to-reward shortened^41^. Importantly, these mid-trial rejections continued to depend on offer size, therefore requiring a memory of the offer (the stimulus was no longer visible) and motivating our search for a sustained neural representation of expected reward.

Finally, unlike the changing rates earlier in the trial, the hazard rate for most non-zero offers converged to a low and static level that did not depend on offer size for the final 1 ∼ 2 s of the work period (*late phase*). A constant hazard rate is a feature of a Poisson process in which events occur with constant probability per unit time^42^, consistent with accidental failures of fixation (i.e., “falling off the log”), and may partly account for the maximal accept rates of less than 100%.

### Heterogeneous encoding of task-relevant variables in individual OFC units

We analyzed the spiking activity of 68 and 342 units from OFC in monkeys N and K, respectively (see Methods and Supplementary Figure 4 for anatomy). Responses from single units and multi-units did not differ qualitatively and were analyzed collectively as “individual” units (see Supplementary Figure 32-34 for analysis of single units exclusively).

As expected, individual units responded robustly to the offer and were modulated by offer size, consistent with encoding of stimulus value. However, we observed broad heterogeneity, both in the dynamics—from rapid and transient (Figure 2a,c) to slow and sustained (b)—and sign of encoding—from positive (a,b) to negative (c; i.e., firing less for larger offers), or reversing during the trial (d; i.e., inverting sign of encoding mid-trial). For a minority of units, we observed non-monotonic encoding, i.e., firing most or least for intermediate offers (Supplementary Figure 5). Individual units encoded not only offer size, but also the animal’s choice (e), consistent with mixed selectivity for multiple variables. Conventional approaches for summarizing this heterogeneity (i.e., classifying responses into discrete categories and reporting the percentage of units per category and time bin^5,15^; see Introduction), obscured how the encoding strength, sign, and/or selectivity of an individual unit changed over the trial—dynamics which were clearly evident in individual responses (e.g., Figure 2).

### Discovering low-dimensional representations of task-relevant variables

Dynamic encoding and mixed selectivity presented a challenge both for describing the population activity and for considering how downstream circuits may decode, or read out, a specific variable at a specific time. Building on prior work^19^, we developed a new analysis, optimal targeted dimensionality reduction (oTDR; Methods), that discovered and quantified population-level representations, while suggesting a neurobiologically plausible readout mechanism. In brief, we identified *a priori* the behavioral variables relevant to performing the task: *benefit* (i.e., number of rewards offered), *choice* (i.e., accept or reject), and *expected reward* (i.e., interaction of benefit * choice, which reflected the outcome given the choice). We quantified how strongly each unit encoded the task-relevant variables by the coefficients derived from linear regression of trial-average firing rate (with each condition) on the task-relevant variables. Across the population of *N* units, the regression coefficients for a given variable defined an *N*-dimensional vector that we termed a “regression axis” (RA) or, alternatively, “population representation”.

We explored two classes of RAs: 1) *dynamic* RAs (dRAs) that were calculated for each time bin and tested how representations changed over the trial, as discussed below, and 2) *static* RAs (sRAs) that tested the suitability of a fixed readout for representing the task-relevant variables throughout the trial. The sRA for a given variable was calculated in the temporal epoch in which we hypothesized *a priori* that the variable was behaviorally relevant. Specifically, we reasoned that the animal encoded external value information during the offer period (0 – 0.5 s), and thus computed the BENEFIT sRA in this epoch. After the offer, the decision process proceeded without external information and thus depended on internal representations of the task variables—CHOICE and EXPECTED REWARD—which were computed during the post-offer, “work” period (0.5 s – …). We confirmed that our conclusions below were robust to a range of temporal epochs (Supplementary Figure 13). (We refer to the names of sRAs in small-caps and to task-relevant variables in lowercase, e.g., BENEFIT refers to the sRA representing the benefit variable.)

### Mixed selectivity for task-relevant variables in individual units

We examined the sRA coefficients to ask whether individual units were indeed selective for multiple variables (i.e., mixed selectivity), as in Figure 2e, or instead specialized for single variables (i.e., categorical selectivity)^5,43^. If specialized, then for a given variable, the absolute value of the corresponding sRA coefficient would be large for some units (i.e., highly selective), but concentrated near zero for most units (i.e., non-selective), resulting in a heavy-tailed distribution. However, the distributions of coefficients were not significantly different from Gaussian (Supplementary Figure 6), consistent with random assignment of coefficients to units, independent of the other variables (i.e., not specialized). Likewise, we did not observe an anatomical organization (e.g., clusters or gradients) of sRA coefficients (Supplementary Figure 7).

Across variables, we found that an individual unit significantly encoded two or more variables at a frequency that was a) much greater than expected if units were specialized for a single variable and b) statistically consistent with independent selectivity across variables (Supplementary Table 1). Second, we compared the magnitude of encoding between pairs of variables (Figure 3a,c). Although some units strongly encoded a solitary variable, these units did not form distinct clusters (i.e., points were not concentrated on the vertical and horizontal meridians, as predicted for specialized encoding) but rather were part of a continuum of selectivity. Third, we noted that apparent mixed selectivity could arise spuriously if unreliable neural responses led to non-zero coefficients masquerading as selectivity for multiple variables. By resampling trial-level data with replacement, we confirmed that the coefficients were highly reliable (Figure 3a,c, horizontal and vertical error bars rarely intersected zero; Figure 3b,d, filled bars; Supplementary Figure 8a,c; Supplementary Table 2, “Within-variable” section). In summary, we observed no statistical evidence that individual units were preferentially selective for single task-relevant variables.

In a separate analysis, we confirmed that mixed selectivity arose in single units, and was not, for example, an artifact of pooling specialized single-unit responses into multi-unit clusters of heterogeneous selectivity (Supplementary Figure 32 and Supplementary Table 1).

### Separability of low-dimensional representations

Thus far, we examined each unit in isolation. Next, we measured the relationship between each sRA, i.e., the population of regression coefficients. We found that pairs of variables were at most modestly correlated (Figure 3b,d, open bars; Supplementary Table 2). However, conventional statistics address the chance of mistaking orthogonal representations as correlated, whereas we were interested in the opposite extreme: whether two (possibly correlated) representations carried at least some independent information that could be separated at the level of the readout. Therefore, we defined two representations as *separable* when their correlation was significantly less than the hypothetical correlation between representations that were perfectly correlated (or anti-correlated, i.e., |r| = 1) but subject to imprecision in estimating the coefficients (see Methods). Indeed, the observed correlations were much less than could be explained by unreliable coefficients alone—that is, the representations were highly separable (Figure 3b,d, open bars vs. dashed lines; Supplementary Figure 8b,d; Supplementary Table 2). These separable representations provided a downstream observer with reliable, independent information about each encoded variable.

### Reading out population activity as low-dimensional representations

The mixed but separable encoding by individual units supported de-mixing of single-unit responses into variable-specific population representations. A crucial insight of oTDR was that the sRA coefficients for a given variable defined a dimension in neural space that best linearly represented the targeted variable. Projections of neural activity onto this dimension corresponded to a weighted sum of population activity that a downstream population could compute by tuning synaptic weights to the sRA coefficients. We replicated this neurobiologically plausible readout, or “activity” of the representation, by projecting the high-dimensional trial-average firing rate onto each sRA. To ensure independence between the projections, the sRAs were optimized to be orthogonal for this and subsequent analyses (Methods).

Examining sRA activity (Figure 4a,b), BENEFIT discriminated offer size rapidly and robustly after presentation, but the selectivity decayed abruptly and was absent by 1.5 s. Moreover, the representation did not discriminate the animal’s choice (thick and thin traces overlap). In contrast, CHOICE discriminated accept and reject choices beginning around 2 s for most offers and with increasing selectivity through the trial (see Supplementary Figure 9 for consideration of single-trial dynamics), but did not discriminate offer size. Thus BENEFIT and CHOICE de-mixed information about their respective variables. The apparent “bleed-through” of offer information into mid-trial CHOICE activity for reject choices (middle panel, thin lines), was in part because larger offers had later rejections and thus later choice selectivity (Supplementary Figures 2 and 9) and because the time-varying representations of benefit and choice (i.e., dRAs) overlapped during this mid-trial period (Supplementary Figure 26). For reject choices, sRA activity necessarily included post-rejection responses, a period of uncontrolled gaze, the contribution of which is discussed in Supplementary Figure 9.

Finally, EXPECTED REWARD discriminated offer size as early as 500 ms—as offer information transitioned abruptly from the orthogonal BENEFIT SRA—but did so only for accept choices. As such, EXPECTED REWARD integrated benefit and choice: activity reflected offer size for accept choices, but was undifferentiated for rejected offers. Temporally, EXPECTED REWARD coincided with the period of mid-trial rejections when no stimulus was present and thus may have provided the internal value representation necessary for mid-trial reevaluation (Figure 1c).

To test the temporal alignment between the sRAs and behavior more directly, we compared sRA activity for early and late rejections and found that choice selectivity emerged later on late-rejection trials, consistent with sRA activity reflecting the underlying decision dynamics (Supplementary Figure 10).

Regarding OFC’s visual sensitivity, the sRA activity for the singleton offer (which was excluded when discovering the sRAs) more closely matched that of conditions with similar value (i.e., 4- or 8-reward; Figure 4b, solid and dashed red lines; Supplementary Figure 11) than those with similar visual properties (i.e., 1-reward).

### Choice probability

Choices to the intermediate offers were inconsistent, suggesting the subjective valuation of these offers fluctuated across trials. We asked whether variability in the neural response to the offer, but *before the rejection*, explained this behavioral variability (so-called “choice probability”, CP), as expected under the widely-held theory that OFC value responses play a causal role in driving downstream decisions^44^. Because our population analysis relied on trial-average responses pooled responses across sessions (and thus could not exclude post-rejection activity on single trials), we measured CP at the individual-unit level (Supplementary Figure 12). We found no evidence that the neural response following a given offer predicted the upcoming choice, either on average or proportional to a unit’s contribution to the BENEFIT sRA (an approximation of the population representation of offer value on single trials). At the population level, CHOICE activity did not discriminate choice until *after* the median rejection time (0.92 or 1.3 s, monkey N or K).

### Validity, specificity and sensitivity of low-dimensional representations

Here we assess the suitability of a static, low-dimensional read out (i.e., sRA) for representing a task-relevant variable. Specifically, the relevant variance explained by an sRA should be greater than the variance explained by alternative dimensions (i.e., valid), unrelated to other variables (i.e., specific), and capture all available relevant variance, obviating the need for additional dimensions (i.e., sensitive). We defined *relevant signal variance* (RSV; Methods) as the portion of variance explained by a given sRA (Supplementary Figure 14) that was linearly related to the sRA’s targeted variable (Figure 4c,d).

#### Validity

We compared the RSV for a given sRA to that expected for an arbitrary dimension, modeled as a set of random vectors. Critically, the density of random vectors was *not* evenly distributed throughout the *N*-dimensional space—a seemingly intuitive procedure that would over-represent rare dimensions and thereby overestimate the rarity of the sRA—but instead was enriched in regions commonly occupied by the *N* neurons (Methods). The resulting p-values were more conservative (i.e., larger) than had the random vectors been distributed evenly. We found the sRAs explained significant relevant variance during three, key periods (Figure 4e,f): encoding external value information from 0.1 to 1 s (BENEFIT), sustaining value information conditioned on choice from 1 s onwards (EXPECTED REWARD), and representing the chosen action from 1 s onwards (CHOICE).

Separately, we confirmed that sRAs derived exclusively from single units explained comparable task-relevant variance (Supplementary Figure 33-34).

#### Specificity

The portion of variance unrelated to the targeted variable (*irrelevant signal variance*, ISV) was generally small and below chance levels (Figure 4c-f, dashed curves), indicating that the sRAs were specific to their targeted variables. Furthermore, very little of the irrelevant variance was related to the other task variables, thus minimizing “leak” between variables that would confuse a downstream observer (Supplementary Figure 15).

#### Sensitivity

We considered two hypotheses to explain the time periods when the sRAs failed to detect significant variance. First, were the task-relevant variables simply not represented by the population during these times? For instance, the gradual increase in RSV by CHOICE may have been due to a gradual influx of choice information to the OFC population at large. Second, the optimal representation may have rotated to alternative dimensions during select times. For instance, the population may have represented choice with constant strength throughout the trial, but done so along a rotating dimension that gradually aligned with the CHOICE sRA.

To distinguish these two hypotheses, we defined dynamic, low-dimensional representations by discovering independent sets of non-orthogonalized dRAs for benefit, choice, and expected reward in non-overlapping, 200 ms time bins. For a given dRA, the magnitude (prior to normalization) defined the strength of population encoding (regarding hypothesis 1) and the direction defined the relative contributions of individual units (regarding hypothesis 2) to the momentary optimal representation of the corresponding variable.

The dRA magnitude was indeed variable (Figure 5a,b), consistent with hypothesis 1. In particular, the representation of benefit was large early in the trial and decayed rapidly, while the representation of choice emerged around 1 s and gradually increased throughout the trial.

To evaluate hypothesis 2, we measured the change in dRA direction as the angle *θ*_*ij*_ between dRA(*i*) and dRA(*j*) computed at times *i* and *j* for the same task-relevant variable. Small angles corresponded to highly similar directions and indicated that the contributions of individual units were highly consistent across the trial. We observed broad periods in which the dRAs were qualitatively similar to each another (Figure 5c,d, areas of warm colors), such as the representation of choice from ∼1 s onwards. However, for some variables, we observed multiple periods of similarity that were dissimilar from one another (e.g., Figure 5c,d, representation of benefit early vs. mid-trial; corresponding periods marked with vertical brackets in left panels of Figure 5e,f), implying that a static dimension (i.e., sRA) could capture the population representation for a *portion* of the trial, but not the *entire* trial. To quantify the population signal *un*explained by the sRAs (i.e., “left on the table”), we could not directly compare the sRAs and dRAs in terms of variance explained (projections onto the non-orthogonal dRAs would not be independent). Instead, we measured the extent of dRA magnitude uncaptured by the sRAs, which was generally small and never greater than expected for a random set of three static dimensions (Supplementary Figure 16).

In summary, we found qualitative evidence for both hypotheses: 1) the magnitude of population-level representation fluctuated over the trial and was largely captured by the sRAs; and 2) the particular contributions of individual units to a given representation were consistent for discrete periods, but for some task variables, changed abruptly mid-trial, limiting the generalizability of the sRAs across all times.

Finally, we compared the sRAs to other low-dimensional subspaces, such as those spanned by the top three principal components (Supplementary Figure 17) or by the dimensions capturing the population’s time-varying signal (i.e., common-condition response; Supplementary Figure 18).

### Dynamic representations: stability and alignment to task events

Thus far, we observed that the time-varying dRAs for a given variable were qualitatively similar for discrete periods of the trial. In this section, we formalized these impressions by systematically delineating periods of similarity and then testing these periods for statistical *stability*—unusually high or protracted similarity—which indicated that a representation may have been specialized for the current task.

We noted that the periods of similarity were not uniform across time or variables, and therefore would not be well-described by a single, canonical “similarity profile” (e.g., Ref^45^; tested in Supplementary Figure 21). To more flexibly capture time-varying periods of stability for each variable, we fit boxcar functions to each row of the heat maps in Figure 5c,d (see Supplementary Figure 22 for fits). The temporal span of each boxcar indicated the period of putative stability, while boxcar height *θ*_boxcar_ indicated the average similarity of the representations during that period (Supplementary Figure 23a,b). (Note: boxcars were fit to 90° - *θ*_*ij*_, thus periods of high similarity had large values of *θ*_boxcar_ but small values of *θ*_*ij*_.)

We next developed a statistical framework to assess the putative stability captured by the boxcars. For instance, how surprising was it that the choice representation changed by only 20° in a 3 s interval? How much would a random neural dimension change during that same interval? The population’s correlational structure could generate similarity trivially by limiting how much a representation *could* change. For instance, correlations between units, or dimensionality, restricted the subspace a representation could occupy. At the limit, if the dimensionality were unity, then a representation could not vary and would be “similar”, indeed identical, across all times. Separately, correlations in an individual unit’s response over time, i.e., temporal smoothness, restricted how quickly a representation could change. If temporal correlations were very high, then even if the dimensionality permitted dissimilar representations, the population may not have been able to transition to an alternative representation within the span of a trial.

To test statistical stability, we compared our observations to identical metrics made on synthetic firing rate data from a hypothetical neural population that did *not* encode the task variables but preserved the correlations between units and across time (see Methods). In each synthetic dataset, we computed the average similarity 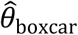 during each period of putative stability, where periods were defined by the boxcar fits from the veridical data. We defined a period as statistically “stable” when the observed similarity (*θ*_boxcar_) was significantly greater than the null values of 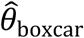 compiled across synthetic datasets (i.e., p(*θ*_boxcar_) < 0.01; Supplementary Figure 23e,f).

Most representations in OFC were statistically stable during discrete periods when the encoded information was behaviorally relevant (Figure 5e,f, colored horizontal bars). Specifically, the representation of benefit was stable for a period of ∼0.5 s (Figure 5e,f, left panel, red bracket) coinciding with the offer presentation and also when benefit was encoded most strongly (i.e., peak dRA magnitude; Figure 5a,b). Subsequently, a new representation of benefit emerged (blue bracket) that, while stable for ∼3 s, was dissimilar from the earlier period and weaker in magnitude. The representation of choice was stable from ∼1.5 s (after most choices were rendered) to the end of the trial. For monkey N, the representation of expected reward was stable from ∼1 s (around the median rejection time and coinciding with the decay of offer information from the earlier benefit representation) to the end of the trial. For monkey K, however, stability of expected reward was inconsistent: while the boxcar fits identified periods of *similarity* during the work period (Figure 5f, dark gray horizontal bars), these periods only occasionally achieved statistical *stability*, i.e., p(*θ*_boxcar_) hovered around chance levels (Supplementary Figure 23f). Our conclusions were robust to the choice of time bins (Supplementary Figure 24).

Separately, the earlier sRA analysis suggested that the time-varying representations of *different* task-relevant variables may have overlapped at certain times (e.g., CHOICE activity discriminated offer size mid-trial; Figure 4a,b), even though the corresponding sRAs—based on broader time periods—were highly separable. Indeed, we found significant similarity between dRAs of different variables during limited periods of the trial (Supplementary Figure 26).

In summary, the low-dimensional representations of the task-relevant variables were statistically stable, i.e., the specific contributions of individual units were more consistent than expected by the data’s dimensionality and temporal smoothness. And yet the timing of these stable periods varied across the task variables, aligning with the concurrent task demands for which the representation was behaviorally relevant.

### Representations transitioned abruptly between trials

Given the magnitude of the static representations (Figure 4c,d) and stability of the dynamic representations (Figure 5c,d) during the trial, we asked whether task-relevant information was preserved across trials. Although our task design did not require cross-trial integration of choice or reward history, the animals nonetheless may have used information from the previous trial to inform upcoming choices, a phenomenon observed even in tasks that do not incentivize history-dependent choices^5,46,47^.

We began by examining the static representations (sRAs) during the transition from one trial to the next. This required defining a new analysis window, the *previous-trial epoch*, aligned to fixation on the present trial and extending retrospectively through the previous trial. Intuitively, we expected the sRAs to explain variance related to the task-relevant variables in the previous trial, just as they had for the present trial. However, to perform this sanity-check, we first had to compute *previous-trial responses* by re-sorting the trial-level neural responses according to the conditions of the previous trial (see Methods for details and rationale). As before, we then projected previous-trial responses onto the sRAs and measured the RSV explained with respect to the *previous-trial* variables: previous benefit, previous choice, and *experienced* reward (previous benefit * previous choice). As expected, the sRAs significantly explained variance related to the previous-trial variables during the previous-trial epoch (Figure 6a-d, left panels), just as they had explained variance related to present-trial variables during the present-trial epoch (Figure 4c-f; Figure 6a-d, right panels). In addition, the sRAs maintained a representation of choice beyond the work period and into the reward period and ITI (Figure 6a-d, orange curve relative to horizontal gray bar indicating previous reward times).

However, just before fixation on the present trial, the task-relevant signals disappeared precipitously from the present-trial sRAs (Figure 6a-d, left panels, orange and, to lesser extent, blue curves as approaching Time = 0), as though the system were clearing its cache to accommodate the upcoming present-trial information. At first blush, there appeared to be no continuous representation of the task-relevant variables across trials, which puzzled us given our (unpublished) behavioral observations: animals were more likely to reject the present offer after accepting the previous offer. And so we wondered if previous-trial signals were transmitted in other dimensions, outside the subspace defined by the sRAs. Using oTDR, we searched deliberately for alternative dimensions carrying task-relevant information between trials. Specifically, we computed *previous-trial* sRAs—PREVIOUS BENEFIT, PREVIOUS CHOICE, and EXPERIENCED REWARD—based on the previous-trial responses during the first 0.5 s of fixation on the present trial, a time when the animal under behavioral control but not yet exposed to the present offer.

Remarkably, just before the animal fixated on the present trial, the new set of previous-trial sRAs suddenly and robustly represented the events from the previous trial, filling-in the gap left by the present-trial sRAs (Figure 6, e-h vs. a-d), as though information were passed between distinct sets of neural dimensions (confirmed below). During the present trial (Figure 6e-h, right panels), PREVIOUS CHOICE (orange curves) continued to encode the previous decision through the fixation period (−0.5 to 0 s), offer period (0 to 0.5 s), and until the time of most rejections (∼1 s). Likewise, in monkey K, PREVIOUS OFFER and EXPERIENCED REWARD significantly encoded their respective variables through all or part of this pre-rejection period (green and blue curves, respectively). As such, the timing of the previous-trial representations was sufficient to bias the present choice based on the previous trial’s outcome, as observed behaviorally.

Looking back in time, the previous-trial sRAs did *not* explain variance related to the previous-trial variables during the previous trial itself, a time when the present-trial sRAs represented the task-relevant variables (Figure 6, left panels, Time << 0, e-h vs. a-d). This implied that the present- and previous-trial sRAs spanned separate subspaces. Indeed, previous- and present-trial representations were at most weakly correlated and corresponded to highly dissimilar, nearly orthogonal dimensions, (r = 0.02 to 0.23; *θ* = 78° to 88°, Supplementary Figure 27, Supplementary Table 3). That is, OFC maintained distinct representations not only of offer, choice and reward events, but also of the relative trial in which those events occurred, allowing a downstream readout to distinguish previous-trial information (e.g., PREVIOUS BENEFIT) from its present-trial counterpart (e.g., BENEFIT).

As above, we confirmed that individual units were selective for multiple previous-trial variables, while previous-trial sRAs were specific to their targeted variable and largely separable (Supplementary Figure 28-31; Supplementary Table 4).

## Discussion

We found that single neurons in macaque OFC represented key task-relevant variables—benefit, choice, and expected reward for both the present and previous trials—while the animals made subjective cost-benefit decisions requiring sustained effort. For individual neurons, the encoding of value and choice were mixed, and the time course of encoding varied widely across neurons. However, using a set of new analysis and statistical techniques, we de-mixed the task-relevant signals into static low-dimensional representations that were separable at the level of the population. Additionally, the timeseries of dynamic representations were statistically stable during periods when the information was behaviorally relevant, and then transformed abruptly at key task events as information transitioned between dimensions. Our findings suggested OFC may organize—forming and disbanding coordinated combinations of neurons—on behavioral timescales to represent and manipulate information relevant to concurrent behavioral demands. Moreover, the nature OFC encoding—mixed single-neuron selectivity and stable population representations—facilitated a neurobiologically plausible mechanism by which downstream circuits could tune synaptic strengths to selectively read out specific task-relevant signals during specific time periods.

### Categorical vs. mixed selectivity

Prior reports have argued for *categorical* selectivity in OFC—i.e., individual units specialized to encode a single variable—either implicitly by classifying otherwise mixed responses into single-variable categories, or explicitly by comparing univariate regressions performed independently for each variable^5–8,15^. In contrast, when variables competed for variance in a multivariate model, we found that units encoded two or more variables at rates equal or above chance, consistent with broad evidence for *mixed* selectivity across cortical areas^9–11^ and shown indirectly for OFC^43^.

Moreover, we found the mixed representations were *separable* at the level of the population. That is, the degree to which a given neuron encoded variables X or Y was independent, consistent with random assignment of encoding strength across neurons. In contrast to cases where the encoding strength of variables is correlated in single neurons (e.g., visual motion and choice selectivity in area MT^48,49^), separable representations increase coding capacity, enable flexible readout of a single variable depending on task demands, and support non-linear combinations of variables (such as expected reward)^12,13,33^.

### Unstable vs. stable selectivity

Several studies in rodent cortex have reported sequences of activity, where the neuron(s) selective for a given task variable changed on the order of 10’s of milliseconds^14,17,50–52^. This *unstable* selectivity may be well-suited for representing variables that vary smoothly in time, such as spatial position, or for generating an eligibility trace for learning^53^. However, for a conserved downstream population to read out these unstable representations, it would have to select for a different combination of neurons at each time point and do so with knowledge of the current trial time. Moreover, the sequences were highly specialized for a single variable. Thus, it is unclear how this coding scheme—categorical and unstable selectivity—would support a readout that could flexibly select among multiple variables and do so at different timescales depending on task demands.

In contrast, the population representations in OFC were highly *stable*, i.e., the selectivity of a given neuron was consistent during the behaviorally relevant period. This permits a downstream circuit not only to select for the representation of a single variable, but to do so via a *static* set of synaptic weights (as prescribed by the sRA coefficients^54^). Unlike the rapid, within-trial updating of readout weights implied by unstable selectivity, these weights could be tuned gradually during learning and then remain constant during mature behavior, a neurobiologically plausible mechanism^55^.

The stability of the OFC population representations was not uniform across the trial. Rather, the representation of a given variable was stable for a period, then, at key task events, changed abruptly to a new, stable representation. This time-dependence could facilitate temporal gating—that is, not just *which* variable is read out, but also *when* that variable influences downstream computation. For instance, benefit was encoded along two, highly dissimilar dimensions during the offer and work periods (Figure 5e,f, brackets). Therefore, a read out tuned to the earlier representation (which corresponded to the BENEFIT sRA) would be minimally sensitive to the later representation, and vice versa. This would permit the *same* information (i.e., benefit) to drive *different* functions. For instance, the early representation could inform the initial decision, while mid-trial benefit could integrate with choice to compute expected reward (see Supplementary Figure 26). A similar temporal gating could apply to choice information, with the CHOICE SRA informing credit assignment during the outcome, while PREVIOUS CHOICE biased the decision on the next trial. In motor cortex, analogous gating mechanisms may select for planning vs. execution signals^35,56^.

A recent report described qualitative similarity between representations of a single variable in lateral PFC^45^. However, to our knowledge, our paper is the first systematic analysis of stability in OFC and the first to offer, for any brain region, a statistical definition of stability and show how stability depends on time and task variable.

#### On classifying stability

At short timescales, the distinction between ‘stable’ and ‘unstable’ representations can be subjective. For instance, we argued that the early benefit representation (Figure 5c,d) was stable for ∼1 s, while another observer may describe it as a slowly evolving sequence. Going forward, developing objective tests to distinguish these cases will be critical, particularly given the prevalence of apparent sequences in the rodent. Nonetheless, the extended stability of most representations (e.g., mid-trial benefit, choice, and expected reward) is unequivocally distinct from sequence-like encoding.

The type of stability we describe in OFC (i.e., consistent selectivity within the trial) contrasts other types of “stability” in the literature. For instance, selectivity may change within the trial, but, for a given time in the trial, remain consistent across experimental sessions^50,51^; or selectivity may be consistent across behavioral contexts, irrespective of temporal stability^6^.

### Diversity of OFC function reconciled under ‘cognitive map’ framework

A multitude of functions have been proposed for OFC—and, in humans, ventromedial PFC—on the basis of seemingly disparate phenomenon observed across physiological and lesion studies in primates and rodents^16^. Across several attempts to reconcile OFC’s diversity under a single, unifying account^57–59^, one promising theory posits that OFC encodes a cognitive map, i.e., a general purpose framework for representing the set of cognitive and behavioral states relevant to the subject’s current goals^60–62^. As behavioral demands change rapidly, so must the specific states represented by a cognitive map, thus reconciling OFC’s seemingly disparate functions as merely time-varying instantiations of a general process. Consistent with this prediction, within the same OFC neurons, we found representations of key variables that support many of OFC’s proposed functions during discrete periods when the variables were behaviorally relevant.

In particular, BENEFIT transformed external stimulus information to a representation of value early in the trial^15,28,63–69^, which, after 1 s, decayed rapidly from the BENEFIT sRA as the dimension representing benefit rotated abruptly.

Simultaneously, EXPECTED REWARD began representing offer size for accept choices throughout the work period, providing real-time expectancy signals that may have supported on-the-fly reevaluation during the work period and, in learning contexts, computation of prediction errors during the outcome period^70–73^.

CHOICE began representing the decision mid-trial—after most rejections and after EXPECTED REWARD reflected the choice—suggesting that CHOICE unlikely drove the animal’s pending decision. Instead, CHOICE represented the selected action, which could facilitate credit assignment^74,75^. In addition, CHOICE activity increased smoothly throughout the trial, potentially representing timing information to compute time-dependent benefit or cost signals during reevaluation.

Finally, between trials, CHOICE transitioned abruptly to a new, nearly orthogonal representation, PREVIOUS CHOICE, that sustained choice history through the next offer, sufficient to drive trial-to-trial choice dependencies, as we observed in our unpublished data and proposed by others^5,76^.

### Does OFC drive trial-to-trial variability in choice?

Despite the widespread model that OFC integrates external value information with subjective preferences to drive decisions^15,16,77–80^, a key prediction—that variation in OFC responses accounts for variability in choice (i.e., choice probability, CP)—has not been reported. A recent study argued that rapid fluctuations in OFC reflected momentary preferences, but lacked the behavioral variation needed to link neural activity and preference^81^.

In contrast, our animals accepted intermediate offers inconsistently, and thus we were well-poised to assess CP in OFC. Nonetheless, we did not observe choice predictive responses. Several factors may have prevented us from observing CP: low signal-to-noise in the individual-unit analyses, the animals’ extensive training history^82,83^, and challenges in interpreting CP^84^.

Alternatively, OFC may not play a causal role in transforming external value cues into an initial behavioral policy. For instance, OFC may represent subjective preference prior to integration with (and independent of) external value information^5^. Another possibility is that OFC maintains an internal representation of value that, during behaviors requiring sustained effort, informs whether to continue the initial policy or change one’s mind midstream. This putative role of OFC in reevaluation would not be observed in the vast majority of tasks used to study OFC—tasks in which the decision is rendered by a brief, all-or-nothing response. Indeed, optogenetic inactivation of OFC in the rat had no effect on choices in a classic value-based task of this sort^85,86^. However, when the same rats were asked to weigh reward against the cost of sustained effort, OFC inactivation disrupted normal value-based behavior.

In our present task, animals had to sustain their initial decision over several seconds to receive reward, and at times reversed their decision mid-trial at rates proportional to the expected reward. Notably, during this period, the EXPECTED REWARD sRA maintained the value information needed to inform this reevaluation, though the serial nature of our recordings precluded linking variability in EXPECTED REWARD activity to single-trial choices. Going forward, sustained-effort tasks paired with simultaneous recordings and/or targeted manipulations of OFC activity would test the role of OFC in reevaluating choices mid-execution.

### Decisions rendered abruptly vs. over time

As discussed, our novel task design permitted subjects to make an initial commitment to accept an offer and then, because the decision was rendered via sustained effort, to reverse their choice midstream. This behavior is akin to deciding to climb a banana tree or attend graduate school, choices which are still binary in nature, but can be reevaluated and reversed *after* initiating the behavior. In contrast, in conventional value-based tasks that require a brief, all-or-nothing response, all deliberation occurs *before* initiating the behavior, which in turn is executed abruptly and irreversibly, akin to choosing amongst candies in a box of chocolates. Such tasks do not encourage (nor designed to observe) reevaluation of the initial decision and thus may not invoke the neural representations of value that might otherwise support this function.

Nonetheless, sustained-effort tasks engage the same fundamental processes—valuation, comparison, and (at least initial) commitment to a behavioral policy—as conventional choice tasks. Others have leveraged these parallels to study, for instance, how value signals in ACC evolve during sustained, effortful engagement^87,88^. In the present study, the representations of BENEFIT, EXPECTED REWARD, and CHOICE were directly analogous to the responses of Offer Value, Chosen Value, and Taste, respectively, reported in the classic studies of OFC by Padoa-Schioppa et al.^15^ A key distinction is that, in the earlier studies, the role of Chosen Value and Taste in driving the current choice was less obvious, whereas in the current task, EXPECTED REWARD and CHOICE may have informed whether to continue the current policy or abort mid-trial. We believe sustained effort-based decision tasks, such as our novel design, offer an essential bridge between conventional choice tasks and more ethological behaviors that may rely preferentially on OFC representations.

Critically, both conventional and effort-based value tasks require goal-directed use of value to produce flexible behavior. In this way, the present task differs from studies of self-control or response inhibition^89,90^. Though these processes are almost certainly at play here, the smooth, graded relationship between offer size and behavior, clearly showed that the monkeys used value information to decide whether to reject an offer outright, reverse their initial decision deliberately, or apply less ‘self-control’ in executing their decision. In so doing, the behavior captures the ethological case in which agents maximize reward per unit *cost*, not only per unit *time*. For instance, one may reject an offer of $1 to carry a heavy package up several flights of stairs simply because the small reward is not worth the high cost. One may even start up the stairs but reject the cost as too onerous after a flight or two. One is not maximizing absolute income, but is conserving resources, which are almost certainly finite and must be allocated judiciously.

### A roadmap for analyzing high-dimensional neural data

As more studies analyze high-dimensional neural data, so grows the need for a common set of measurements and statistical framework^21^. We offer a systematic, principled and statistically rigorous roadmap for population analysis that can be applied, either cohesively or modularly, to any neural population. Our approach not only de-mixes low-dimensional representations of task-relevant variables, but, critically, emphasizes the relationships between representations, both between different variables at the same time point (i.e., separability) and between different times (i.e., stability). To our knowledge, our paper is the first to assimilate these aspects of population coding—dimensionality reduction, separability, and stability—into a single statistical framework. Moreover, we are the first to apply contemporary population analyses of any sort to OFC.

Unlike other de-mixing techniques, oTDR synthesized several assumptions—regression, orthogonalization, weighting by observation count, and noise-reduction—in a single objective function, thereby discovering the optimal linear representations of the variables *given the model assumptions.* An earlier method applied subsets of these assumptions, but did so serially and thus approximated a solution to the original objective^19^. An alternative method could not accommodate unbalanced designs in which not all combinations of task variables are observed^20^, as frequently encountered in decision-making studies. Moreover, oTDR can generalize to any arbitrary number of variables and epochs, while computing dimensions for each variable in one or more epochs and assuming orthogonalization between any subset of dimensions. As such, oTDR is a general-purpose method suitable for discovering static representations of known relevant variables in high-dimensional data.

Independently of oTDR, the metrics we developed—including formal definitions of separability and stability, as well as assessments of sensitivity and specificity (i.e., RSV, ISV and partial RSV)—are applicable to any set of dimensions, and thus any dimensionality reduction method.

Finally, we introduced novel applications of recent statistical tools^21,35^ to test the significance of these metrics (again, applicable to any de-mixing technique). A rigorous statistical framework is crucial to rule-out epiphenomenal findings attributable to the population’s intrinsic, low-level features, to which high-dimensional systems are particularly susceptible^21^. Our methods estimated the contribution of these features more accurately—and estimated statistical significance more conservatively—than prior approaches.

Specifically, our novel tests of representational sensitivity and specificity compared the variance explained by the sRAs to that of random dimensions that preserved the data’s dimensionality. Prior methods (by distributing null dimensions isotropically) assumed all dimensions were equally likely and thus overestimated the significance of the high-variance dimensions^20,33^. Similarly, in our novel statistical test of stability, we generated surrogate firing rate data that preserved the temporal smoothness *and* dimensionality of the original responses. In contrast, typical shuffling procedures—that randomize across *either* time points *or* task conditions—maintain *either* the dimensionality *or* smoothness of the data, respectively, but not both, and therefore are prone to overestimate the significance of a given finding.

The present toolkit may be useful to other high-dimensional datasets, such as large-scale simultaneous recordings^39,40,50^ or surveys across multiple brain areas, where resolving the relative contributions and temporal sequence of representations has been difficult at the single-neuron level^9,11,77^. Finally, in calcium imaging and fMRI data, our tools may discover meaningful low-dimensional representations and assess their similarity across time *and* space.

## Supplementary Figures

### Behavior

**Supplementary Figure 1.**
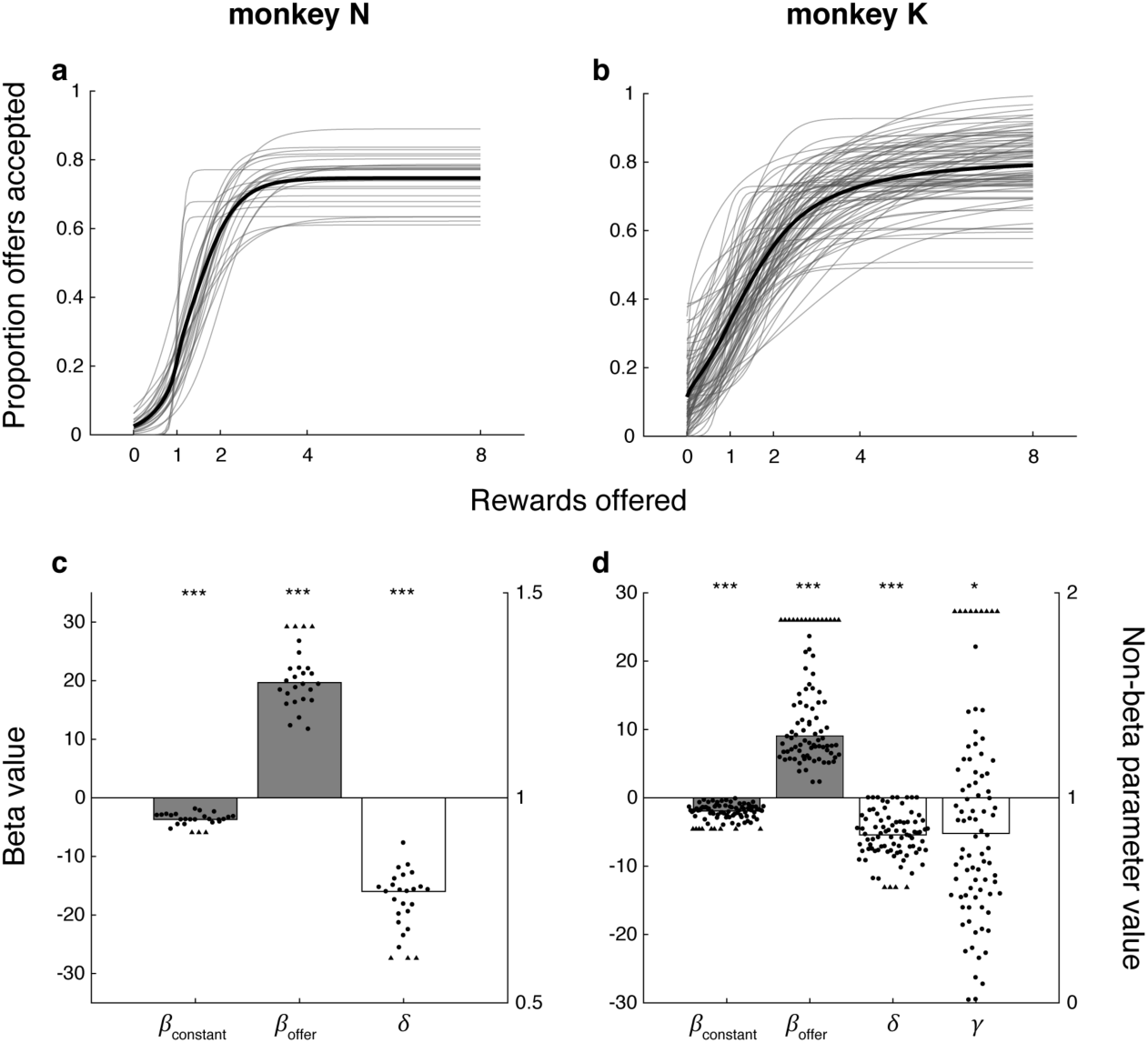
Logistic regression of choice behavior. **(a,b)** The logistic function (Eq. 1) fit to the choice behavior from individual sessions is plotted (thin gray curves) as a function of offer size for monkeys N (a) and K (b). The average curve across sessions (thick black curve) was computed as the mean of the individual curves for visualization only, and does not represent a separate logistic function fit to the aggregate data. The smoothly increasing accept rate within individual sessions demonstrated that valuation of the offer varied from trial to trial, not merely between sessions (as demonstrated in Figure 1b). Alternatively, an abrupt step function would have suggested that animals applied a constant threshold for offer size, above which they accepted all offers. **(c,d)** The logistic model included coefficients for offer size (*β*_offer_) and a constant (*β*_constant_) and additional parameters for a sub-unity saturation point (*δ*) and sublinear utility function (*γ*, monkey K only). For monkeys N (c) and K (d), bar height represents the median parameter value across experimental sessions for beta (*β*; gray bars) and non-beta (open bars) parameters, which reference the left and right ordinates, respectively. Parameter values from individual sessions are shown as circles with outliers (triangles) plotted not-to-scale (all values were included in median calculation and hypothesis testing). Spread along the horizontal axis is arbitrary and for display purposes only. Asterisks indicate probability that the true median equaled the null hypothesis (horizontal line; * p < 0.05; *** p < 0.001; Wilcoxon signed rank test). Null hypotheses for beta and non-beta parameters were zero and one, respectively. Specific values for medians and associated p-values for monkey N or K, respectively, are as follows: *β*_constant_ = -3.7 or -1.9, p < 10^−6^ or 10^−24^; *β*_offer_ = 20 or 9.0, p < 10^−6^ or 10^−24^; *δ* = 0.77 or 0.82, p < 10^−6^ or 10^−24^; and, for monkey K only, *γ* = 0.83, p < 0.013. In summary, both monkeys were significantly more likely to accept an offer the larger its magnitude, though with an overall tendency to reject all offers and a maximal acceptance rate for the largest offers around 80%. In addition, monkey K exhibited a sub-linear utility function as reflected in the fitted values of the parameter *γ* (see Methods), i.e., the increase in utility for a given increase in offer size became less as offer size increased.

**Supplementary Figure 2.**
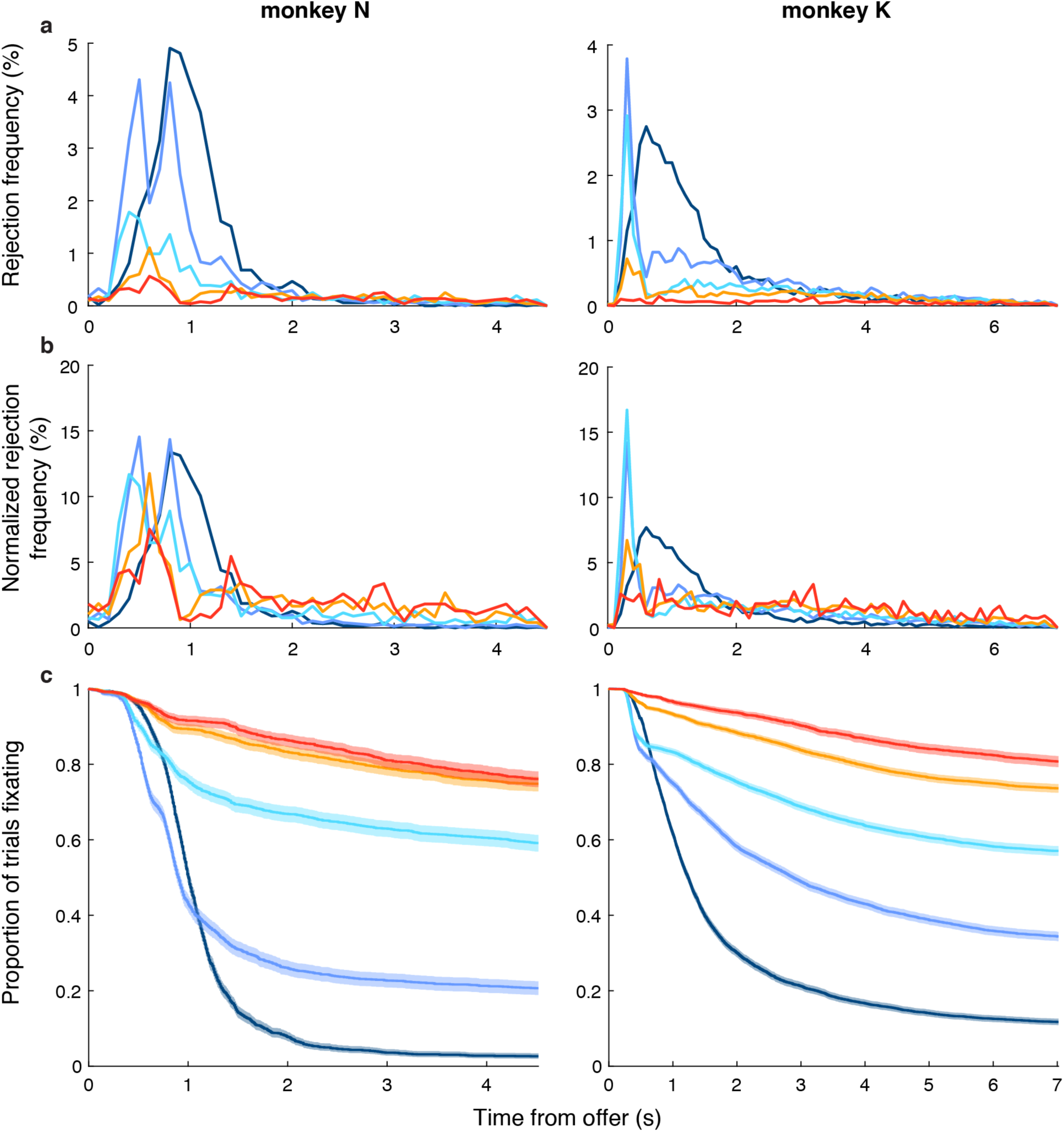
Rejection timing. **(a,b)** The frequency of fixation breaks (i.e., reject choices) per 100 ms time bin is shown for each offer size (colors as in Figure 1) as a function of time from the onset of the offer period as the percentage of all rejections (a) or as percentage of rejections for the given offer size (b) for monkeys N and K (left and right panels, respectively). **(c)** The Kaplan-Meier estimator of the fixation survival function is shown for each offer size (colored lines as in (a,b)) as the proportion of trials in which the animal is fixating as a function of time from the onset of the offer period. Shading represents the 95% confidence interval. The derivative of the survival function conditioned on maintenance of fixation approximates the hazard rate functions shown in Figure 1c. The hazard rates scaled inversely with offer size (Cox proportional hazards, excluding 0-rewards: *β*_offer_ ± s.e.m. = -0.29 ± 0.0098 or -0.29 ± 0.0069, p < 10^−16^ or 10^−16^, monkey N or K, respectively; see Supplementary Figure 3 regarding 0-reward offers).

**Supplementary Figure 3.**
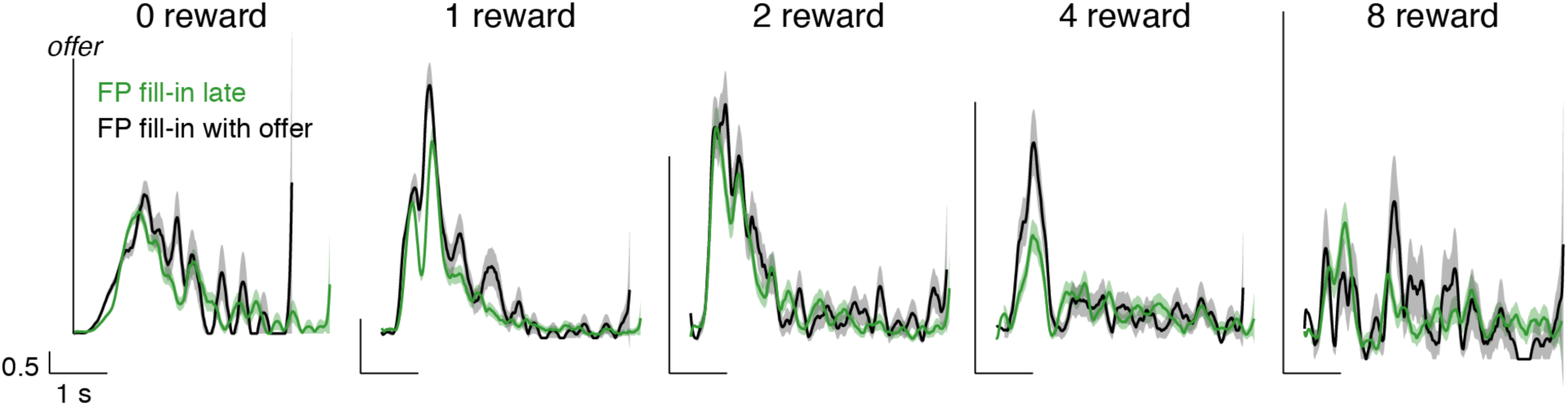
FP fill-in timing and effect on rejection time. In a subset of early experimental sessions for monkey N, the fixation point (FP) transitioned from an open annulus to a filled circle at the beginning of the work period (FP fill-in late), instead of the beginning of the offer period (FP fill-in with offer), as in all other sessions. Therefore, for these early sessions, the animal did not receive an explicit cue as to the beginning of the offer period on 0-reward offers (in constrast, on non-zero offers, the onset of the offer icons cued the offer period). We reasoned that this 0.5 s delay on 0-reward offers may have contributed to a delay in responding to the zero-reward offers in particular, and potentially to any size offer. Here we plot the hazard rate of fixation breaks as a function of time from the offer (as in Figure 1c), with the rate shown separately for FP fill-in late (green curves) vs. FP fill-in with offer (black curves) sessions. Shading represents standard deviation of hazard rate. Vertical and horizontal scale bars indicate 0.5 hazard rate and 1 s duration, respectively. For all offer sizes, we observed no appreciable differences in the temporal profile of hazard rate depending on the timing of the FP fill-in cue. We concluded that the animal did not rely strongly on the FP fill-in cue in determining the timing of rejection responses. Nonetheless, we observed a marked delay in hazard rates for the 0-reward offer compared to all other offers for both animals N and K (see Figure 1c and Supplementary Figure 2). The cause of this delay was unclear. The 0-reward cue may have been less salient, contributing to slower reaction times. However, intuitively, the distinction between FP fill-in late vs. with offer would likely have been more salient, and yet the FP timing did not account for the slower responses to 0-reward offers. This suggested that a temporal component of the decision process was fundamentally different for zero vs. non-zero offers and argued for analyzing the timing of 0-reward offers separately (as was done for the Cox proportional hazards analysis in Supplementary Figure 2).

### Anatomy

**Supplementary Figure 4.**
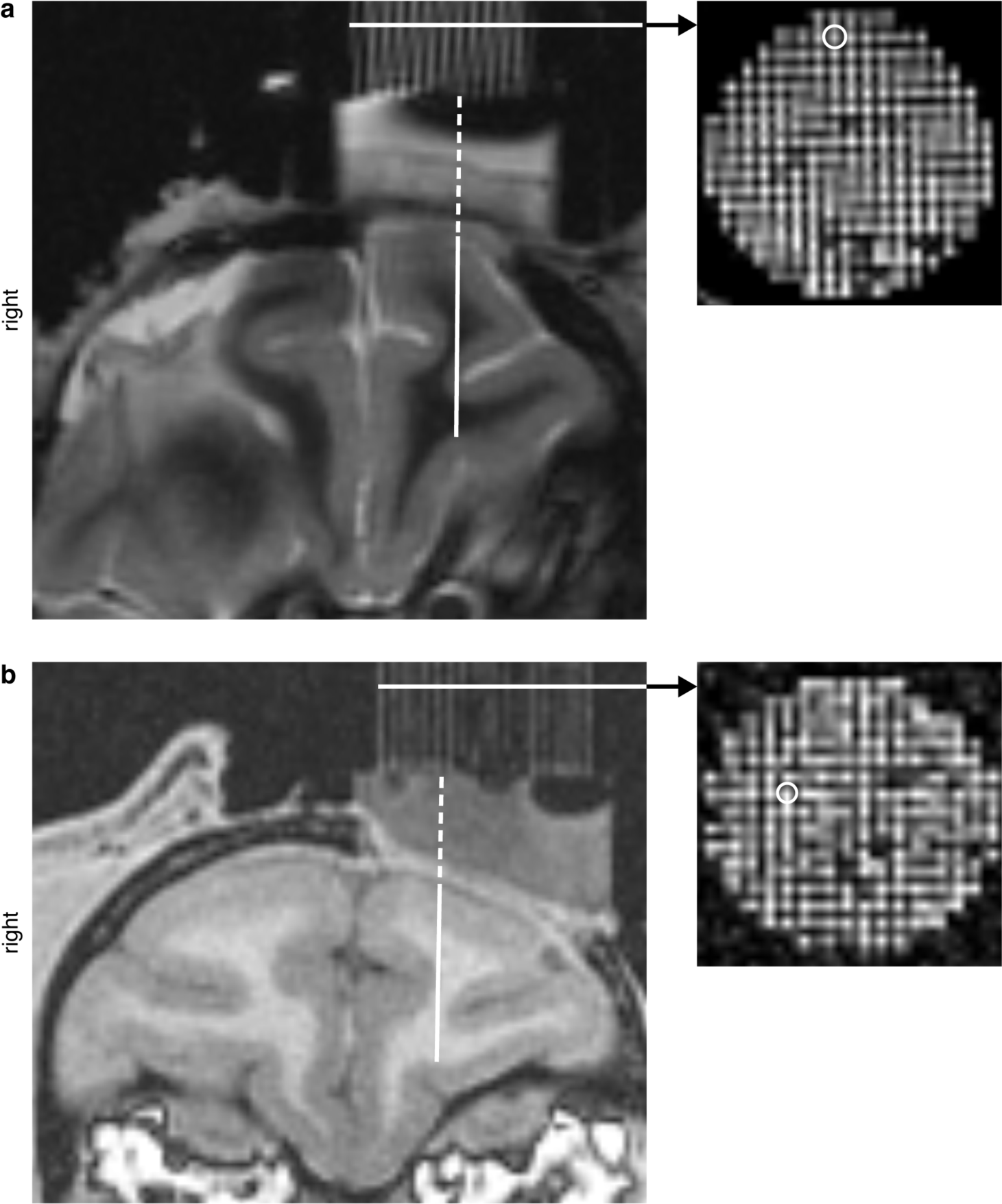
MRI localization. Anatomical MRI sequences were acquired with recording grid in place prior to physiological recording. **(a,b)** T2- or T1-weighted coronal slice is shown at the median anterior-posterior extent of recorded sites for monkey N (a) or K (b), respectively. Animal’s right side is shown on the left side of the image, per radiological convention. The inset shows an axial slice through the recording grid (white horizontal arrow), with a representative grid hole circled corresponding to grid position 2A or 0G (43.1 or 36.6 mm anterior to the interaural line, measured via intra-surgical stereotaxic coordinates; 6.7 or 6.3 mm left of midline, measured from the post-surgical MRI) for monkey N or K, respectively (see Supplementary Figure 7). Recording grid was filled with salinized agarose solution to facilitate contrast on MRI. By registering the axial slice through the recording grid and the coronal slice containing the selected grid position, we traced the virtual electrode trajectory from the bottom of the recording grid to the dorsal surface of cortex (vertical dashed white line), and then from the dorsal surface to the ultimate recording site (vertical solid white line) for individual electrode penetrations. We estimated position along the trajectory with a calibrated micromotor. Here we show the virtual trajectory to the most dorsal aspect of OFC reached by the representative grid positions (above): the fundus or lateral bank of the medial orbital sulcus at 14.5 or 13.2 mm below the dorsal surface for monkey N or K, respectively. Note that the coronal slice prescription for monkey N was oblique to the coronal plane with the left hemisphere rotated anteriorly, accounting for the asymmetry in (a).

### Examples of non-monotonic neural responses

**Supplementary Figure 5.**
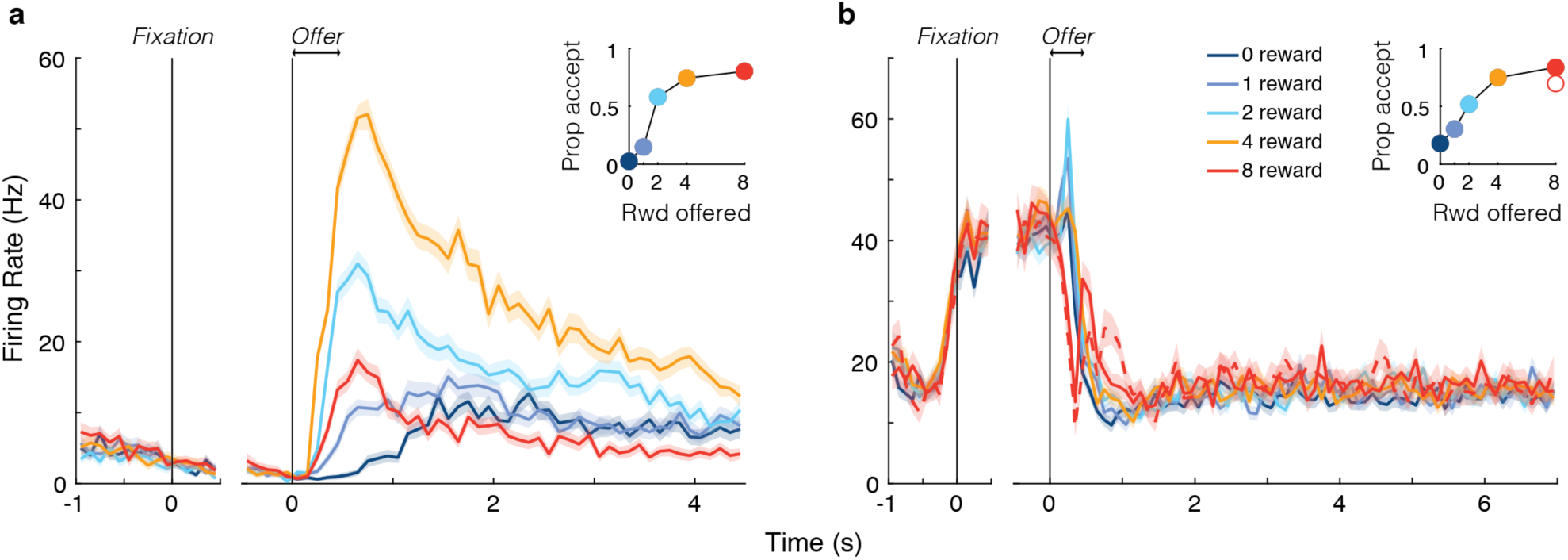
Non-monotonic encoding of offer size. (a,b) Mean (solid curves) and s.e.m. (shading) of the responses of example units from monkeys N (a) and K (b) to the various offer sizes (see legend), presented as yellow icons, as a function of time from either acquiring fixation (left panels) or onset of the offer period (right panels). Response to the singleton offer (single purple icon worth 8 juice drops) is shown by dashed curve (b only). These example units showed non-monotonic encoding of the offer size, i.e., firing the most for middle offers. This encoding was *not* due to distorted behavioral valuation of the offers, as confirmed by monotonic behavioral preference during the specific recording sessions (see inset showing proportion of offers accepted as function of offer size; open circle shows accept rate for singleton offer). These monotonic responses comprised a small proportion (∼10%) of all offer-sensitive responses, which were dominated by monotonically increasing or decreasing profiles (as in main text examples; Figure 2). We did not select units based on response profile, as uninformative units would be inherently deemphasized (i.e., smaller beta coefficient) in oTDR.

### Properties of individual-unit contributions to low-dimensional representations

**Supplementary Figure 6.**
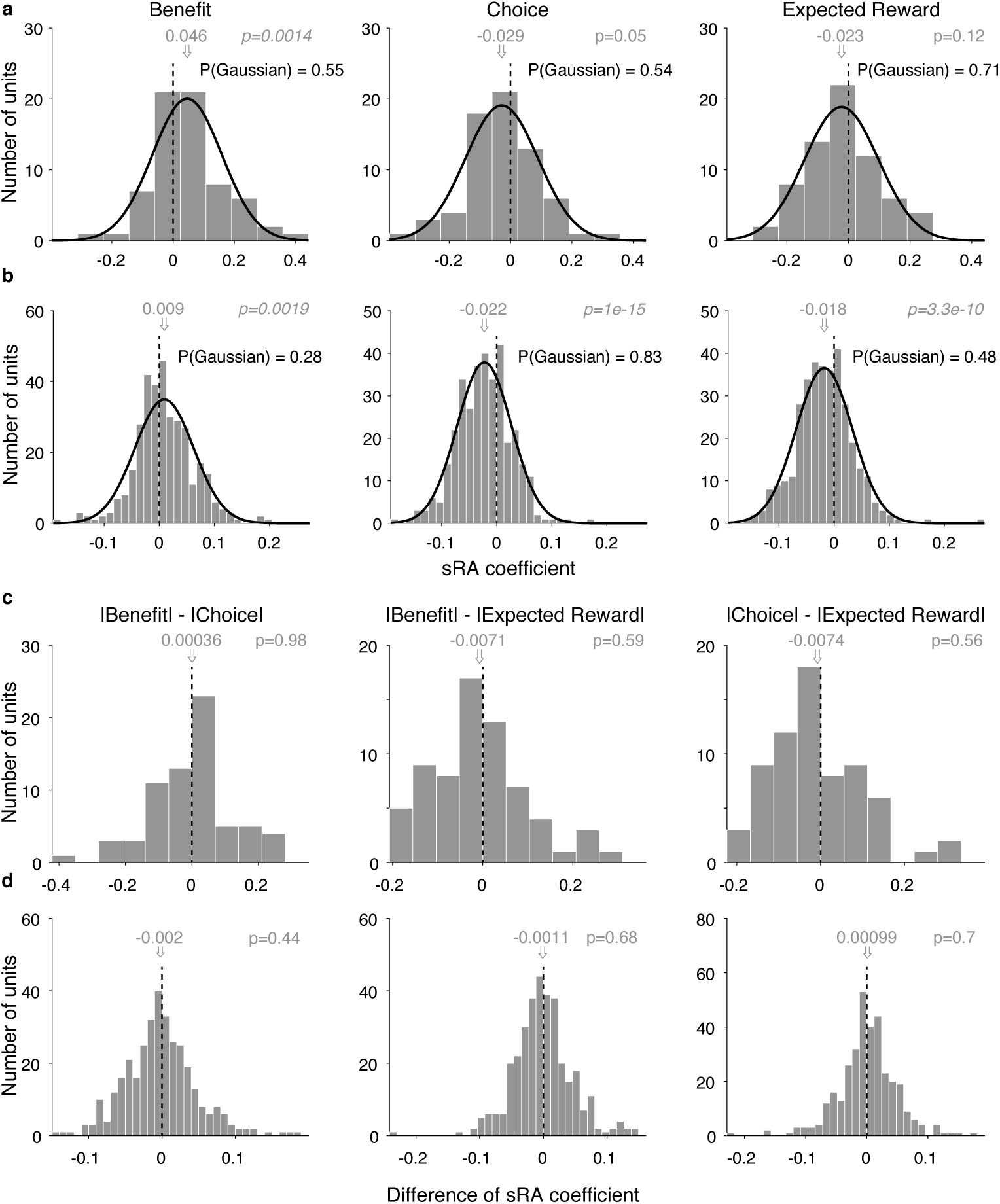
Contribution of each unit to low-dimensional representations. **(a,b)** Histogram of regression coefficients across the population that specified each unit’s contribution to the static low-dimensional representations (sRAs) of BENEFIT (left panel), CHOICE (middle panel), and EXPECTED REWARD (right panel) for monkeys N (a) and K (b). Distribution mean (gray arrow and text) and p-value (gray text) from two-tailed *t*-test of null hypothesis that mean = 0 (vertical dashed line) are shown. Gaussian functions were fit to each distribution (black curves). Probability (black text) of null hypothesis that observed distribution was Gaussian was computed via two-tailed Kolmogorov–Smirnov test. No distribution differed significantly from Gaussian. **(c,d)** Histograms of difference between absolute value of regression coefficients for pairs of representations (|BENEFIT| - |CHOICE|, left; |BENEFIT| - |EXPECTED REWARD|; middle; |CHOICE| - |EXPECTED REWARD|, right) are shown for monkeys N (c) and K (d). All conventions as in (a,b). No distribution of differences differed significantly from zero, i.e., the absolute strength of encoding did not differ for linear (i.e., benefit and choice) vs. non-linear (i.e., expected reward) variables, as observed in other prefrontal areas^13^ but not in parietal cortex, where linear terms were represented preferentially^33^. The encoding of linear vs. non-linear variables maps directly onto the concepts of linear vs. non-linear mixed selectivity, which confer distinct advantages for population decoding^12^. For present figure, all regression coefficients were found without orthogonalizing the sRAs.

**Supplementary Figure 7.**
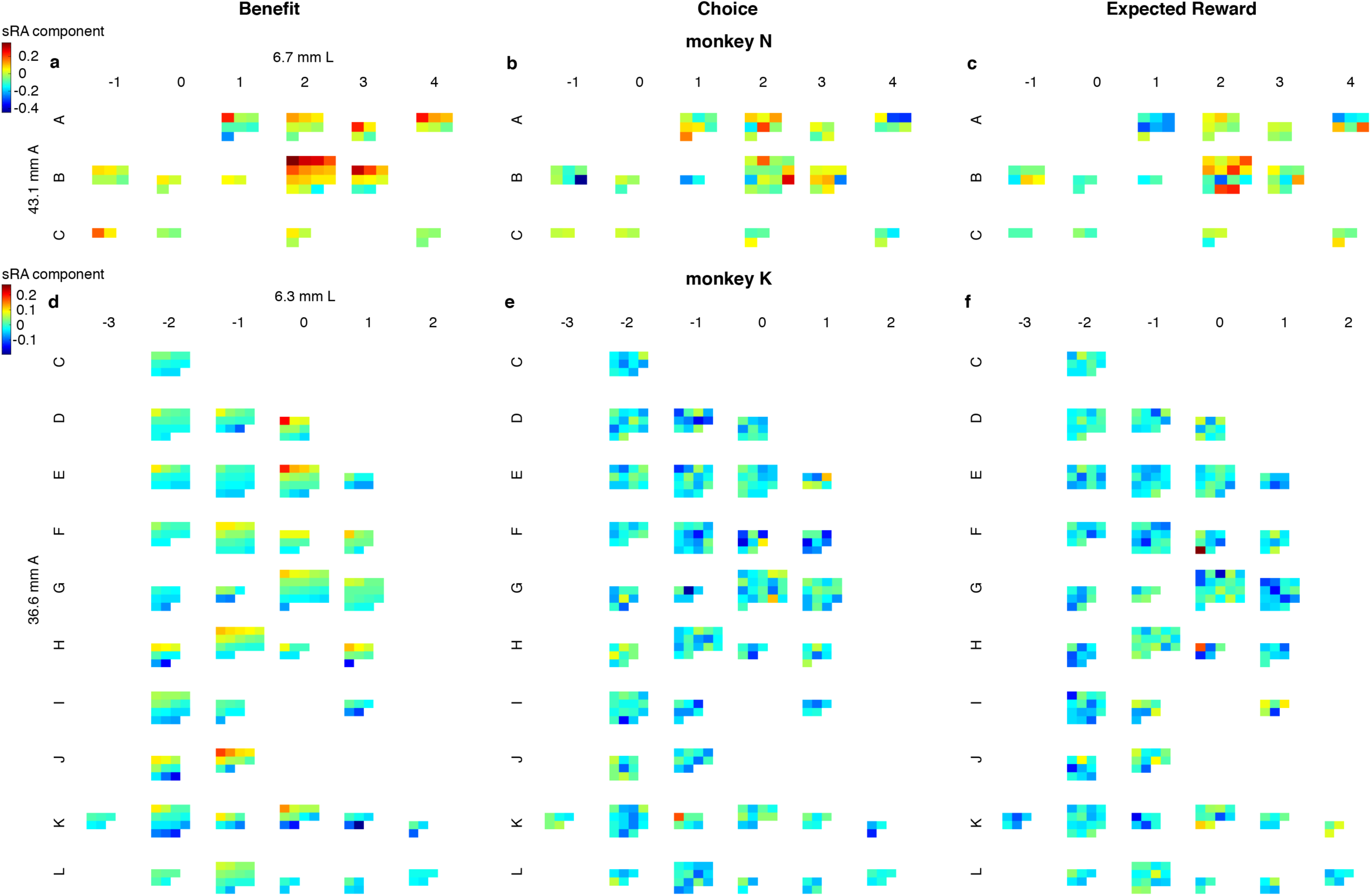
Anatomical distribution of sRA coefficients. The contribution of each unit to the low-dimensional representations (sRAs) of BENEFIT (a,d), CHOICE (b,e), and EXPECTED REWARD (c,f) are shown as a function of anatomical recording site for monkeys N (a-c) and K (d-f). Each pixel corresponds to an individual unit and the pixel color (referencing animal-specific color scale at left) corresponds to the value of the regression coefficient of the corresponding variable, i.e., the contribution of that unit to the corresponding sRA. Units (pixels) recorded from the same recording grid position (irrespective of electrode depth) are grouped together and ordered (left to right, then top to bottom) based on value of the benefit coefficient for the given unit from largest to smallest; the same order is preserved for the other representations. Blocks are arranged by corresponding 1 mm × 1 mm recording grid position from anterior to posterior (top to bottom rows) and medial to lateral (left to right columns). The grid row (letter) and column (number) labels are arbitrary. In panels (a,d), the coordinates of the grid position referenced in Supplementary Figure 4 are labeled in mm anterior to the interaural line and left of midline. Note that the anterior-posterior coordinate was measured stereotaxically in reference to the center of the recording cylinder and at the level of the recording grid, ∼30mm superior to the recording sites. Therefore, the AP coordinates of the actual recording sites may differ given small deviations in electrode trajectory in the coronal and/or sagittal planes. In contrast, the left-right coordinate was measured in the MR images at the level of the recording site.

**Supplementary Figure 8.**
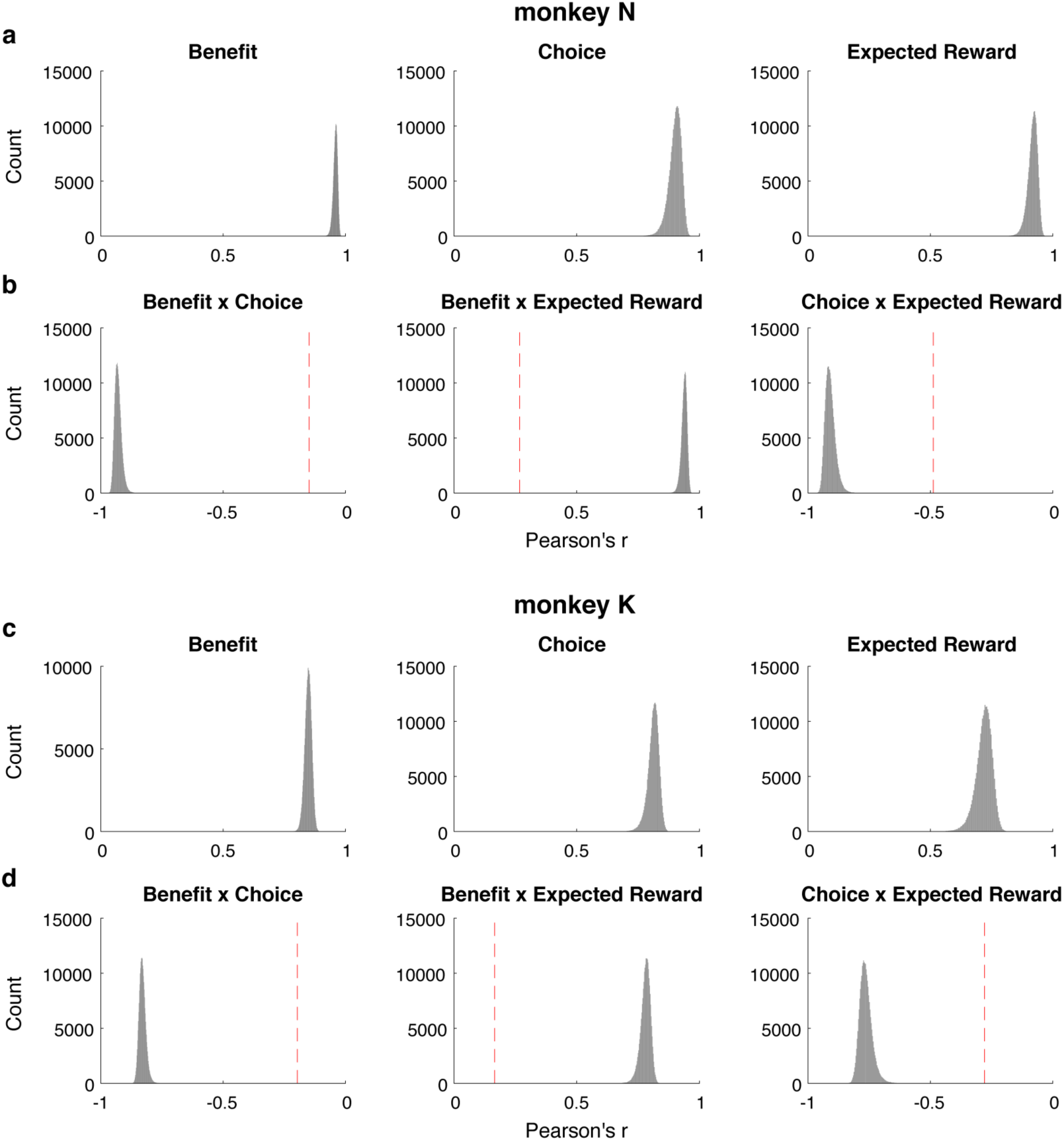
Reliability of and correlation between low-dimensional representations. We generated *S = 700* sampling datasets by randomly selecting *Q* of *Q* trials with replacement for each unit, and discovering the trio of non-orthogonalized sRAs independently for each dataset (Methods). To estimate the reliability of the sRAs, we computed the Pearson’s correlation 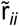 between sRAs for variable *i* (*i* ∈ {BENEFIT, CHOICE, EXPECTED REWARD}) from all pairs of sampling datasets (i.e., for S datasets, we measured (*S*^2^ - *S*)/2 pairs per variable). **(a,c)** Historgrams of 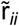 revealed very high reliability of the sRAs (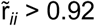 or 0.75 for monkey N (a) or K (c), respectively; Supplementary Table 2). We were interested in the separability between representations of distinct variables *i* and *j* (*i,j* ∈ {BENEFIT, CHOICE, EXPECTED REWARD}). We defined representations of *i* and *j* as separable when the correlation r_*ij*_ between sRAs for variables *i* and *j* was less than expected by chance given a null model in which representations of *i* and *j* were perfectly correlated (or anticorrelated, i.e., |r| = 1), but subject to imperfect reliability (i.e., independent noise), as estimated by 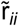 and 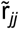, thus resulting in an observed absolute correlation of less than unity. We defined the null model empirically as 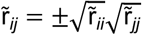 (see Methods) and took only the positive values. For display purposes only, we multiplied 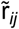 by the sign *z* of the observed correlation r_*ij*_ (all statistical tests were performed on |r_*ij*_| and 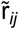). **(b,d)** Histograms of the null model 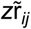 for monkeys N (b) and K (d) are shown for each pair of variables *i* and *j* and compared to the observed correlation r_*ij*_ (vertical dashed red line). For all pairs of variables, the between-variable correlations were found to be highly separable (i.e., 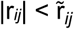 via 1-tailed t-test, p < 10^−16^; Supplementary Table 2).

### Activity of low-dimensional representations

**Supplementary Figure 9.**
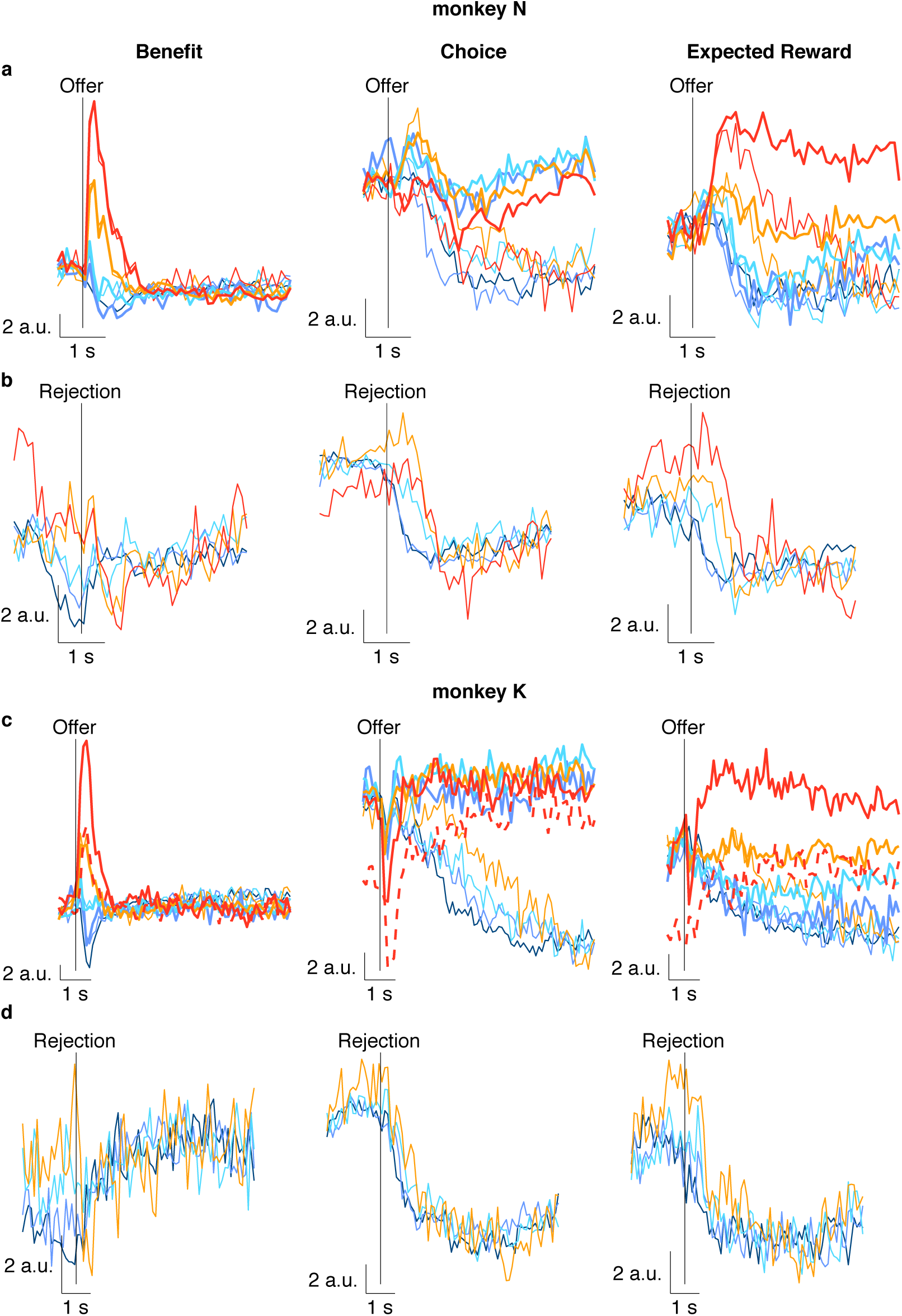
Activity of low-dimensional representations aligned to offer and rejection. Neural responses 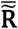 (which included the common-condition response; see Methods) were aligned to the time of the offer **(a,c)** or rejection **(b,d)** for monkeys N (a,b) and K (c,d) and projected onto the low-dimensional representations (sRAs) of BENEFIT (left panels), CHOICE (middle panels), and EXPECTED REWARD (right panels). The response for each combination of offer size (colors as in Figure 1) and choice (accept or reject choices in thick or thin curves, respectively) is shown as a function of time. We examined these projections onto the sRAs (i.e., sRA “activity”) so as to interpret the gradual separation of CHOICE activity for accept and reject choices seen in Figure 4a,b. The slow dynamics could arise from underlying single-trial responses with similarly slow dynamics (i.e., “ramps”), or from responses with rapid changes in activity (i.e., “steps”), but with variable temporal offset from the offer time proportional to variability in the rejection time across trials. By including the common-condition response and aligning to the offer (a,c), we again observed gradual separation of accept and reject responses for both animals. However, for monkey N, we now observed a rapid change in CHOICE activity that preceded the separation and therefore was common to accept and reject choices. In addition, most of the subsequent gradual separation was due to the slow recovery of the accept response, not the reject response as would be predicted by temporally offset steps aligned to the rejection. By aligning to the rejection (b,d), we observed a rapid change in the reject response after the rejection; however, this change was comparable in rate and magnitude to the offer-aligned change shared by accept responses, thus suggesting the rapid rejection-aligned dynamics were explainable by a common-condition response to the offer (i.e., common to accept and reject choices) and not a rapid step response to the rejection. In contrast, for monkey K, we observed that the offer-aligned separation of accept and reject responses was primarily driven by a gradual change in the reject response (c). When aligning to the rejection (d), the dynamics of the reject response appeared qualitatively more rapid than when aligned to the offer, suggesting that the separation of accept and reject responses was likely driven by a rapid change in neural activity on reject trials that itself was likely temporally aligned to the time of the rejection. Taken together, the separation of CHOICE activity into accept and reject responses may derive from different sources for the two animals: a slowly evolving response to accept choices (i.e., ramp) in monkey N, whereas a rapid response to the rejection (i.e., step) in monkey K. In addition, we considered the apparent “bleed through” of offer size information onto the CHOICE sRA when aligning to the offer, particularly for reject choices (a,c, middle panels). Notably, the choice selectivity appeared increasingly offset in time for larger offers. As the median rejection times were later for larger offers, this suggested that the apparent offer-selectivity was merely choice-selectivity arising at systematically later delays for larger offers. We tested this prediction by aligning to the time of rejection (b,d, middle panels), which would eliminate differential responses to offer size that were merely due to different rejection times. For monkey K, we observed no sensitivity to offer size (d), consistent with our prediction. However, for monkey N, the rejection-aligned responses showed *less* sensitivity to offer size than when aligned to the offer (b vs. a), but were still somewhat sensitive to offer, implying some overlap in the dimensions that represented benefit and choice mid-trial—overlap we examined specifically in Supplementary Figure 26. Finally, we considered the contribution of post-rejection gaze to differential accept vs. reject CHOICE activity. For monkey N, the fact that the choice discrimination was primarily due to changes from baseline on *accept* trials (a, middle panel, thick lines) suggested that gaze on *reject* trials played little role in differential CHOICE activity. Likewise, the lack of evidence for a post-rejection (i.e., post-saccadic) transient in monkey N, which might be expected if reject responses were highly gaze dependent, further argued against gaze-related activity driving differential CHOICE responses. However, for monkey K, the choice discrimination appeared to be driven by the reject responses and evidence for a post-rejection transient was stronger (discussed above); thus we could not exclude the role of post-rejection gaze in monkey K.

**Supplementary Figure 10.**
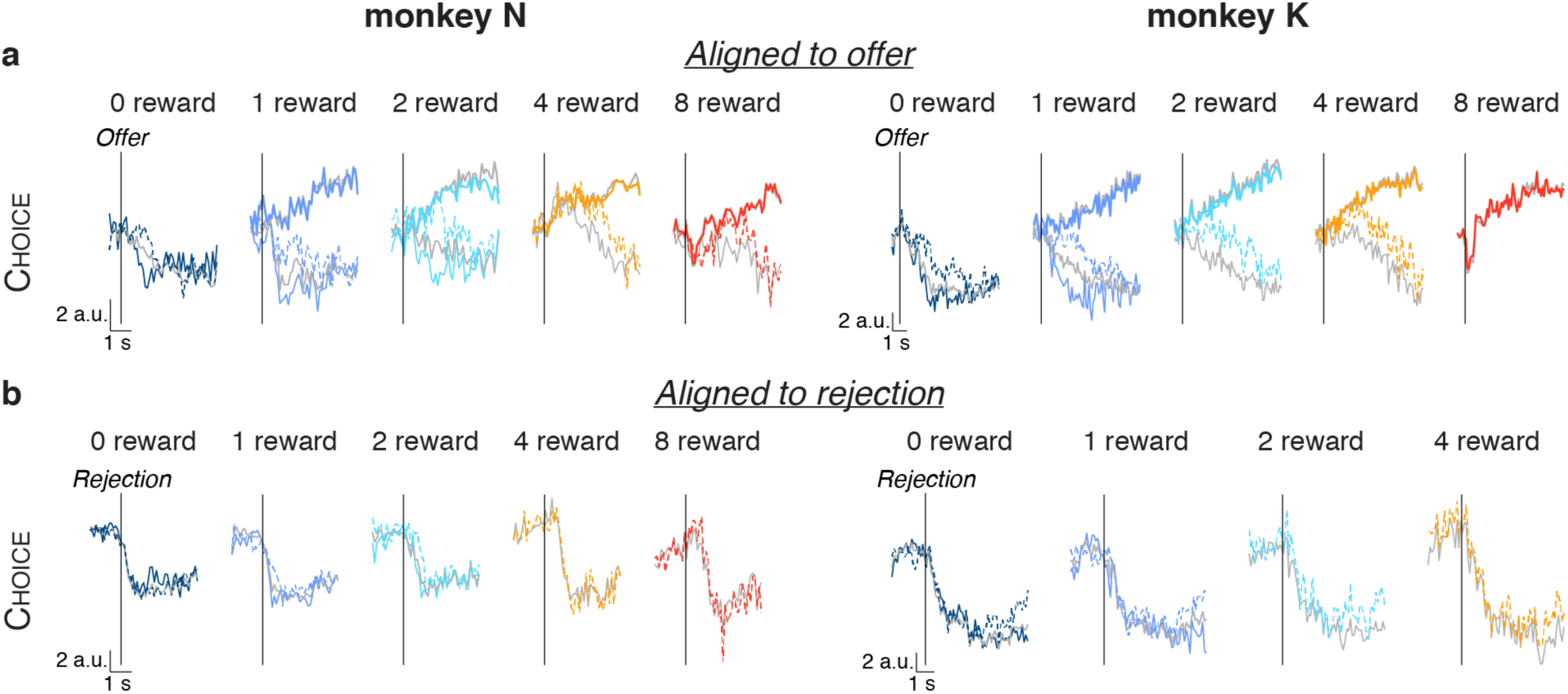
Relating neural dynamics to decision dynamics. Here we relate the low-dimensional neural dynamics to the trial-level decision dynamics. Specifically, we reasoned that if the CHOICE sRA reflected the animal’s decision process, then the time at which it discriminated the animal’s choice should depend on the time at which the choice was rendered. **(a,b)** The time course of the high-dimensional population activity projected onto the CHOICE sRA (i.e., CHOICE ‘activity’) is shown on separate panels for each offer size (colors as in Figure 1) and separated by accept and reject choices (thick and thin curves, respectively) for monkeys N (left panels) and K (right panels). Reject choices are further separated by ‘early’ and ‘late’ rejections (solid and dashed curves, respectively), defined as occurring before or after, respectively, the median rejection time across all offers for each animal (0.92 or 1.3 s, monkey N or K). In **(a)**, the responses are aligned to the onset of the offer and mean-subtracted (as in Figure 4a,b); in **(b)**, the responses are aligned to the rejection but are *not* mean-subtracted (as in Supplementary Figure 9b,d). Colored curves reflect activity of the subset of units with adequate trial counts for the present analysis (see below). For reference, the projections for *all* accept and reject trials from the full population are shown in thick and thin gray lines, respectively (identical to middle panels in Figure 4a,b and Supplementary Figure 9b,d). Aligning to the offer (a), the CHOICE activity diverged from accept choices (thick curves) at a later time for late than early rejections (dashed vs. solid color thin curves). For offers with too few early rejections to be observed separately, we compared late rejections and *all* rejections (dashed color vs. solid gray thin curves), which also showed a delayed divergence for later rejection times. This sensitivity of CHOICE activity on rejection time is consistent with our proposal that the sRA dynamics reflected the underlying decision dynamics. When aligning to the rejection (b), the CHOICE activity was virtually identical for early vs. late rejections both before and after the rejection, further supporting the thesis that the CHOICE sRA reflected the binary state of the animal’s decision at the time of the choice, independent of the time elapsed since the offer. As discussed in Supplementary Figure 9, we could not definitely exclude the contribution of post-rejection gaze to the differential CHOICE activity. However, we observed that the breaking eye movements were smaller for later rejections and for larger offers (not shown), consistent with accidental fixation breaks (see Figure 1 and related main text). Despite these overt post-rejection behavioral differences between early vs. late rejections, they did not manifest as difference in the post-rejection CHOICE activity for early vs. late rejections, further reducing the likelihood that the CHOICE sRA reflected an indirect consequence of the decision, rather than the decision itself. The activity of the BENEFIT and EXPECTED REWARD sRAs is not shown for simplicity. In summary, activity of the BENEFIT sRA was virtually identical for early- and late-rejections. Though rejection time may have reflected the animal’s initial valuation of the offer (i.e., presumably later rejections reflected higher initial valuation), any differences in initial valuation were not reflected in the BENEFIT sRA activity, consistent with the lack of a choice predictive, post-offer signal in the single-neuron analysis (Supplementary Figure 12). The impact of rejection timing on the activity of the EXPECTED REWARD sRA was equivocal. However, to the extent that activity for late-rejection trials differed from either early- or all rejection trials, the late-rejection activity tended to hew more closely to the activity on the corresponding accept trials. This trend was consistent with EXPECTED REWARD encoding the animal’s real-time valuation that, in turn, informed how likely the animal maintains its initial behavioral policy to accept the offer.

#### Condition and unit selection

As in the main text, certain conditions—now defined by offer, choice, *and* rejection time—had too few trials per unit to accurately estimate trial-average responses and were excluded. We required that a given condition have at least 5 trials per unit for that condition to be included, as in the main text. To preserve as many conditions as possible in the present analysis, we allowed for conditions that did not meet this minimum trial count for *all* units, but did meet the criterion for at least 85% of units. As such, we eliminated 2 of the original 9 conditions for monkey N (early rejections of 4- and 8-reward offers) and 2 of the original 8 conditions for monkey K (early rejections of 2- and 4-reward offers). (Recall that some conditions had already been eliminated for insufficient trial count in the main analysis.) Units not meeting the minimum trial count for one or more of the included conditions were eliminated: 16 of 68 units (24%) for monkey N and 73 of 342 units (21%) for monkey K.

To allow for comparison with the main text, we preserved the CHOICE sRA coefficients from the main text. However, because we eliminated certain units in the present analysis, we effectively eliminated the corresponding dimensions from the sRA. Since this step necessarily reduced the magnitude of the new projections, we compared the accept trials between the pared-down and full populations (color vs. gray *thick* curves) and observed only a minor decrement in magnitude. This decrement was markedly smaller than the differences between all rejections in the full population (gray *thin* curves) and than either the early or late rejections in the pared-down population (color *thin* curves), meaning these differences were unlikely due to the changes in population membership. Note that the principal comparisons between accept choices, early rejections, and late rejections (colored curves), discussed above, were all within the pared-down population, and thus were not attributable to changes in population membership.

**Supplementary Figure 11.**
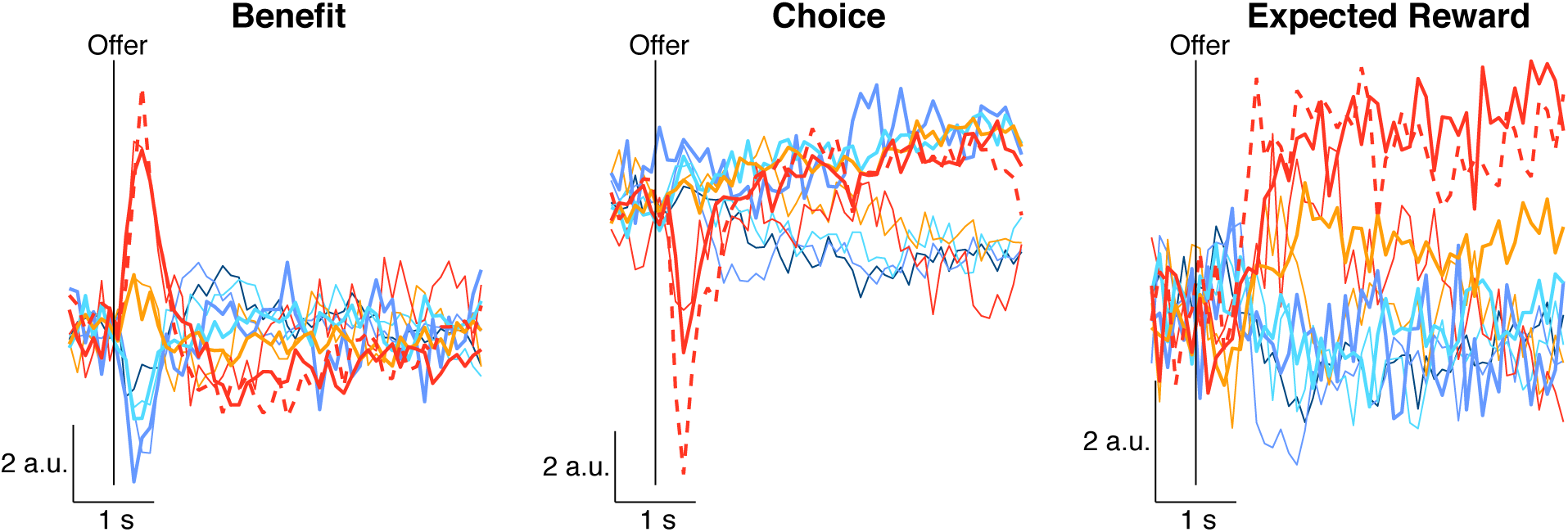
Activity of low-dimensional representations including singleton offer for monkey N. We separately discovered the low-dimensional representations (sRAs) in a subpopulation limited to units recorded with the singleton offer for monkey N. (For monkey K, all units were recorded with the singleton offer, as shown in the main text). As for monkey K, the singleton response was not included when computing the sRAs. We projected the population response onto the sRAs of BENEFIT (left), CHOICE (middle), and EXPECTED REWARD (right) as a function of time from the onset of the offer period (vertical black lines), with all conventions as in Figure 4a,b. The response to the singleton offer (thick dashed red curve) was comparable to the value-matched, 8-reward non-singleton offer (thick solid red curve), consistent with the OFC population encoding the value and not visual properties of the stimulus. The animal rarely rejected the singleton offer, and so too few trials existed to analyze this condition.

### Choice probability in individual units

**Supplementary Figure 12.**
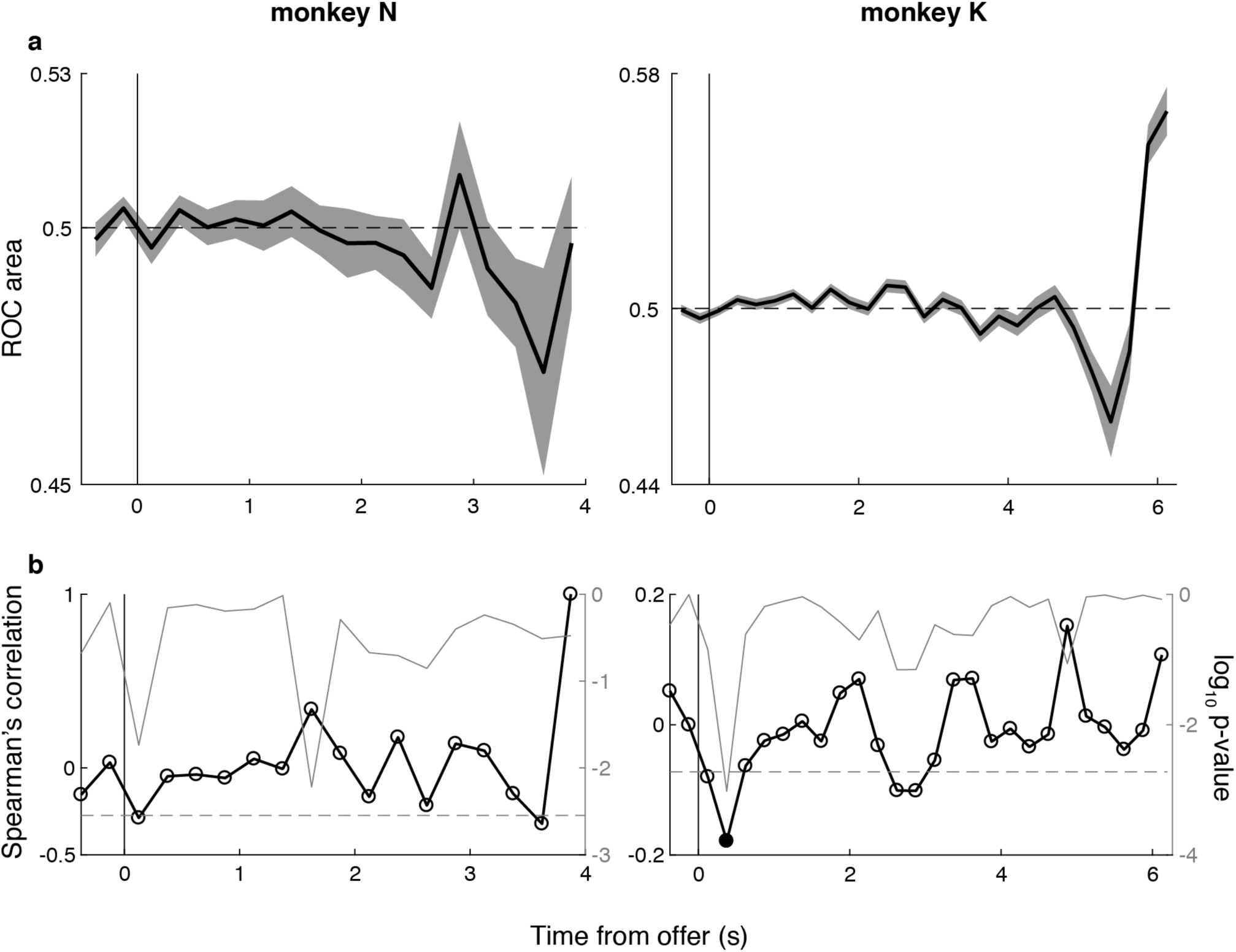
Choice probability in individual units. We reasoned that if OFC activity were used by downstream circuits to estimate the value of a given offer, and that trial-to-trial fluctuations in value accounted for variability in choice of a given offer size, then we would expect trial-to-trial variation in OFC activity to predict the animal’s choice. Choice-predictive responses have been observed in sensory areas representing the sensory stimulus in perceptual decision-making studies^48,91^. However, evidence for robust choice-predictive responses in OFC has been equivocal^15,80^. We quantified the choice predictivity in OFC using the well-established choice probability (CP) metric, as described elsewhere^48^ and in Methods. Briefly, for each unit, we extracted the neural response r(*t*,T) in 250 ms non-overlapping time bins *t* aligned to the offer presentation on each trial T. So as to isolate responses *predictive* of an upcoming choice, we excluded trials for a given time bin in which the animal had rejected the offer prior to that time bin. Therefore, later time bins necessarily included fewer trials. To control for the known influence of offer size on the neural response, we z-transformed r(*t*,T) within each offer size (provided at least two trials were observed for a given offer and time bin) and combined responses across offers to give the normalized response R(t,T). We stratified R(*t*,T) according to the choice on trial T and compared the distributions of responses on accept vs. reject trials at each time bin *t*, which we quantified as the area under the receiver operator characteristic (ROC) curve, or equivalently, CP(*t*)—the probability of an ideal observer correctly classifying a trial as an accept choice given the neural response^92^. We eliminated time bins with fewer than 10 trials (across offers) contributing to the accept or reject distribution. CP(*t*) = 0.5 indicated chance performance. Values significantly greater or less than 0.5 indicated the response was choice predictive, with higher firing rates predicting accept or reject choices, respectively (significance defined below). **(a)** The median CP(*t*) across units (i.e., ROC area; thick black curve) is shown as a function of time from the onset of the offer period for monkeys N and K (left and right panels, respectively). Gray shading indicates +/- the median absolute difference (i.e., median(|CP(*t*) – median(CP(*t*))|)). Note that the curves do not extend the full length of the trial because the later time bins had too few trials to include (i.e., most rejections had occurred by this time). At no time was CP(*t*) significantly different from chance (i.e., p(CP = 0.5) > 0.05 for all time bins) by Wilcoxon signed rank test of null hypothesis that median CP = 0.5 (horizontal dashed-line), Sidak-corrected for multiple time bins. The computation of CP required analysis of single-trial data so as to isolate responses occurring before the time of rejection, a time which varied across trials. In contrast, all population-level analyses, such as oTDR, depended on trial-average data so as to pool across serially recorded units. Thus, the choice predictive analyses were limited to individual units. In an effort to relate the above individual-unit analysis to the population-level analyses discussed in the main text, we compared CP(t) to the contribution individual units made to the population low-dimensional representations (sRAs). Specifically, we reasoned that units representing variation in value across offer sizes (as measured by their contribution to the BENEFIT sRA) may have represented trial-to-trial fluctuations in value within an offer size (as measured by CP), a relationship that was observed for units in area MT: units encoding visual motion direction across different levels of motion strength also predicted the subject’s perceptual report within a given level of motion strength^49^. In addition, this hypothesis would address a second limitation of averaging CP across units. Namely, averaging assumed a common sign of choice predictive encoding (e.g., firing more before accept vs. reject choices), whereas the sign of encoding may have been heterogeneous across the population, which would be obscured by simple averaging. By relating CP to the encoding of offer size, we inferred the sign of within-offer value encoding from the sign of between-offer value encoding. **(b)** Specifically, we computed the Spearman’s correlation coefficient (thick black curve and open circles, referencing left ordinate) between CP(*t*) for a given unit and the BENEFIT sRA coefficient corresponding to that unit as a function of time *t* from the onset of the offer period for monkeys N and K (left and right panels, respectively). The log_10_ probability p of observing the correlation coefficient by chance (thin gray curve) is plotted on the same panels and references the right ordinate. We marked as significant (filled circles) those time bins when p < 0.05 (Sidak-corrected for multiple comparisons across time bins, as above; corrected p-threshold shown as gray horizontal dashed line). At no time was a significant correlation observed for monkey N, and only a single time bin was significant for monkey K (filled circle). In summary, individual units in OFC did not demonstrate choice predictivity. That is, on average, trial-to-trial variation in firing rate was not systematically related to trial-to-trial variation in choice. In addition, for a given unit, we did not observe a general relationship between its putative choice predictive signal (CP) and its contribution to the population’s encoding of benefit. We observed a single instance of a relationship between CP and BENEFIT (single time bin, monkey K only), which was only modestly more likely than chance. To address the question of choice predictivity in OFC, simultaneous recording of many units would be required to observe single-trial choice representations at the level of the population.

### Sensitivity and specificity of the low-dimensional representations

**Supplementary Figure 13.**
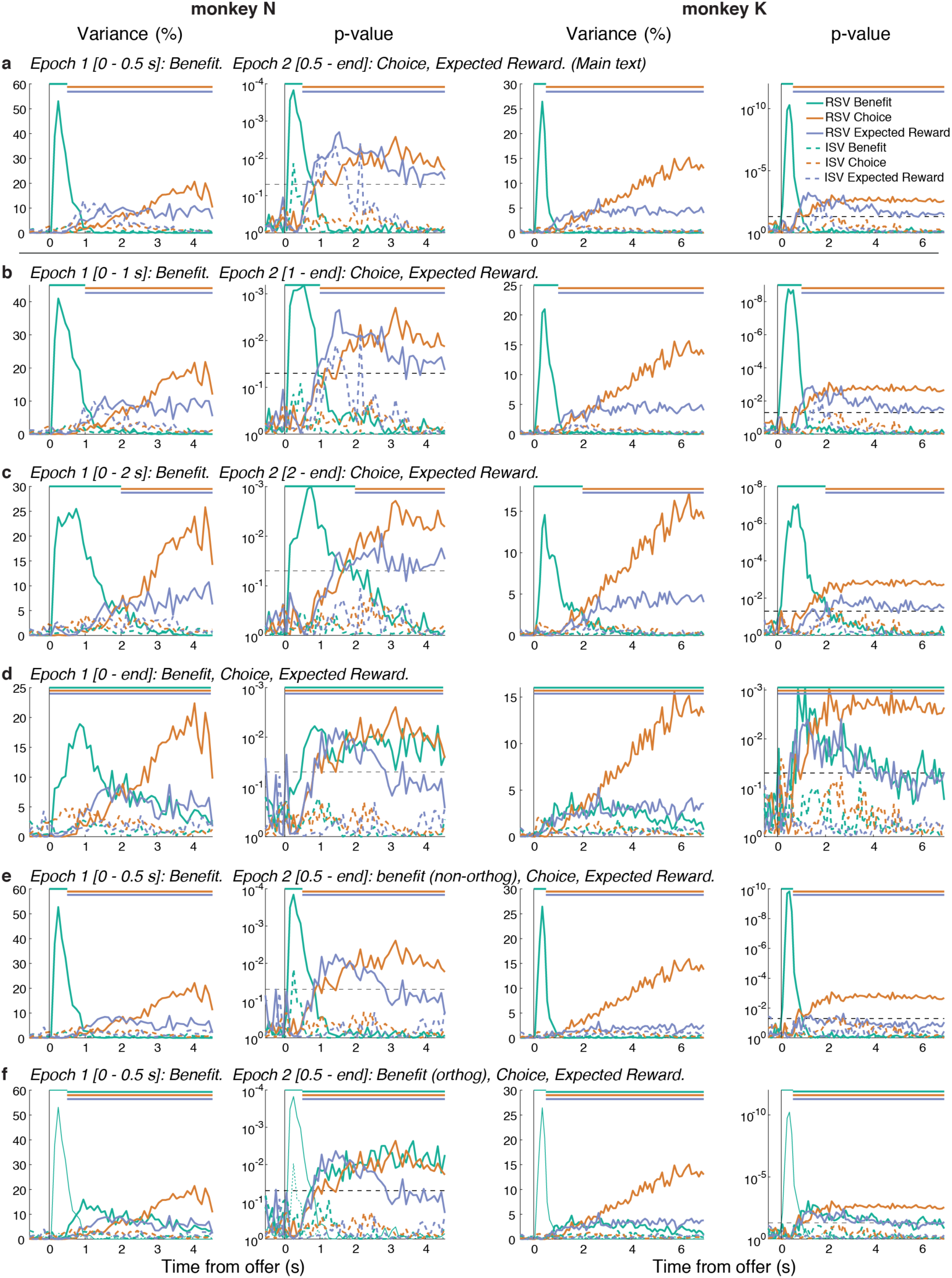
Effect of temporal epoch on the sRAs. **(a-f)** The relevant and irrelevant signal variance (RSV and ISV, solid and dashed curves, respectively) are shown in columns 1 and 3, with associated p-values in columns 2 and 4, for BENEFIT (green), CHOICE (orange), and EXPECTED REWARD (blue) as a function of time from the onset of the offer period for monkeys N (columns 1-2) and K (columns 3-4). Each row shows a separate implementation of oTDR with distinct temporal epochs for computing the sRAs (italicized headings). The colored horizontal bars at the top of each panel indicate the temporal epoch in which the color-matched sRA was computed. **(a)** The oTDR implementation used in the main text is shown, with BENEFIT computed in Epoch 1 (corresponding to the offer period, 0 – 0.5 s) and CHOICE and EXPECTED REWARD computed in Epoch 2 (corresponding to the work period, 0.5 s to end of the trial). The sRAs explained significantly large portions of relevant variance throughout the epoch in which they were computed and little variance outside that epoch (solid curves). In addition, the ISV (i.e., variance unrelated to the targeted variables; dashed curves) was generally very small and statistically insignificant; this was true across the various implementations of oTDR and is not further discussed. **(b,c)** We tested the impact on sRA sensitivity of extending the duration of Epoch 1 to either 1 s (b) or 2 s (c), while shortening Epoch 2 accordingly. Extending Epoch 1 reduced the RSV explained by BENEFIT (green curves) and, particularly for the 2 s duration (c), marginally increased the RSV for CHOICE (orange curves). These changes were predicted from the dRA similarity analysis: the encoding of benefit was stable during the 0.5 s offer period, then changed abruptly (Figure 5c-f, left panels). Therefore, targeting an sRA exclusively to the 0.5 s offer period would more fully capture the available signal, whereas extending the epoch in which the sRA was computed would necessarily dilute the specificity of the resulting sRA for the time-limited representation, as the sRA would now capture a compromise between two, highly dissimilar representations (corresponding to the periods marked with red and blue brackets in Figure 5e,f, left panels). Likewise, because the representation of choice did not emerge until ∼1.5 s (Figure 5c-f, middle panels), targeting the CHOICE sRA exclusively to this temporal epoch (as in (c)) resulted in a (marginally) greater RSV. Nonetheless, in the case of the choice signal, the representation was highly stable for an extended period and was without a competing representation prior to this period. Thus the CHOICE sRA suffered very little by including, as in the main analysis, the roughly 1 s period (0.5 s ∼ 1.5 s) prior to the emergence of choice encoding. Despite these various quantitative effects of extending Epoch 1, the qualitative conclusions remained unchanged: the static sRAs of BENEFIT, CHOICE, and EXPECTED REWARD captured a high and statistically significant portion of variance related to their respective signals throughout the temporal epochs in which they were computed. **(d)** Next we considered the case of having no *a priori* assumptions about the timing or stability of the task-relevant representations. In this implementation, we computed all three sRAs within a single epoch spanning the entire trial. As expected, the maximum explanatory power of the BENEFIT sRA decreased, but remained significant and spanned a longer portion of the trial, consistent with pooling over different representations of benefit during different phases of the trial, as discussed above. The RSV for CHOICE was largely unaffected by use of a single temporal epoch, showing how oTDR is generally robust to inclusion of non-coding activity (i.e., before 1.5 s). However, the RSV for EXPECTED REWARD was diminished, particularly in the last 1 ∼ 2 s of the trial, where it dropped below the significance threshold. Note that in the single-epoch model, EXPECTED REWARD was forced to compete with BENEFIT to explain variance during a time when the two signals were correlated (see Supplementary Figure 26a,b, middle panels, showing similar dRAs for benefit and expected reward late in the trial). Therefore, the decrease in RSV for EXPECTED REWARD was likely due to the inclusion of the temporally overlapping benefit regressor during this later period, rather than due to the expansion of the epoch in which EXPECTED REWARD was computed (i.e., starting at 0 s instead of 0.5 s). This conclusion was further supported by the next two analyses: including a separate benefit regressor in Epoch 2, while maintaining the original duration of Epoch 2, caused a decrease in RSV for EXPECTED REWARD compared to the original analysis. Based on the single-epoch results (d), we advocate for a temporally targeted implementation of oTDR either when motivated theoretically and/or when suggested by the dRA analysis (both of which are true in the present dataset). Nonetheless, even with the least constrained assumptions, oTDR discovered statistically sensitive and specific representations of the task-relevant variables. Thus far, we have specified oTDR to assume that each task-relevant variable was represented in one and only one temporal epoch. However, in specifying the general form of the objective function (Eq. 6), one may assume that any arbitrary set of variables was encoded in one or more of any arbitrary number of temporal epochs. Moreover, one may assume that any arbitrary subset of variable-by-epoch representations were orthogonal with one another, while making no orthogonality assumptions about any remaining representations. In the final two implementations, we demonstrated this flexibility of oTDR as applied to the current dataset. As discussed, benefit was encoded by at least two distinct representations: during the offer (0 − 0.5 s) and after the offer during the work period (> 0.5 s) (also see Figure 5c-f, left panels, colored brackets). Here we use oTDR to explore this latter representation for two purposes. First, one may want to control for the overlap between benefit and expected reward encoding during Epoch 2 (as discussed above and shown in Supplementary Figure 26a,b, middle panels). In this case, one is *not* interested in the read-out of the mid-trial benefit signal, only controlling for its impact. In oTDR, we assume that sRAs are orthogonal because this guarantees that the readout (both for experimenter and downstream circuit) is independent between sRAs. However, in the case of only controlling for the mid-trial benefit signal, we do not require an sRA for mid-trial benefit and thus do not orthogonalize its representation. In **(e)**, we show the results of a model that included a non-orthogonalized regressor for benefit in Epoch 2, in addition to the orthogonalized regressors that gave rise to the sRAs from the main text (i.e., BENEFIT in Epoch 1 and CHOICE and EXPECTED REWARD in Epoch 2). (Note that because coefficients from the mid-trial benefit regressor were not orthogonalized with respect to the sRAs, any variance-related metrics, i.e., RSV and ISV, for mid-trial benefit would not be independent of the sRAs and thus were not computed or plotted.) We found that including the mid-trial benefit regressor had virtually no impact on the BENEFIT and CHOICE sRAs, but reduced the explanatory power and statistical significance of EXPECTED REWARD, particularly late in the trial for monkey N and throughout the trial for monkey K. As discussed, this is likely due to the similarity of benefit and expected reward representations during the impacted trial periods (Supplementary Figure 26a,b). By removing the orthogonality constraint, the mid-trial benefit representation would have an advantage over the EXPECTED REWARD sRA, which had to remain orthogonal with the other sRAs, in competing for shared variance. Thus, this implementation of oTDR was the most conservative in addressing the specific concern of misattribution of variance related to benefit late in the trial. Second, one may have a functional interest in the mid-trial benefit representation, such as considering how the early and mid-trial benefit representations maybe readout independently by downstream circuits. To test this hypothesis, we implemented oTDR using the same regressors as panel (e), only now we assumed that the mid-trial benefit representation *was* orthogonal to the other sRAs (and likewise discovered a dedicated MID-TRIAL BENEFIT sRA). As shown in **(f)**, this model discovered sRAs that were virtually identical to those from the previous non-orthogonalized model, except that now the RSV for EXPECTED REWARD in monkey K was both larger and significant for the first 3.5 s of the work period. This “rescue” of EXPECTED REWARD was due to the fact that the competing mid-trial benefit regressor now had to abide the orthogonality constraint along with the other sRAs. Note that this did not confer any special advantage for EXPECTED REWARD to explain this variance; that is, if the mid-trial variance were more related to offer size, then it would be explained by the MID-TRIAL BENEFIT sRA, leaving EXPECTED REWARD to explain little to no variance. Finally, we now also could observe the MID-TRIAL BENEFIT sRA (f, thick green line), which explained a significant portion of benefit information from just after the offer (0.5 s) to the middle (monkey K) or end (monkey N) of the trial, as anticipated from the dRA analysis (Figure 5c-f, left panels).

**Supplementary Figure 14.**
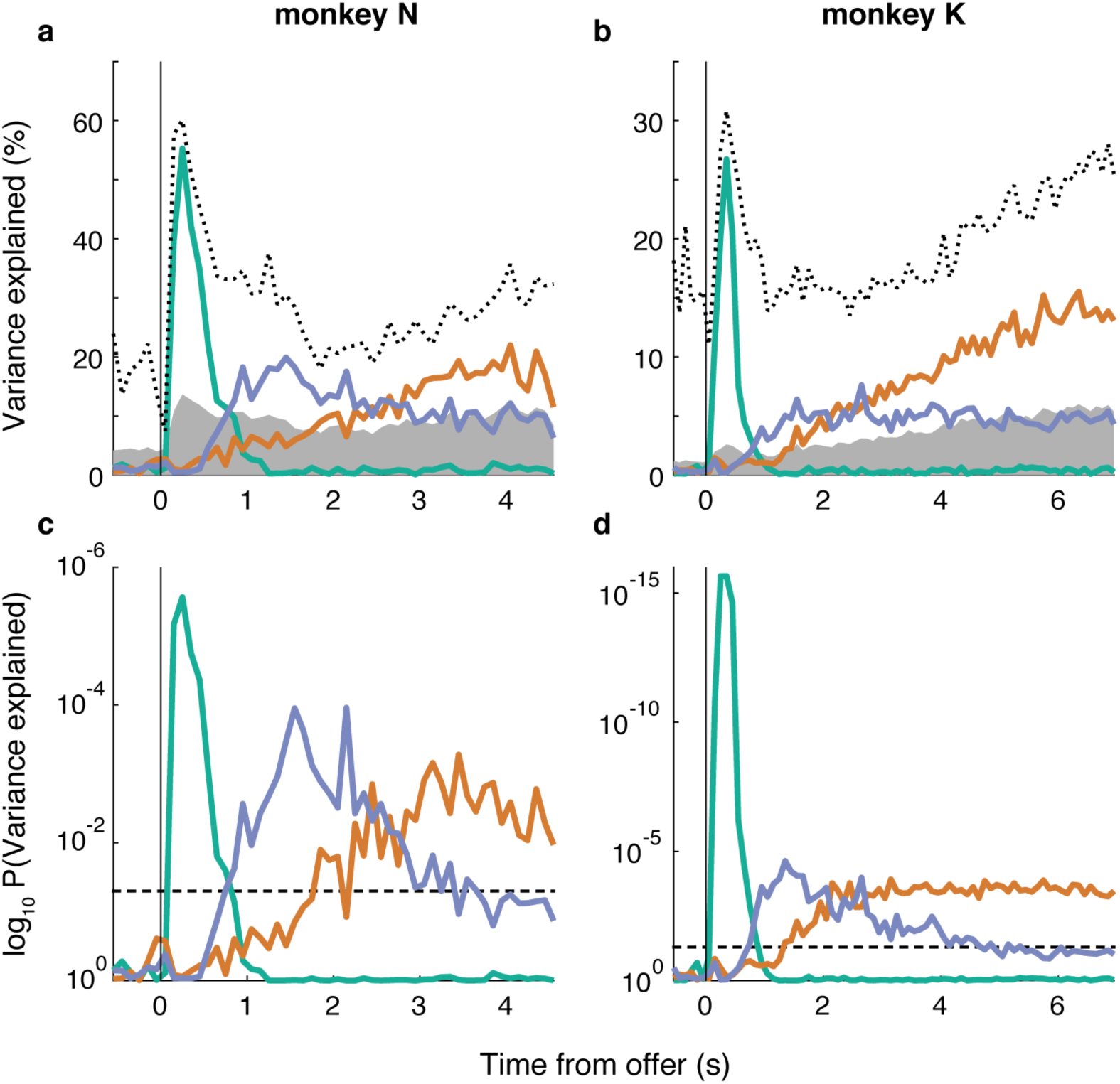
Variance explained by low-dimensional representations. **(a,b)** The percentage of variance explained (i.e., cross-condition variance of the projection onto a given sRA normalized by the cross-condition variance across all dimensions) by the sRAs of BENEFIT (green curve), CHOICE (orange curve), and EXPECTED REWARD (blue curve) is plotted as a function of time from the onset of the offer period (black vertical line) for monkeys N (a) and K (b). The area of gray shading represents the variance explained by 95% of random vectors reflecting the dimensionality of the data. To estimate the upper-bound for variance explained any single, time-varying dimension (note that the sRAs were static, in contrast), we performed principal component analysis independently at each time bin and plotted the variance explained by the top component (black dotted curve). **(c,d)** We computed the probability, P(V), of observing the percent variance explained by chance based on the distribution of variance explained by the set of random vectors. We plotted log_10_ P(V) for each sRA (colors as in a,b) as a function of time from the onset of the offer period (black vertical line) for monkeys N (c) and K (d). For monkey K, P(V) at times 0.25 and 0.35 s was less than machine precision, and the values were replaced with the precision floor value. The threshold P = 0.05 is plotted (horizontal dashed line) for reference.

**Supplementary Figure 15.**
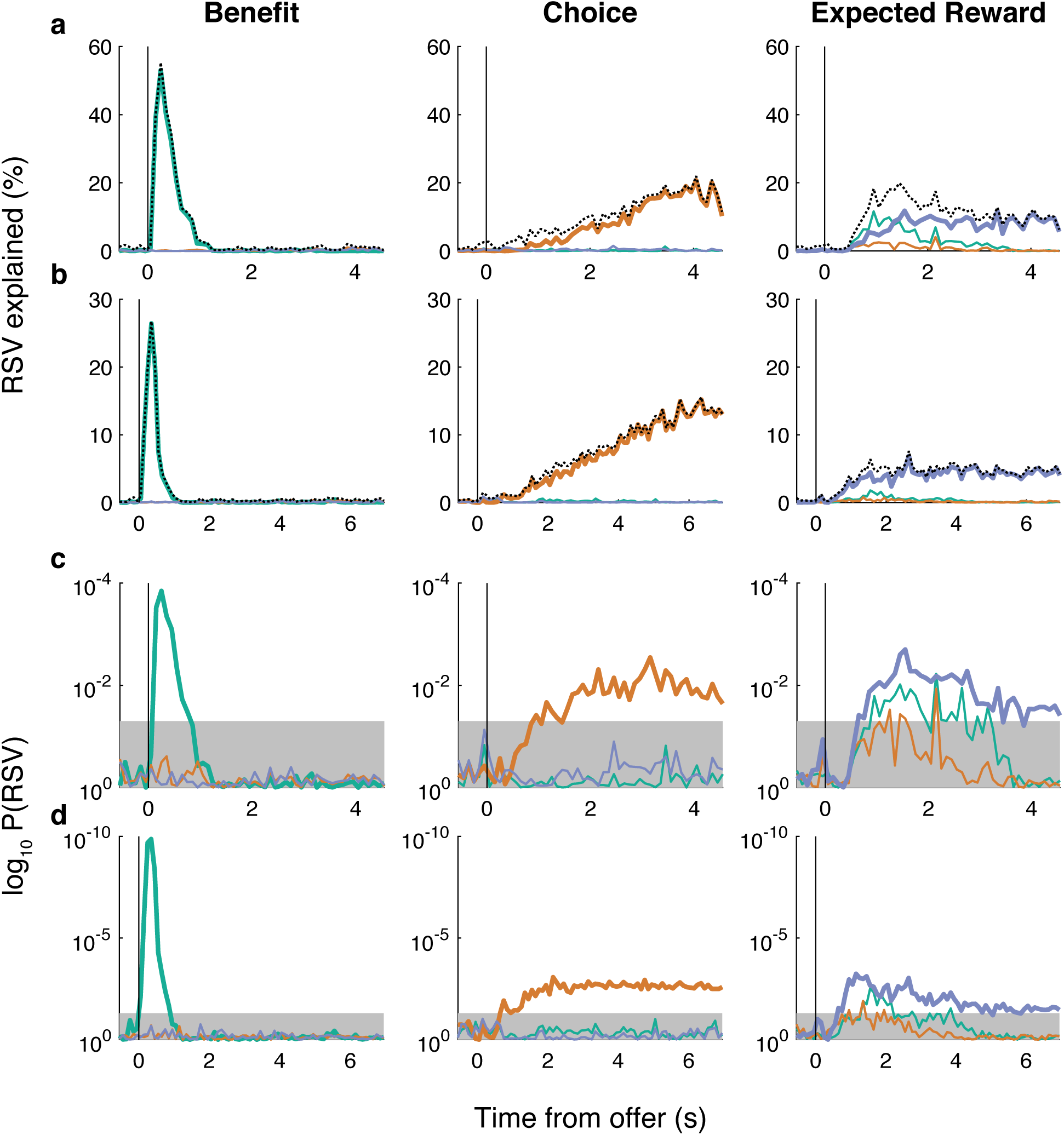
Specificity of low-dimensional representations. We were interested in the specificity of the low-dimensional representations, that is, the extent to which the sRAs explained variance related to the targeted variable of interest and not related to off-target variables. For each sRA, we computed the relevant signal variance (RSV) with respect to each variable (Methods), thereby generating 3 RSV values for each RSV at each time bin. **(a,b)** The RSV explained by the sRAs of BENEFIT (left panel), CHOICE (middle panel), and EXPECTED REWARD (right panel) is plotted (solid colored curves) as a function of time from the onset of the offer period (vertical black line) for monkeys N (a) and K (b). Colors refer to the variable with respect to which the RSV was computed: benefit (green), choice (orange), and expected reward (blue). The on-target RSV is shown in thick curves (e.g., RSV explained by BENEFIT with respect to the benefit variable is shown in left panel in thick green curve) and recapitulates Figure 4c,d. Off-target RSV is shown in thin curves (see below). The overall variance (V) explained by the sRA (dotted black curve) recapitulates the curves in Supplementary Figure 14a,b and served as an upper-bound for RSV explained. When the difference between V and RSV was zero, the sRA was perfectly specific to the on-target variable. Any lack of specificity of the sRA (i.e., irrelevant signal variance; Figure 4c,d, dashed curves) must have arisen within the difference between V and RSV. The RSV with respect to the off-target variables depended on the correlations between the variables themselves, which ranged as high as Pearson’s r = 0.86 for expected reward and benefit for monkey K. For an intuition, if the correlation between variables A and B were r = 1, then the RSV with respect to A and B would be equal. To address the correlation between variables, we used semi-partial correlation to compute the off-target RSV with respect to off-target variable *q* that would be expected given the correlation r_*kq*_ between *q* and on-target variable *k* (see Methods). The off-target RSV (thin colored curves) is plotted in (a,b), where the color indicates the off-target variable *q* with respect to which the RSV was computed (the on-target variable *k* is given by the column). **(c,d)** The probability of observing on- and off-target RSV (thick and thin curves, respectively) by chance is plotted for monkeys N (c) and K (d), where panels and colors are as in (a,b). Probabilities were derived from empirical null distributions of on- and off-target RSV (see Methods). The area of gray shading represents P(RSV) > 0.05. In general, on-target RSV closely approached the upper-bound V, indicating high-specificity of the sRAs. In addition, the probability of observing the on-target RSV by chance was less than 0.05 during key task-relevant periods (defined in main text). The off-target RSV was generally very small and below chance levels. However, for both animals, EXPECTED REWARD explained significant variance related to benefit from 1 ∼2 s after the offer (thin green curve, right panels). This “leak” of variance related to benefit onto EXPECTED REWARD suggested that the transformation from encoding of pure benefit to encoding of benefit conditioned on choice (i.e., expected reward) was not instantaneous, but rather evolved gradually during and just after the period when the animals rendered most of their choices, as can be observed in the trial-average responses projected onto EXPECTED REWARD (Figure 4a,b, bottom panels), which separate rapidly by offer size in the first second after the offer prior to additionally separating by choice (see Supplementary Figure 26 for relationship between dynamic regression axes for different variables). Of note, the lack of benefit information explained by the BENEFIT sRA during this time (thick green curve, left panels) indicated that the residual benefit information during the transformation was in a direction orthogonal to BENEFIT, consistent with the reduced sensitivity of BENEFIT during this time (see Supplementary Figure 16). The cross-talk between benefit and expected reward information was distinct from the leak of benefit information onto CHOICE mid-trial, as also observed in Figure 4a,b (middle panels) and discussed in the main text.

**Supplementary Figure 16.**
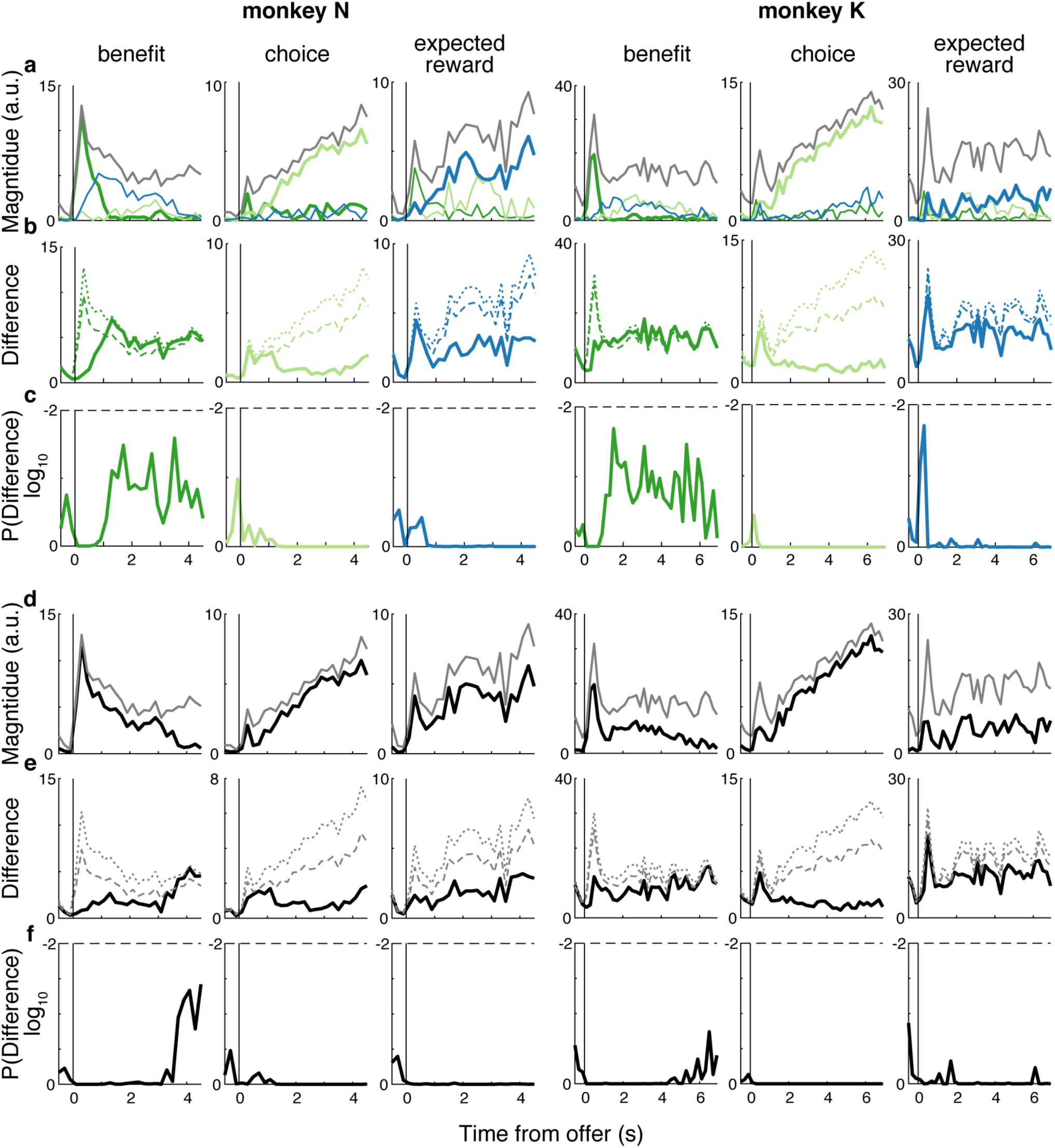
Sensitivity of low-dimensional representations. We were interested in the sensitivity of the static regression axes (sRAs) to the encoding of the task-relevant variables at the individual-unit level. That is, how much of the encoded information available in the population at large was captured by the low-dimensional, static space spanned by the sRAs? We reasoned that the dynamic regression axes (dRAs) discovered the best linear representation of the variables at each time bin, and thus could be used to quantify the available momentary information. However, the approach most analogous to our prior analyses (i.e., projecting the neural data onto the RAs and measuring the variance explained) was not possible for the dRAs because the axes were non-orthogongal, and thus the projection onto a given dRA would necessarily contain off-target variance related to the other variables. Instead, we relied on the observation that the vector magnitude ‖dRA_*k*_(*t*)‖, as defined in the high-dimensional space and prior to vector normalization, was proportional to the degree of neural representation of variable *k* at time *t* relative to the other variables and time bins (recall that predictors were scaled and centered and neural responses were mean-centered). In addition, we reasoned that the degree to which dRA_*k*_(*t*) aligned with each sRA, or with the low-dimensional space spanned by all three sRAs, provided a measure of how sensitive the sRAs were for variable *k* at time *t*. Therefore, we projected dRA_*k*_(*t*) onto each sRA_*i*_ (*i,k* ∈ {benefit, choice, expected reward}) and measured the resulting magnitude, *z*_*k,i*_(*t*). When *z*_*k,i*_(*t*) approached ‖dRA_*k*_(*t*)‖, then sRA_*i*_ captured a high proportion of the available information about variable *i* at time *t*, i.e., it was sensitive to variable *i*. We measured *z*_*k,i*_(*t*) both for the on-target variable (i.e., *i* = *k*), as well as for the off-target variables (i.e., *i* ≠ *k*), which indicated the extent to which sRA_*i*_ was sensitive to the other variables. **(a)** We plotted *z*_*k,i*_(*t*) as a function of time *t* from the onset of the offer period (vertical black line) for each dRA_*k*_(*t*). Within the left (monkey N) or right (monkey K) set of three columns, the left, middle, and right panels correspond to *k* = benefit, choice, and expected reward, respectively. Line color corresponds to the sRA_*i*_ onto which the dRA was projected (sRAs of BENEFIT, CHOICE, and EXPECTED REWARD plotted in green, orange, and blue, respectively). Thick or thin colored lines indicate the on- or off-target variable (i.e., *k* = *i* or *i* ≠ *k*), respectively. For reference, the magnitude of dRA_*k*_(*t*) in the high dimensional space—an upper-bound for *z*_*k,i*_(*t*)—is shown in gray curves. Qualitatively, it appeared that the sRAs captured a high proportion of available information about their targeted variable (i.e., thick colored curves approached this gray curve), at least for periods of the trial. However, we sought to quantify the extent to which the sRAs did *not* capture the available information and to compare this extent to that expected for an arbitrary trio of static dimensions. As such we computed the difference *d*_*k,i*_(*t*) = ‖dRA_*k*_(*t*)‖ – *z*_*k,i*_(*t*), or the population encoding of variable *k* undetected by sRA_*i*._ We developed a statistical null model for *d*_*k,i*_(*t*) by generating *S* random sets of three orthogonal vectors (i.e., null sRAs) biased to the dimensionality of the data using the same biased sampling method as for single vectors (Methods, Random dimensions in neural space). We then projected dRA_*k*_(*t*) onto each null sRA from a given random set, computed the null magnitude 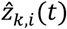 and corresponding null difference 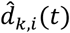, and compiled across random sets so as to ultimately generate a null distribution 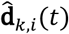 of undetected information. Because the null sRAs were generated without regards to the variables of interest, there was no *a priori* pairing between dRA_*k*_(*t*) and a given null sRA. Therefore, we designated the null sRA most aligned to the dimension of greatest variance in the data as the “on-target” null sRA and used this null sRA to compute 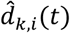 for *i* = *k*. (This designation was the most conservative approach, i.e., produced the smallest null differences 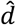 and thus was most likely to identify a given observed difference *d* as statistically large). The undetected information about off-target variables (i.e., 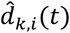 for *i* ≠ *k*) was of less interest, and so the remaining two null sRAs were assigned arbitrarily to the remaining two “off-target” variables. (However, below we summed 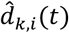 across all null sRAs *i* for computing 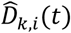.) **(b)** The difference *d*_*k,i*=*k*_(*t*) is shown for the on-target variables in thick curves following the same color and column conventions as in (a). The 50^th^ and 99^th^ percentiles of the corresponding null distribution 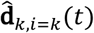 are provided for reference (dashed and dotted curves, respectively). Finally, we computed the probability p_*k,i*_(*t*) of obtaining the observed *d*_*k,i*_(*t*) or greater by chance as the upper-tail of 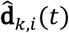 from *d*_*k,i*_(*t*) to ∞. (Note that we integrated 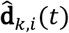 numerically, and thus p was lower-bounded at 1/*S*.) When p_*k,i*_(*t*) was sufficiently large, we concluded that the amount of encoded information undetected by sRA_*i*_ was no more than expected by chance. **(c)** We plotted log_10_ p_*k,i=k*_(*t*) for the on-target variable using the same color and column conventions as in (a) and labeled the threshold p = 0.01 (horizontal dashed line) for reference. We observed that CHOICE and EXPECTED REWARD generally detected a consistent proportion of available population encoding (i.e., consistent spacing between thick colored and gray curves, (a)), suggesting that the observed dynamics of RSV(*t*) or V(*t*) (Figure 4c,d or Supplementary Figure 14a, respectively) were due to changes in the magnitude of the dynamic representation (i.e., ‖dRA_*k*_(*t*)‖), not due to changes in the direction of representation. Conversely, for BENEFIT, we observed both a decrease in the dynamic representation magnitude with time (i.e., decrease in gray curve, (a)) *and* a decrease in the proportion of detected population representation (i.e., increase in difference *d*_*k,i*_(*t*), (b)), suggesting that the representation of benefit was dynamic both in magnitude and direction (i.e., contributions of individual units). For all sRAs at all times, the magnitude of *un*detected encoding of the on-target variable (i.e., *d*_*k,i*=*k*_(*t*)) was less than expected by 99% of random sRAs (c). In other words, the high sensitivity of a given static population representation for a specific variable at key times (e.g., early for BENEFIT, late for CHOICE) did not compromise the ability of the representation to detect the encoding of that variable at other times in the trial (compared to any arbitrary trio of static dimensions). **(d)** We were also interested in how well the entire static three-dimensional (3-D) space spanned by the sRAs captured the dynamic representations. As such, we measured the magnitude *Z*_*k*_(*t*) of the projection of dRA_*k*_(*t*) into the 3-D space, taken as the vector norm 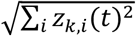 across sRAs *i* (recall that the sRAs were orthogonal). We plotted *Z*_*k*_(*t*) (black curve) as a function of time and included the magnitude of dRA_*k*_(*t*) in the high-dimensional space (gray curve) as an upper-bound for *Z*_*k*_(*t*). Like for the individual sRAs, we measured the extent of information *un*captured by the 3-D space as the difference, *D*_*k*_(*t*) = ‖dRA_*k*_(*t*)‖ − *Z*_*k*_(*t*), between the magnitudes of the dRA in the high-D and 3-D spaces. Likewise, to generate a null model, we computed the difference, 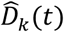, between ‖dRA_*k*_(*t*)‖ and the magnitude of dRA_*k*_(*t*) projected into a given set of three orthogonal random vectors, 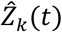. Finally, we compiled 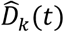 across random sets to produce the null distribution 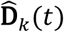. **(e)** The difference *D*_*k*_(*t*) (thick solid black curves) and 50^th^ and 99^th^ percentiles of 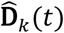 (dashed and dotted gray curves, respectively) are plotted. As above, we computed the probability P_*k*_(*t*) of obtaining *D*_*k*_(*t*) or greater by chance as the upper-tail of 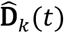 from *D*_*k*_(*t*) to ∞. **(f)** We plotted log_10_ P_*k*_(*t*) (black curve) and labeled the threshold P = 0.01 (horizontal dashed line). The static, 3-D space captured a high proportion of the population encoding of the task-relevant variables (black curves are close to gray curves, (d)). Exceptions tended to occur during periods of lower absolute representation (i.e., smaller ‖dRA_*k*_(*t*)‖), such as the representation of benefit or choice late or early in the trial, respectively. However, we noted that the proportion of available expected reward representation captured by the low-dimensional space was considerably lower than for the other task-relevant variables, including during periods of relatively large dynamic representation. Statistically, for all variables at all times, the magnitude of *un*detected encoding (i.e., *D*_*k*_(*t*)) was less than expected by 99% of random sets of sRAs (f). In other words, the static low-dimensional space, in addition to representing the variables of interest at key times, was no worse than chance at capturing the available population encoding of the variables across all times in the trial. Though not its intention, the present analysis afforded an alternative measure of specificity to complement Supplementary Figure 15. The extent of information about a given variable captured by the low-dimensional space was generally captured by the sRA specific to that variable (i.e., thick curves greater than thin curves, (a)). An exception was the dynamic representation of benefit (a, left panels), which was captured by BENEFIT early in the trial but by CHOICE and EXPECTED REWARD in the mid-to-late trial. This cross-talk reflected the co-alignment between benefit and choice or expected reward representations mid-trial that may have related to the computation of expected reward from benefit (see Supplementary Figure 26 for alignment between dRAs).

### Principal component analysis

**Supplementary Figure 17.**
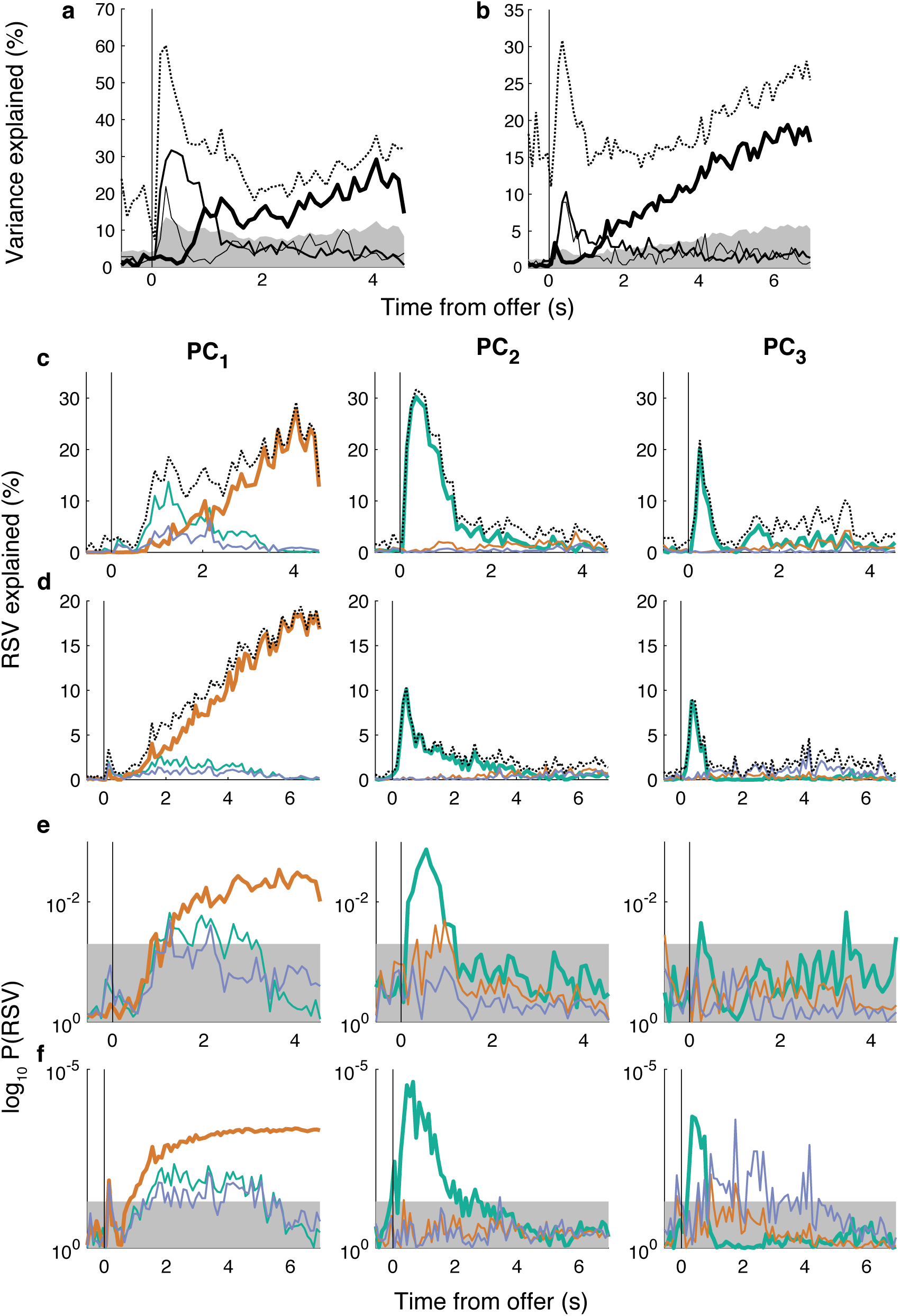
Variance explained by principal components. Principal components analysis (PCA) provided an alternative, traditional approach to dimensionality reduction. We compared the low-dimensional space spanned by the sRAs to that spanned by the top three principal components (PCs), as discovered by PCA on variance across time and conditions in the high-dimensional space. As with oTDR, the analysis was performed on z-transformed, common condition-subtracted responses, 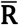, which were reshaped to matrix form with rows (dimensions) corresponding to units and columns (observations) corresponding to concatenation of time bins and conditions. **(a,b)** Variance explained by PC_1_ (thick solid curve), PC_2_ (medium solid curve), and PC_3_ (thin solid curve) is shown as a function of time from the onset of the offer period (black vertical line) for monkeys N (a) and K (b). The area of gray shading represents the variance explained by 95% of random vectors reflecting the dimensionality of the data, and the black dotted curve represents the variance explained by the top PC from a separate analysis in which we performed PCA independently at each time bin to estimate the upper-bound for variance explained by any single, time-varying dimension. Compared to the sRAs (Supplementary Figure 14), the PCs explained less peak variance early in the trial and moderately greater variance toward the end of the trial. However, the variance explained by the top PCs was not necessarily related to the variables of interest. To address the specificity of the PCs, we computed the relevant signal variance (RSV) explained by each PC with respect to the variables of interest. Because the PCs were not targeted to specific variables *a priori*, we designated as “on-target” the variable for which the cumulative RSV (summed across time bins) was greatest: choice for PC_1_ and benefit for PC_2_ and PC_3_ **(c,d)** The on- and off-target RSV (thick and thin solid curves, respectively) explained by PC_1_ (left panel), PC_2_ (middle panel), and PC_3_ (right panel) is plotted as a function of time from the onset of the offer period (vertical black line) for monkeys N (c) and K (d). Colors refer to the task-relevant variables with respect to which RSV was computed: benefit (green), choice (orange), and expected reward (blue). The overall variance (V) explained by each PC (thin black dotted curve) recapitulates the solid curves in (a,b) and served as an upper-bound for RSV. Any lack of specificity of a given PC for a given variable arose within the difference between V and on-target RSV. We quantified the portion of variance related to the “off-target” variables (i.e., the two variables not designated as on-target) as the off-target RSV, the computation of which controlled for the correlation between on- and off-target variables (see Methods and Supplementary Figure 15). **(e,f)** The log_10_ probability of observing on- and off-target RSV (thick and thin curves, respectively) by chance is plotted for monkeys N (c) and K (d), where panels and colors are as in (c,d). Probabilities were derived from empirical null distributions of on- and off-target RSV (see Methods). The area of gray shading represents p > 0.05. Compared to the sRAs (Supplementary Figure 15), the top PCs were less specific to any given variable of interest (i.e., greater difference between V and on-target RSV, (c,d)). Moreover, the variance explained by the PCs that was *unrelated* to the on-target variable was not merely irrelevant to the task, but rather was significantly related to the off-target variables (e,f), more so than for the sRAs, such as RSV explained by PC_1_ with respect to benefit and expected reward (left panels, thin green and blue curves), RSV explained by PC_2_ with respect to choice for monkey N (middle panel, thin orange curves), and RSV explained by PC_3_ with respect to expected reward for monkey K (right panel, thin blue curves). Finally, the sensitivity for a given variable of interest was spread between PCs. In particular, benefit was represented significantly and most greatly by PC_2_ and PC_3_, rather than consolidated onto a single dimension, as by the BENEFIT sRA. In summary, compared to the sRAs, any given PC was less sensitive and specific to a particular variable of interest, and thus the PCs less well separated the task-relevant signals into low-dimensional representations specific to each variable.

### Common condition response

**Supplementary Figure 18.**
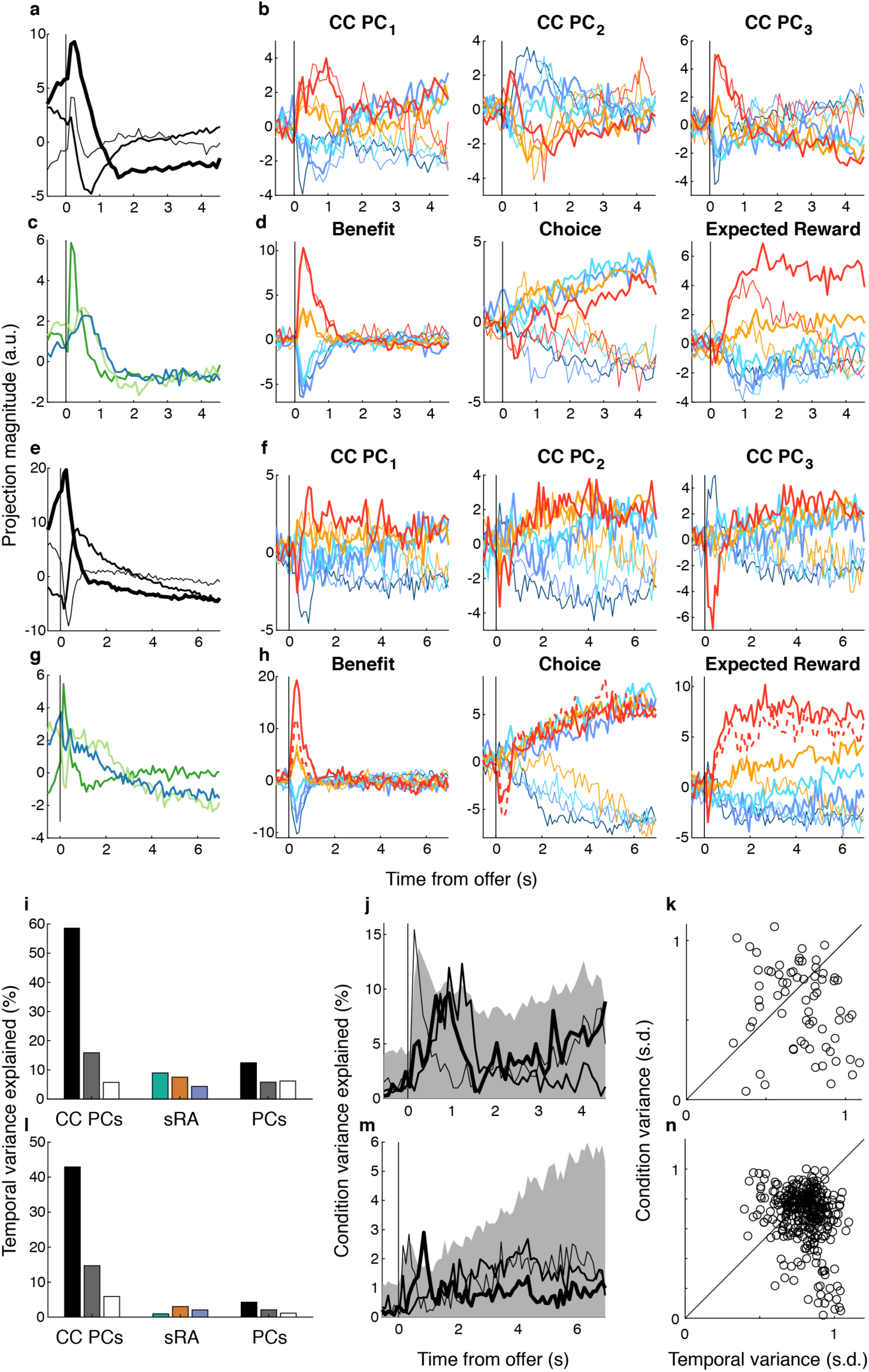
Common condition response. For monkeys N (a-d, i-k) and K (e-h, l-n), we examined the common-condition response (CC), i.e., the mean response across conditions taken at each time bin, and compared the CC to the condition-specific response 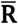 from which the CC was subtracted. In particular, we were interested in comparing the temporal variance of the CC (i.e., variance across time bins) to the cross-condition variance (i.e., variance across conditions at each time bin) in terms of both magnitude and overlap of the low-dimensional subspaces that captured the temporal or cross-condition variance. We performed principal component analysis (PCA) on the CC (units x time bins), thus discovering the top three principal components (CC PCs) that captured the greatest temporal variance. To measure the “activity” of a given representation (as in Figure 4a,b), we projected the CC onto **(a,e)** CC PC_1_ (thick curve), CC PC_2_ (medium curve) and CC PC_3_ (thin curve), and onto **(c,g)** the sRAs of BENEFIT (green curve), CHOICE (orange curve), and EXPECTED REWARD (blue curve) and plotted the resulting magnitude as a function of time from the onset of the offer period (vertical line). Likewise, we projected 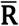 (i.e., condition-specific response) onto **(b,f)** CC PC 1, 2 or 3 and **(d,h)** sRAs of BENEFIT, CHOICE, or EXPECTED REWARD (left, middle, or right panels, respectively), where curve color and thickness represent offer size and choice, respectively, as in Figure 4. The magnitudes of the projections are in arbitrary units related to the z-transformed firing rates but are comparable between dimensions (i.e., CC PCs vs. sRAs) and animals. The projections of 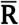 onto the sRAs (d,h) are identical to those in Figure 4a,b, and are recapitulated here (with explicit magnitudes added) to facilitate comparison to projections onto the CC PCs (b,f). We next examined the extent to which the various low-dimensional subspaces captured temporal and cross-condition variance. The magnitude of projecting the CC onto the CC PCs was greater than for onto the sRAs (a vs. c and e vs. g; note different y-axes), suggesting the CC PCs captured greater temporal variance than the sRAs. **(i,l)** To test this observation, we measured the percent of temporal variance (bar height) explained by the CC PCs, sRAs, and, for completeness, the condition-sensitive principal components (PCs; discovered in matrix 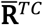 and described in Supplementary Figure 17), where black, gray, and white bars refer to the first, second and third PC or CC PC, and green, orange and blue bars refer to the sRAs of BENEFIT, CHOICE, and EXPECTED REWARD, respectively. Indeed, the CC PCs explained markedly greater temporal variance than the PCs or sRAs. In contrast, when projecting the condition-specific neural response (i.e., 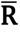) onto the various dimensions, the range of magnitudes for the CC PCs (b,f) was less than for the sRAs (d,h), suggesting the CC PCs captured less cross-condition variance than the sRAs. In addition, the extent to which the CC PCs discriminated the variables of interest (i.e., orderly separation of projections by offer size and/or choice) was qualitatively less than for the sRAs. **(j,m)** Indeed, the cross-condition variance explained by CC PC_1_ (thick curve), CC PC_2_ (medium curve) and CC PC_3_ (thin curve) was less than expected by 95% of random vectors reflecting the dimensionality of the data (gray shading) and less than explained by the sRAs (compare to Supplementary Figure 14a,b). (Whereas the present figure compares variance explained for individual dimensions across the three static subspaces, see Supplementary Figure 19 for comparison of the subspaces as a whole.) Finally, we were interested in directly comparing the magnitudes of temporal and cross-condition variance in OFC. That is, were the condition-specific responses dwarfed by the time-varying, condition-independent response, or were these two sources of variance comparable? Qualitatively, we observed that the range of magnitudes of the projections of the CC onto the CC PCs (a,e) was comparable to that of 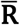 onto the sRAs (d,h), suggesting that condition-independent and condition-specific responses were of similar magnitude in OFC. To test this hypothesis more rigorously, for each unit *n*, we computed the temporal variance for each condition *c* from the z-transformed, trial-average responses 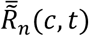, which contained the CC, and took the median variance across conditions. Separately, for each unit, we computed the cross-condition variance of 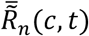 at each time bin and computed the median variance across time bins. **(k,n)** We plotted the median temporal (abscissa) and cross-condition (ordinate) variance for each unit (circles) in terms of standard deviations from the unit’s mean response across all times and conditions. Points below the unity line (black diagonal) indicated greater temporal than cross-condition variance, and vice versa. We computed the pairwise difference of temporal minus cross-condition variance, which was significantly greater than zero for the population (median = 0.10 or 0.07 and p = 0.0075 or 2.5 * 10^−15^ via Wilcoxon signed-rank test, monkey N or K, respectively), indicating greater temporal than condition variance. However, the magnitude of this difference was small: less than 0.1 s.d. on average and always less than 1 s.d. Thus, we concluded that OFC responses exhibited marginally but significantly greater temporal than condition variance.

**Supplementary Figure 19.**
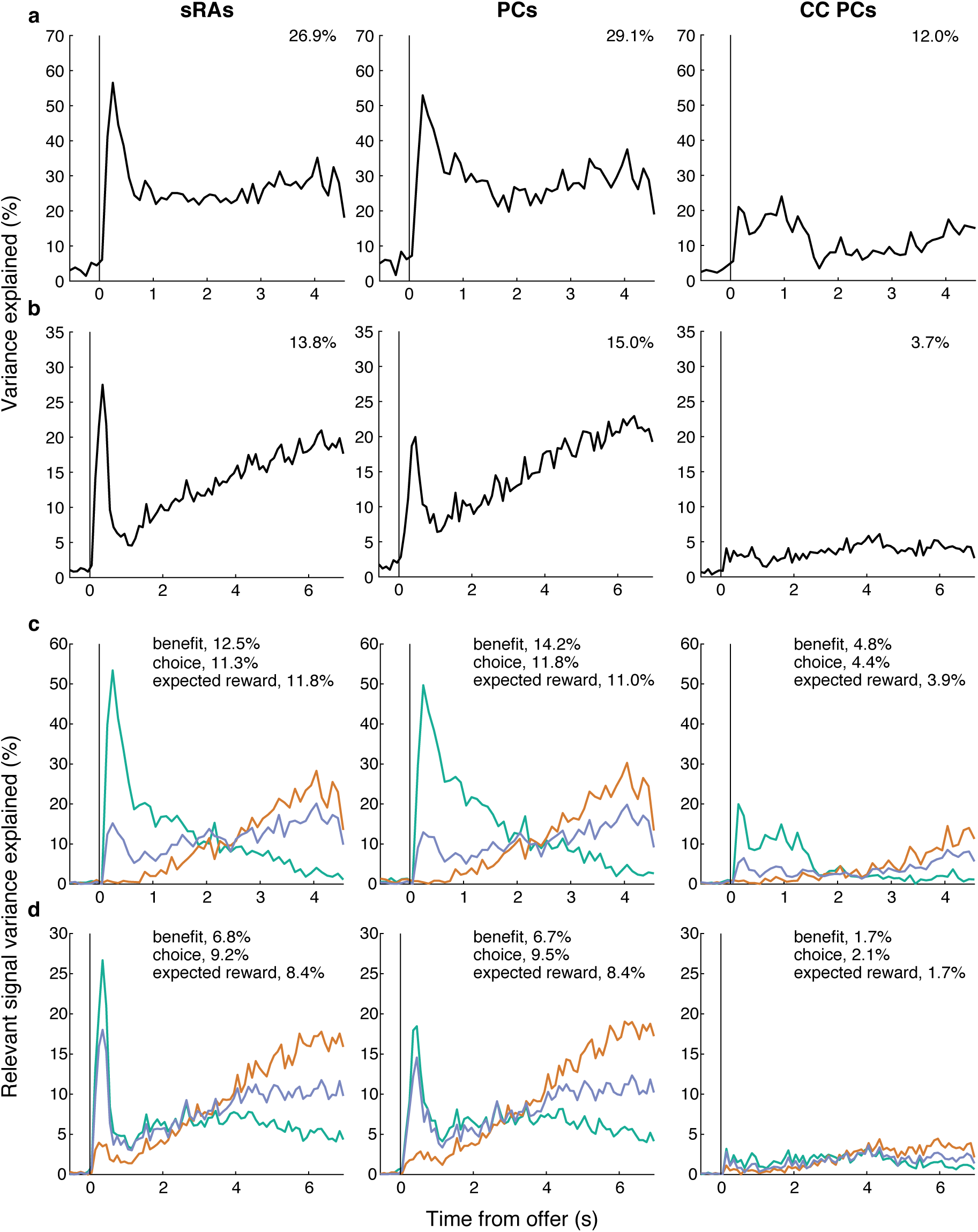
Variance explained by 3-D spaces. We were interested in comparing the explanatory power of the above 3-dimensional subspaces—static regression axes (sRAs; left panels; see Figure 4 and Supplementary Figure 14), principal components (PCs; middle panels; see Supplementary Figure 17), and common-condition principal components (CC PCs; right panels; see Supplementary Figure 18)—as a whole, instead of just their component dimensions examined separately. **(a,b)** Variance or **(c,d)** relevant signal variance (RSV) explained by a given 3-D subspace (as summed over the three contributing orthogonal dimensions) is plotted as a function of time from the onset of the offer period (vertical black line) for monkeys N (a,c) and K (b,d). The mean variance or RSV across time bins is indicated in the top right of each panel. For RSV, line color refers to the task-relevant variable with respect to which RSV was computed: benefit (green), choice (orange), and expected reward (blue). In terms of variance (a,b), the PCs explained the greatest mean variance, as expected by the design of principal components analysis. However, the variance explained by the sRAs was nearly as great on average and greater at peak time bins, consistent with a high proportion of variance in the neural population as a whole being related to the task-relevant variables. In contrast, the space capturing temporal variance, as defined by the CC PCs, explained much less of the cross-condition variance, consistent with the temporal and cross-condition variances occupying largely non-overlapping dimensions (see Supplementary Figure 18). In terms of RSV (c,d), the comparison between the sRAs and PCs was similar as for variance explained. However, the mean RSV explained by the sRAs was even closer to and at times exceeded the mean RSV explained by the PCs, consistent with the greater sensitivity of the sRAs to the task-relevant variables, despite the marginally greater overall variance explained by the PCs. Note that we summed RSV across sRAs without controlling for correlations between the variables, in contrast to our prior analyses that used partial correlation to compute RSV for off-target variables (Supplementary Figure 15). As a result, the total RSV across variables explained by the 3-D subspace (i.e., if one were to sum the three colored lines in (c,d)) could take values greater than variance explained by the same subspace (a,b). We emphasize that, for a given dimension, RSV was always bounded by variance explained, and likewise the sum of RSV for a given variable across dimensions was bounded by variance explained summed across those same dimensions. In addition to comparing the variance explained by the various subspaces, we compared their overlap directly as measured by the alignment index (Methods), which ranged from 0 to 1. The overlap between the sRA and PC subspaces was high (alignment index = 0.84 or 0.65, monkey N or K, respectively), consistent with the OFC population being primarily modulated by the task-relevant variables. In contrast, the overlap between the sRA and CC PC subspaces was small (alignment index = or 0.11, monkey N or K, respectively), consistent with the relative specificity of the CC PCs or sRAs for temporal or cross-condition variance, respectively.

### Stability of dynamic representations

**Supplementary Figure 20.**
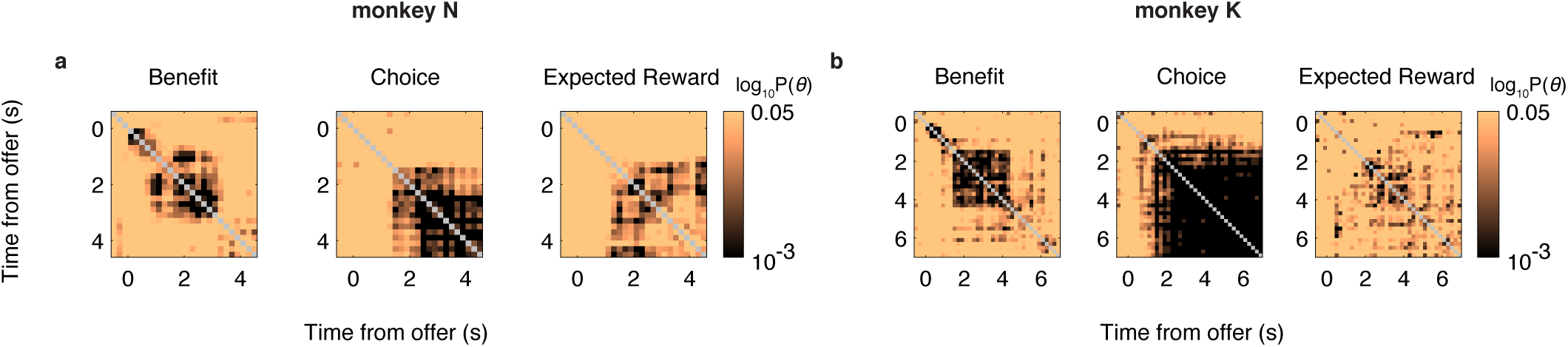
Probability of pairwise similarity between dynamic representations. **(a,b)** The log_10_ probability of observing by chance the similarity *θ* between each pair of dynamic low-dimensional representations (dRAs; see Figure 5) is given by the pixel color (referencing right-sided color scale), and the times of the dRA pair (relative to the onset of the offer period) are given by the pixel’s row and column positions for task-relevant variables of benefit (left column), choice (middle column), and expected reward (right column) for monkeys N (a) and K (b). The diagonal compares identical dRAs (*θ* = 0, p(*θ*) = 1) and is colored gray.

**Supplementary Figure 21.**
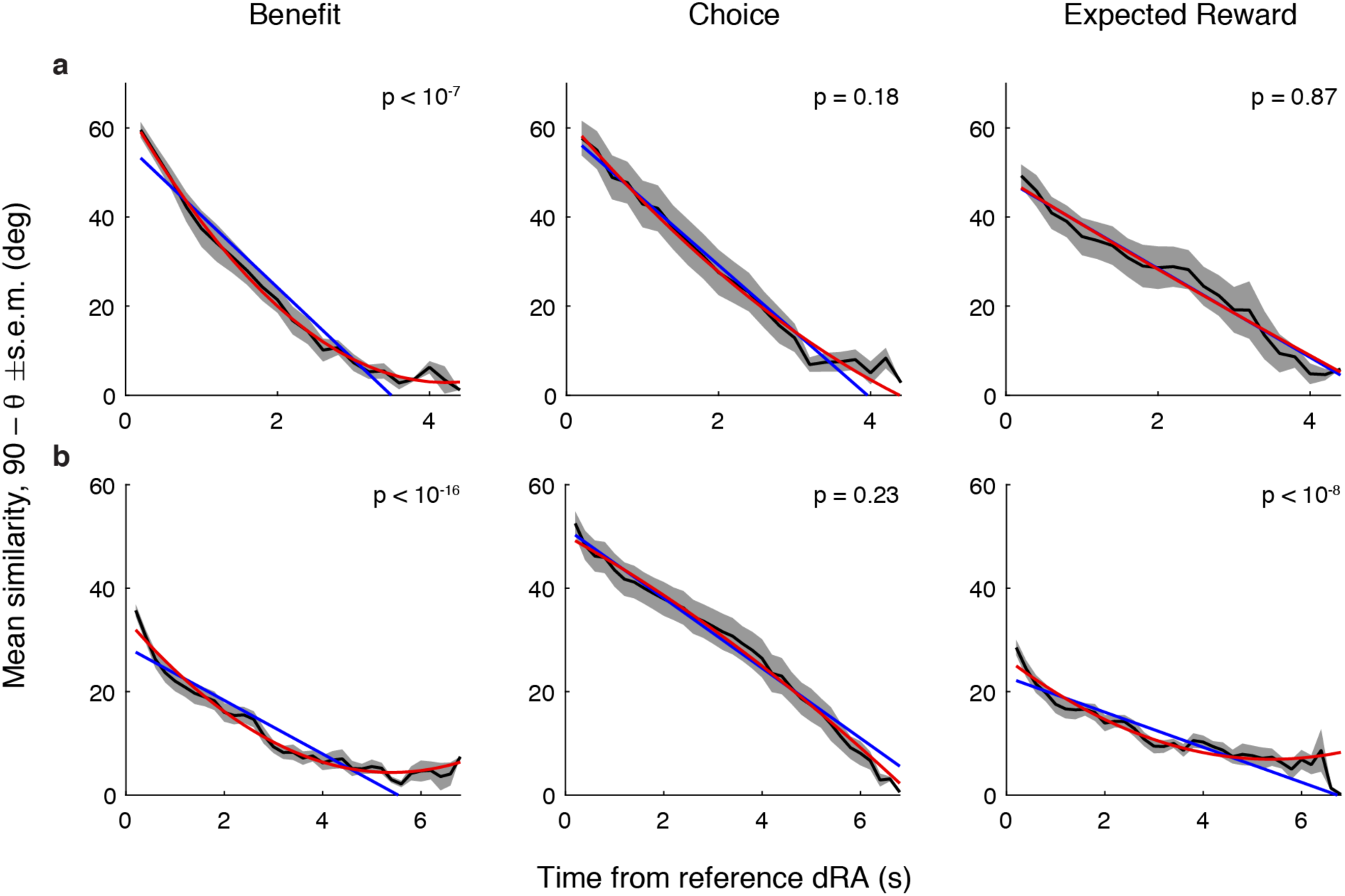
Average similarity between dynamic representations. We computed the similarity between pairs of dynamic representations (dRAs) as the time-varying angle *θ*(*t, dt*) between reference dRA(*t*) and dRA(*t*+*dt*) for time *t* > 0 relative to the onset of the offer period and temporal offset *dt* relative to *t*. (*θ*(*t, dt*) was equivalent to black traces in Supplementary Figure 22.) **(a,b)** We plotted the mean angle 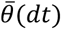 (black curve) and associated s.e.m. (gray shading), averaged across times *t*, as a function of offset *dt* for task-relevant variables benefit (left column), choice (middle column), and expected reward (right column) for monkeys N (a) and K (b). As 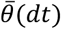 was symmetric about *dt* = 0, mean angles are shown only for *dt* ≥ 0. The probability p of the null hypothesis that the function *θ*(*t, dt*) (estimated across observations *t*) was explained as well by a first-order (blue line) as a second-order (red curve) polynomial, controlling for degrees of freedom, is shown in the top-right corner of each panel (*F*-test of likelihood ratios). Sufficiently small values of p suggested that the relationship was non-linear. A recent report argued that representational similarity in prefrontal cortex decayed smoothly with *dt* and reflected the timescale of dynamic encoding (see Figure 3c,d in Ref^45^). This “common dynamics” hypothesis implied that *θ*(*t, dt*) decayed equivalently for all times *t* and for representations of all variables (i.e., benefit, choice, and expected reward). As such, we would expect the similarity *θ*(*t, dt*) to decay smoothly with *dt* for all times *t* and to share similar shape and magnitude across variables. On the other hand, we considered that similarity was a feature of time- and/or variable-dependent behavioral functions subserved by the representations. This “functional dynamics” hypothesis implied time- and variable-dependent predictions. If the behavioral function varied with time, then *θ*(*t, dt*) would depend on *t* according to the time course of the associated function. As such, we would expect 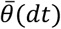 to be a non-linear function of *dt* for one or more reasons. First, *θ*(*t, dt*) may be non-linear without dependence on *t* (this result would be consistent with both the common and functional dynamics hypotheses). Second, *θ*(*t, dt*) may be linear, but the slope and/or offset may depend on *t*, and thus 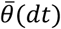 would take an exponential form. Third, *θ*(*t, dt*) may be non-linear *and* depend on *t*. In particular, we considered when *θ*(*t, dt*) took a sigmoidal form or threshold non-linearity, as apparent in some traces in Supplementary Figure 22, resulting in a sigmoidal form for 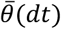. In addition, the functional hypothesis predicted that 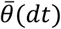 would depend on the specific variable being represented, as different variables may have fundamentally distinct representations and/or the time course of functional relevance may differ between variables (e.g., benefit being relevant briefly, while choice being relevant over a longer time period). In contrast, the common dynamics hypothesis predicted that 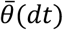—regardless of its particular form—would not depend on the specific variable. In the present figure, we observed that 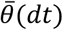 appeared to take several non-linear forms, including exponential, sigmoidal, and threshold-linear, across the task variables and animals. Statistically, we found that 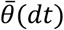 was significantly (p < 0.05) better explained by a non-linear than linear function in half the cases. Finally, we observed that the magnitude and particular form of 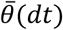 appeared to depend on the specific variable being represented. While the above analysis did not distinguish between the many possible causes of time- and/or variable-dependent forms of 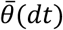 outlined above, it argued strongly against the common dynamics hypothesis and motivated a finer examination of how similarity between representations depended on the specific time and variable being represented.

**Supplementary Figure 22.**
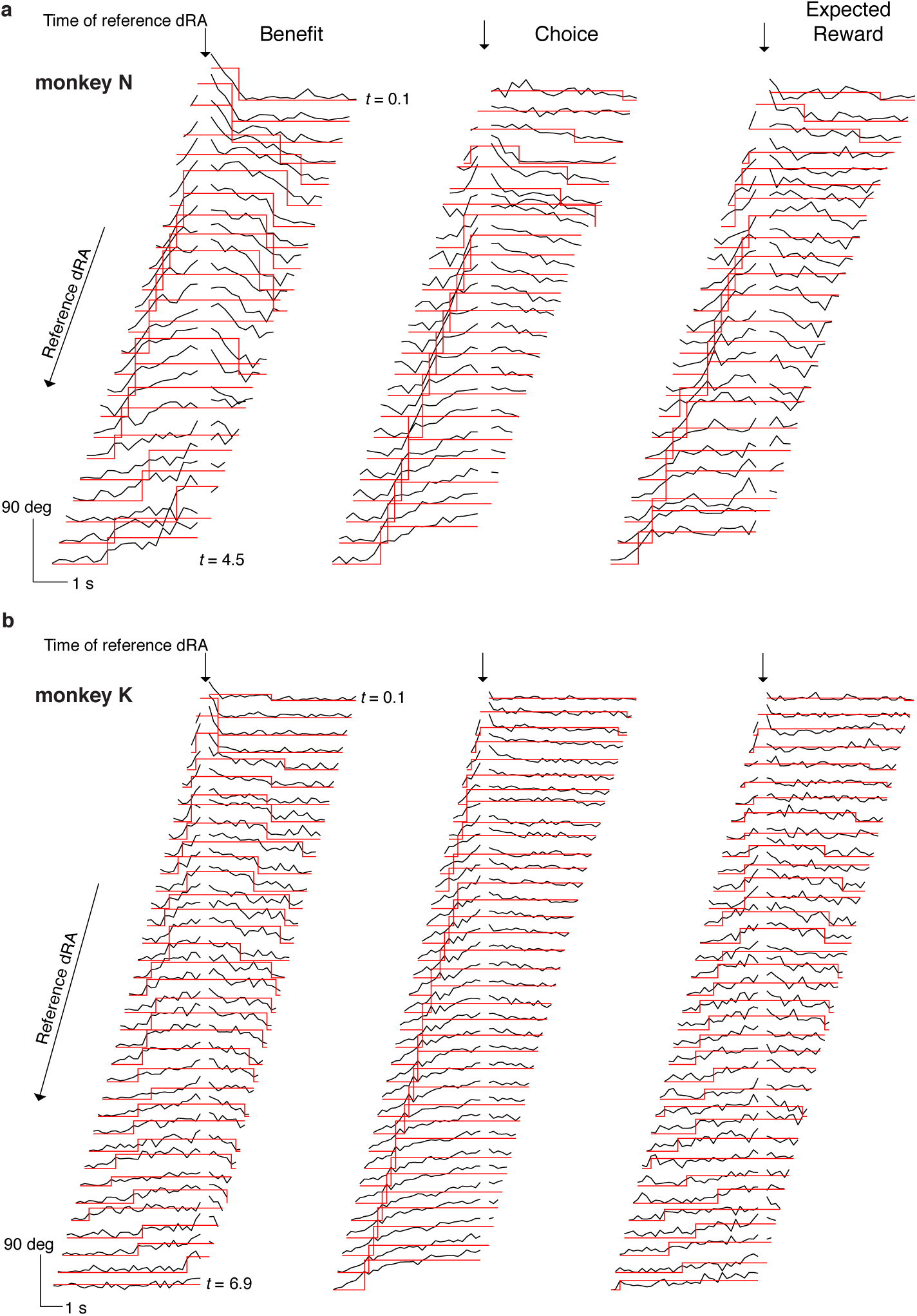
Boxcar functions fit to similarity between dynamic representations. **(a,b)** The time-varying angle *θ* between pairs of dynamic low-dimensional representations (dRAs) for the task-relevant variables of benefit (left column), choice (middle column), and expected reward (right column) are shown for monkeys N (a) and K (b). Each black trace indicates 90° – *θ* (such that higher values correspond to greater similarity) between reference dRA(*t*), or dRA*, taken at time *t* and all other dRA(*t*+*dt*) taken at temporal offset *dt* relative to dRA*. The absolute trial time of dRA* is given by the row position of the black trace from early (top) to late (bottom) in the trial (center of time bin for first and last dRA* is labeled) and corresponds to the rows in Figure 5c,d. The spacing between black traces is arbitrary and for display purposes only. Black traces are aligned horizontally to the time of dRA* (downward arrow) and are redacted at this time (white space) since 90° – *θ* = 90° when dRA(*t*+*dt*) = dRA*. Boxcar functions (red traces) were least-squares fit to the black traces, excluding the time of dRA* and times before the offer (i.e., excluding t < 0). The width of the boxcar function indicated the putative period of stability and the boxcar height indicated the magnitude of similarity, as depicted in Figure 5e,f. Horizontal and vertical scale bars indicate trial time and dRA similarity (i.e., 90° – *θ*), respectively.

**Supplementary Figure 23.**
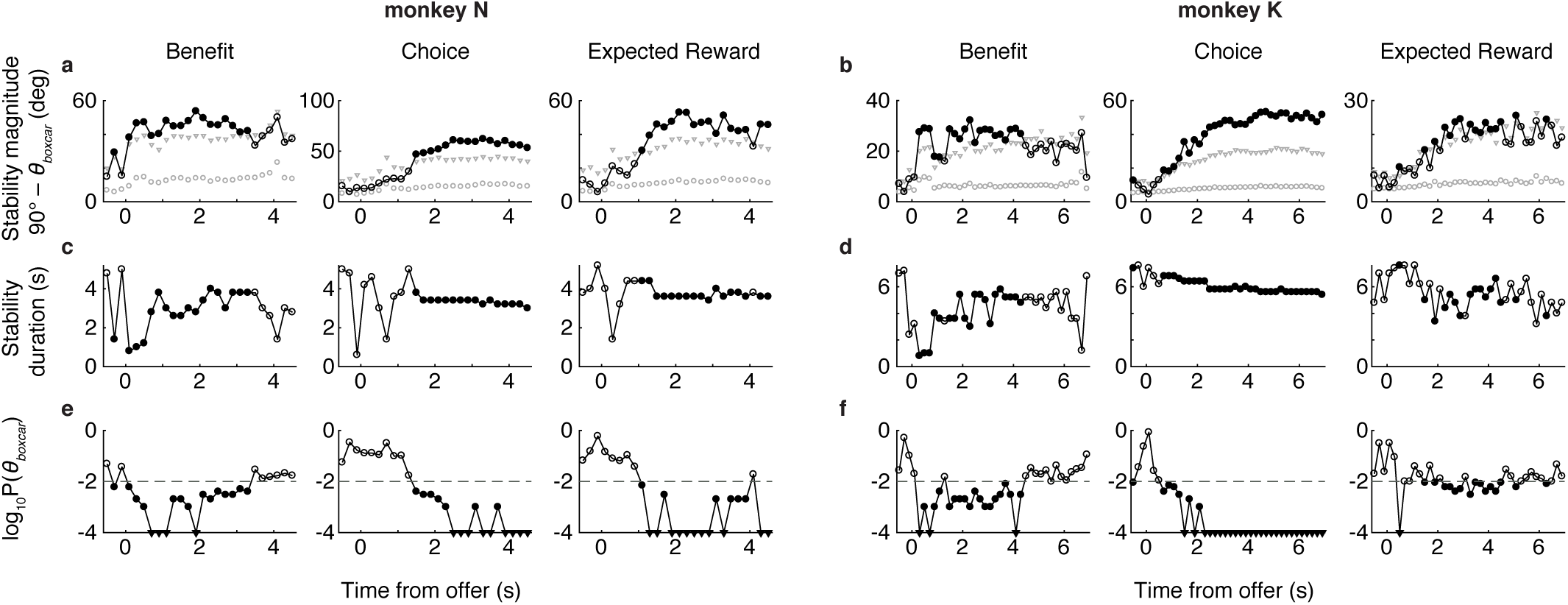
Boxcar function parameters and statistics. The parameters and statistics of the boxcar functions fit to the angle between dynamic low-dimensional representations (dRAs) for task-relevant variables of benefit (left column), choice (middle column), and expected reward (right column) are shown for monkeys N (a,c,e) and K (b,d,f). Boxcar fits were used to discover putative periods of stability with respect to a reference dRA* and adjacent dRAs (Supplementary Figure 22) and the associated parameters are shown as a function of the time of the reference dRA* relative to the onset of the offer period. **(a,b)** The boxcar height *θ*_boxcar_, i.e., magnitude of similarity, is plotted (open or filled circles) as 90° – *θ*_boxcar_ (i.e., larger values indicate greater similarity). In all panels (a-f), filled circles indicate that the probability p of observing *θ*_boxcar_ by chance was p(*θ*_boxcar_) < 0.01, as computed from the distribution of chance angles 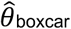 between surrogate dRAs (Methods). The median (gray circles) and 99^th^-percentile (gray triangles) of the null distribution 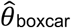 is shown for each time bin. **(c,d)** The boxcar width, i.e., duration of similarity, is plotted. **(e,f)** The log_10_ p(*θ*_boxcar_) is plotted (also shown by color of vertical bands to left of the heat maps in Figure 5e,f). Filled triangles indicate p < 10^−4^, which occurred when *θ*_boxcar_ was less than all 10,000 values of 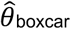. The arbitrary threshold of p = 0.01 (dashed horizontal line) appeared to discriminate categorically distinct ranges of p(*θ*_boxcar_), justifying use of this threshold for display purposes (e.g., Figure 5e,f). However, the periods of putative stability were not consistently below this threshold. In particular, p(*θ*_boxcar_) for monkey K expected reward ((f), right panel) hovered around the significance threshold. In addition, the stability magnitude during these periods ((b), right panel, note smaller y-axis scale) was likewise smaller than for other variables. In general, we considered both similarity magnitude and statistical significance when interpreting periods of putative stability.

**Supplementary Figure 24.**
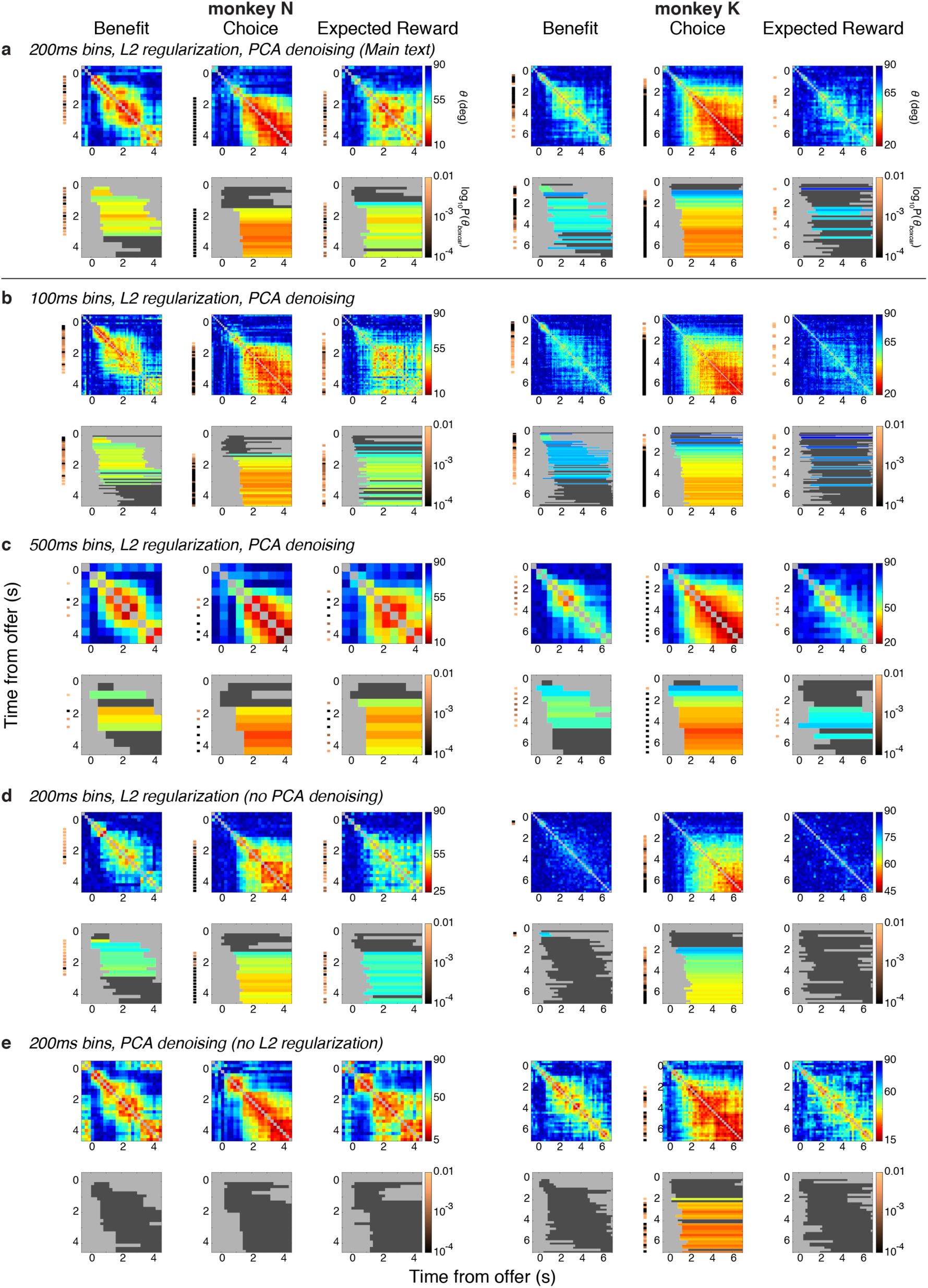
Effect of varying model parameters on stability analysis. **(a-e)** For monkeys N (left 3 columns) and K (right 3 columns), the stability of the dynamic low-dimensional representations (dRAs) of benefit, choice, and expected reward is shown (left, middle, and right columns, respectively, for a given animal) for separate implementations of the dRA analysis. Each pair of rows corresponds to a separate analysis using distinct parameters (italicized headings). Conventions are as in Figure 5. In brief, for a given pair of rows, the top row (heatmaps) shows the angle *θ* in degrees between pairs of dRAs for the same variable as given by the pixel color (referencing right-hand color scale within row), while the pixel’s grid position gives the times of the dRA pair relative to the onset of the offer. Smaller angles (warmer colors) correspond to greater similarity between representations. The bottom row shows the boxcar fits to the above heatmaps. Periods of significant similarity (i.e., p(*θ*_boxcar_) < 0.01) are colored, with bar color indicating boxcar height (i.e., extent of similarity during period spanned by the boxcar) and referencing right-hand color scale in above row. Non-significant boxcars (p(*θ*_boxcar_) ≥ 0.01) are shaded dark gray. The thin vertical bands to the left of each panel in (c-f) show the log_10_ probability of observing *θ*_boxcar_ by chance for the corresponding time, with colors referencing the right-hand color scale in the lower of each pair of rows (no color is shown when p(*θ*_boxcar_) ≥ 0.01). **(a)** The analysis from the main text is recapitulated in (a). Alternative implementations are discussed below. In general, both the profile of angle similarity (heatmaps) and significance of boxcar fits did not depend qualitatively on the duration of dRA time bins (100, 200, or 500ms) or application of PCA-based denoising or regularization. **(b)** Shortening the time bin from 200ms, as used in main text, to 100ms reduced the precision in estimating the dRA in any given time bin (indeed the dRA magnitudes, not shown, and angles were more variable across time), but did not change the fundamental stability conclusions: benefit was stable early (0 – 0.5 s), then changed to a new representation that was stable during the middle of the trial (0.5 s until about 1 s before the reward); choice was stable from about 1 s onward; and expected reward was stable from about 1 s onward for monkey N, but was equivocal for monkey K. **(c)** Similarly, lengthening the time bin to 500 ms had little effect on the same fundamental conclusions with a single exception: no early period of stability (0 – 0.5 s) was found for benefit. This was easily explained by the fact that this entire period was subtended by a single 500 ms time bin, which by its singular nature could not be “stable” as there was no other time bin (within the period in question) with which to compare it! **(d)** The effect of omitting PCA-based denoising is shown (regularization and 200 ms time bins were maintained). Compared to the main analysis (a), the absolute magnitude of similarity was reduced (note larger minimum angles in red-to-blue color bars), as expected given that the dRAs could now span a higher dimensional space, thus permitting higher extents of dissimilarity. However, this pressure toward dissimilarity was also present for the random surrogate data. Thus the similarity between the veridical dRAs was much greater than between the surrogate dRAs and, consequently, the estimated probability of observing the veridical dRAs by chance remained small (i.e., statistically significant), as observed by qualitatively similar boxcar plots (bottom row in (d)). Monkey K was an exception in that the absolute magnitudes of similarity for benefit (during the middle period) and expected reward were so markedly reduced by the lack of denoising as to be indistinguishable from chance. Monkey K may have been particularly susceptible to the lack of denoising because of the markedly higher number of dimensions in its full dataset (342 units compared to monkey N’s 68), making dissimilarity much more common. In addition, even in the main text analysis (a), the stability for monkey K during these periods was much lower than for monkey N. **(e)** The effect of omitting L2-regularization is shown (PCA-based denoising and 200 ms time bins were maintained). Recall that L2-regularization penalizes extreme values across the set of beta coefficients for the task-relevant variables for a given neuron within a given time bin. That is, the regularization assumes that a given neuron contributes at least partially to the representation of multiple variables (consistent with mixed selectivity). When the representations are robust within the population (i.e., high dRA magnitude), the impact of omitting regularization is minimal. However, when a significant degree of noise is present (i.e., firing rates are poorly explained by the task-relevant variables), unregularized dRAs will reflect these small, random changes in firing rate (i.e., overfitting) and vary accordingly over time. The stability heatmaps (e, top row) are consistent with these predictions: during the trial, when encoding of the task-relevant variables is strong, the similarity of dRAs is comparable to the main text analysis (a), albeit with slightly greater extremes of similarity (note smaller minimum angles in red-to-blue color bars). However, during periods of low or even absent encoding (e.g., prior to the offer), the unregularized dRAs exhibit some similarity. (Note: this similarity is not reflected in the boxcar plots because boxcars fits were limited to times during and after the offer.) The putative similarity during this pre-offer time must be spurious (i.e., resulting from dRAs overfit to random variation) because no signal related to benefit or expected reward could exist prior to the offer. Thus applying regularization to mitigate this overfitting is warranted, as done in the main analysis (a). Finally, the absence of regularization all but eliminated the statistical significance of the boxcar magnitudes (e, lower row). We confirmed this was not due to a reduction in the magnitude of the boxcar fits (not shown), which were nearly identical to those in the main analysis. Rather, the lack of significance was due to a massive increase in the magnitude of boxcars fit to the surrogate data (recall that larger boxcar magnitude corresponds to higher similarity of contributing dRAs). To provide an intuition for this phenomenon, consider finding the dRAs within a single surrogate dataset. Say, by chance, the average firing rate in a given time bin was highly correlated with the offer size, but not so much with the other task-relevant variables. Because this correlation resulted from random variation, it did not generalize well when measured within subsets of trials (i.e., poor cross-validation) and thus would be suppressed by regularization. However, without regularization, the exclusive correlation with benefit and the lack of generalization were tolerated by the analysis, resulting in an unusually large beta coefficient for benefit and negligible coefficients for the other variables. In the adjacent time bins, the fact that temporal smoothness was preserved in the surrogate data caused the same random correlation with benefit to be maintained across several time bins, resulting in spuriously high similarity for that surrogate dataset. Of course, this spurious benefit signal was likely absent in a different surrogate dataset, while a similar pattern of spurious similarity would be observed for a different variable. Across datasets, this process inflated the similarity within the surrogate data and thus increased our estimates of observing a given level of similarity by chance. (The lack of random encoding in some surrogate datasets did not offset the spuriously high encoding/similarity in other datasets because of a floor effect: similarity could not be less than zero.) In contrast, by applying regularization, we mitigated the tendency to overfit the dRAs and reduced the resulting “similarity” of dRAs explaining random variation in firing rate that did not generalize across trials. Ultimately, use of regularization in the main analysis more accurately estimated chance levels of similarity.

**Supplementary Figure 25.**
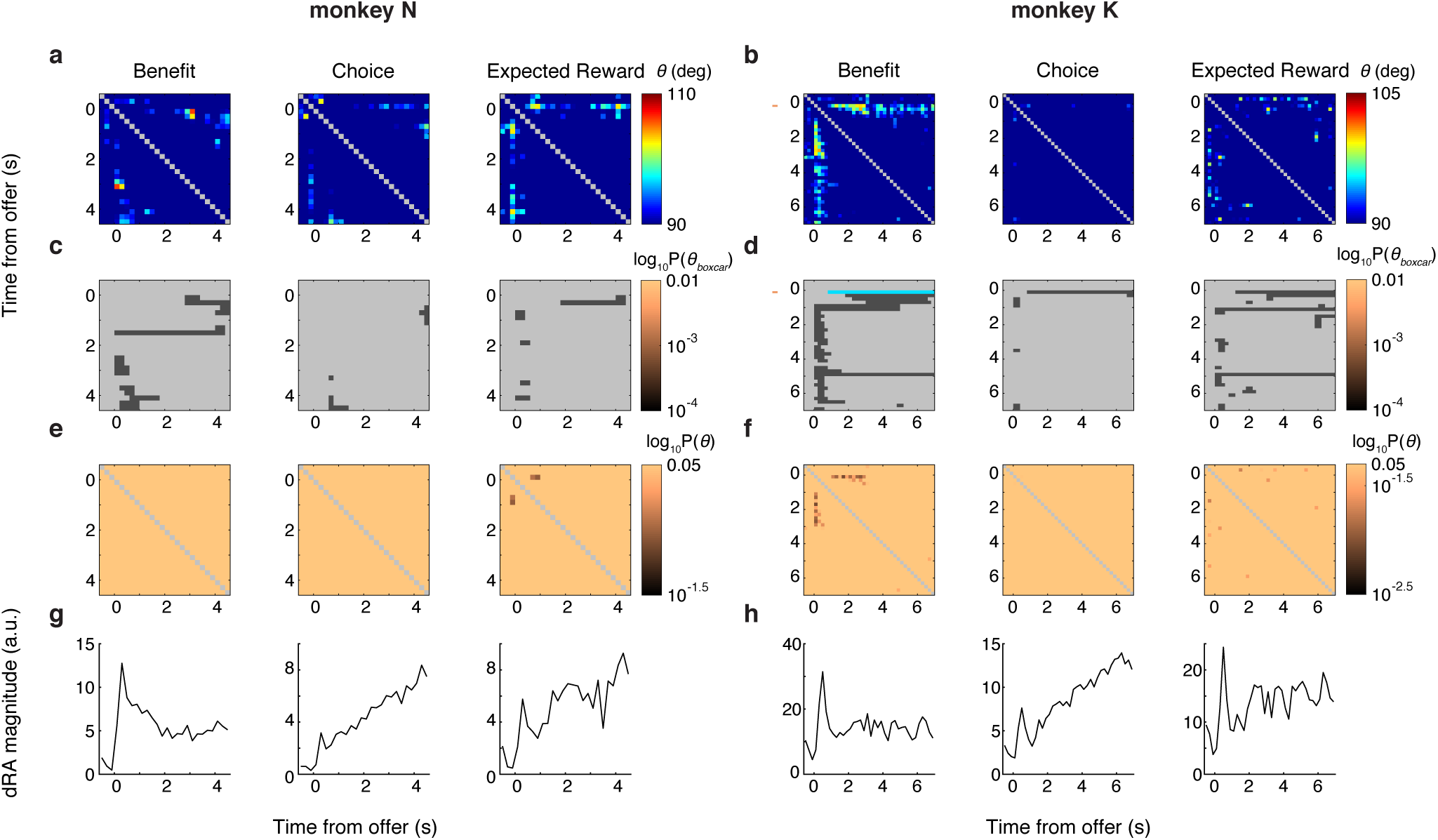
Sign reversals in dynamic low-dimensional representations. The analyses above examined the folded angle *θ* (Eq. 15; Figure 5; Supplementary Figure 20), which measured the absolute similarity between dynamic low-dimensional representations (dRAs) by treating angles equidistant from 90° as equivalent. However, the folded angles did not capture reversals in the sign of representation, i.e., when the absolute magnitude of encoding remained consistent over the course of the trial, but the sign reversed (e.g., Figure 2d). To detect reversals, we computed the unfolded angle *θ*′ (Eq. 16), which was limited to the range [90, 180°]. The present figure examines sign reversals for monkeys N (left 3 columns) and K (right 3 columns) based on *θ*′ between the dynamic representations (dRAs) for benefit, choice, and expected reward (left, middle, and right columns, respectively, for a given animal). **(a,b)** Angle *θ*′ in degrees between a pair of dRAs of the same variable is given by the color of each pixel (referencing right-hand color scale in (a,b)), and the times of the dRA pair (relative to the onset of the offer period) are given by the pixel’s row and column positions. Larger angles (warmer colors) correspond to greater similarity between representations but with opposite sign of representation. Values of *θ*′ < 90° are coded as 90° for display purposes only (these values were excluded from analysis of *θ′*). The diagonal compares identical dRAs and is colored gray and was excluded from analysis. **(c,d)** Boxcar function with baseline of 90° was fit to each row of angles in (a,b), and the period of non-zero boxcar height is indicated by a colored or dark gray horizontal bar in the corresponding row of (c,d). Periods of significant similarity (i.e., 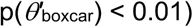 are colored, where bar color represents the boxcar height 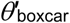, i.e., magnitude of similarity, measured in degrees and referencing the right-hand color scale in (a,b). When the log_10_ probability p of observing 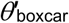 was < 0.01, the value of p is indicated by the color of the vertical band to the left of the corresponding row in (a-d) and references the right-hand color scale in (c,d). (In effect, only a single row for a single variable was significant; (b,d), right panel). Boxcars for which 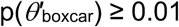 are shaded dark gray in (c,d). Periods where *θ*_boxcar_ ≤ 90° are shown in light gray. **(e,f)** The log_10_ probability of observing by chance the pairwise similarity *θ*′ (a,b)—not the period of contiguous similarity, 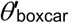 —is given by the color of the corresponding pixel and references the right-hand color scale in (e,f). The diagonal compares identical dRAs and is colored gray. **(g,h)** Recapitulated from Figure 5, the vector magnitude of each dRA is shown as a function of time from the onset of the offer period and provides a reference as to the magnitude of population encoding available for representation at a given time.

**Supplementary Figure 26.**
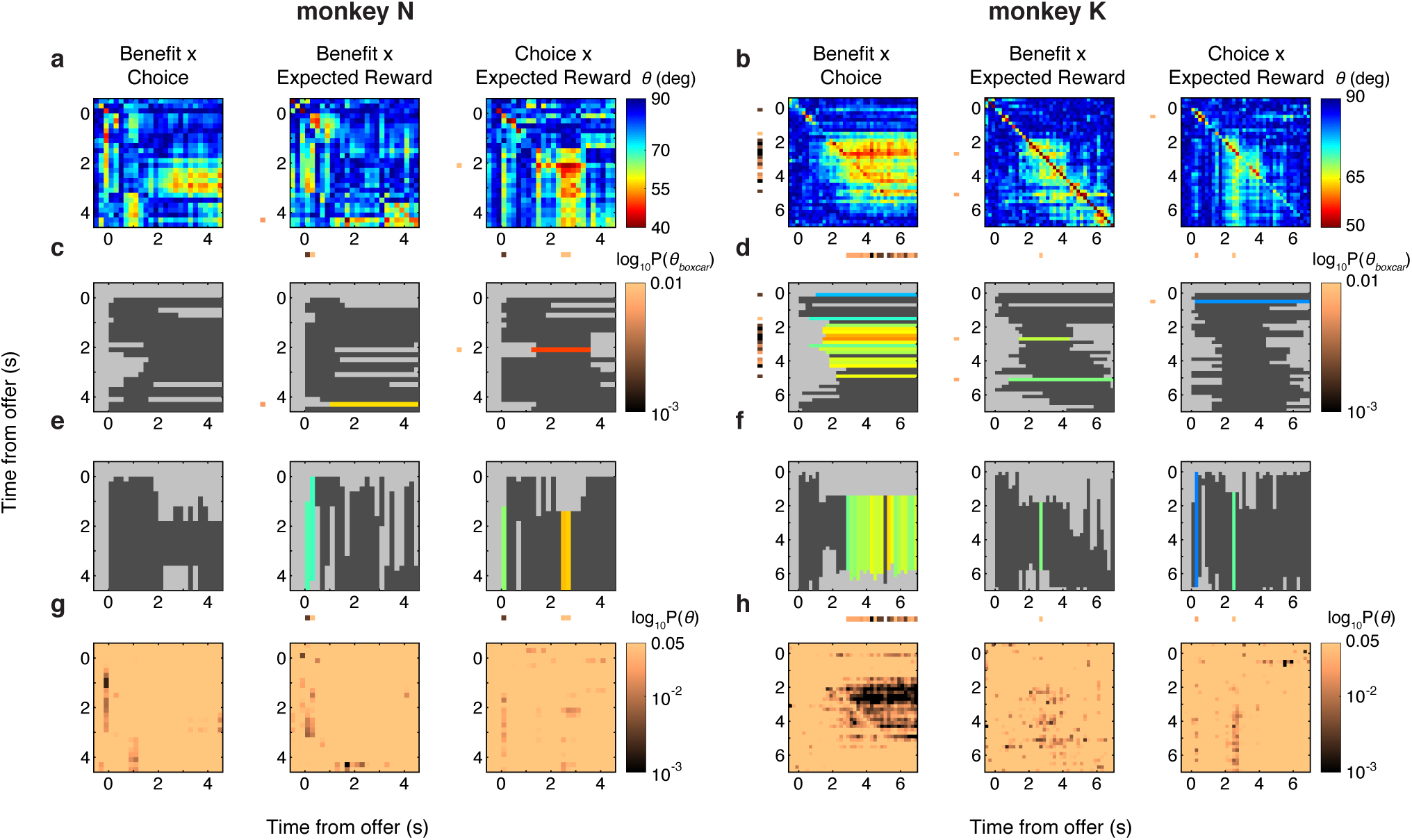
Similarity between representations of different variables. The above analyses compared dynamic low-dimensional representations (dRAs) of the same task-relevant variable. Here we examined the similarity, or “cross-talk”, between dRAs of different variables for monkeys N (left 3 columns) and K (right 3 columns). For a given animal, the dRAs between variables A and B (A x B) were compared: benefit x choice (left column), benefit x expected reward (middle column), and choice x expected reward (right column). **(a**,**b)** Angle *θ*_*ij*_ in degrees between pair of dRAs for variable A at time *i* (ordinate) and variable B at time *j* (abscissa) is given by the color of pixel *ij* (referencing right-hand color scale in (a,b)), and the times of the respective dRAs relative to offer onset are given by the pixel’s row and column position. Smaller angles (warmer colors) correspond to greater similarity between representations. The colored bands to the left of or below each panel are described below. **(c-f)** Unlike comparing dRAs of the same variable (Figure 5), the present matrix *θ* was not symmetrical. Thus, a boxcar function was fit separately to each row or column of angles in (a,b) and the period of non-zero boxcar height is indicated by the horizontal or vertical bar in the corresponding row or column of (c,d) or (e,f), respectively. Periods of significant similarity (p(*θ*_boxcar_) < 0.01) are colored, where bar color represents the boxcar height *θ*_boxcar_, i.e., magnitude of similarity, measured in degrees and referencing the right-hand color scale in (a,b). The log_10_ probability p of observing *θ*_boxcar_ is indicated by the color of the vertical or horizontal band to the left of or below the corresponding row or column in (a-f), respectively, and the colors reference the right-hand color scale in (c,d). Boxcars for which p(*θ*_boxcar_) ≥ 0.01 are shaded dark gray and the corresponding entries within the colored bands are omitted. Periods where *θ*_boxcar_ = 0 are shown in light gray. **(g**,**h)** The log_10_ probability of observing by chance the pairwise angle *θ*_*ij*_ in (a,b) is given by the color of the corresponding pixel and references the right-hand color scale in (g,h). We observed that the representations of different variables overlapped during periods of the trial. For instance, the mid-trial dRAs for benefit were similar to the mid-to-late trial dRAs for choice (a,b, left panels), though the similarity was only statistically significant for monkey K. This overlap suggested a means to compute expected reward, as a readout tuned to this mid-trial shared representation would inherently integrate offer and choice information. However, though compelling conceptually, the timing of the representations in the present study was inconsistent with this mechanism: the EXPECTED REWARD sRA detected significant RSV prior to when the benefit and choice dRAs overlapped (Figure 4c-f). Future studies may leverage the change and overlap of representations over time to understand how the network computes a given variable.

### Previous-trial low-dimensional representations

**Supplementary Figure 27.**
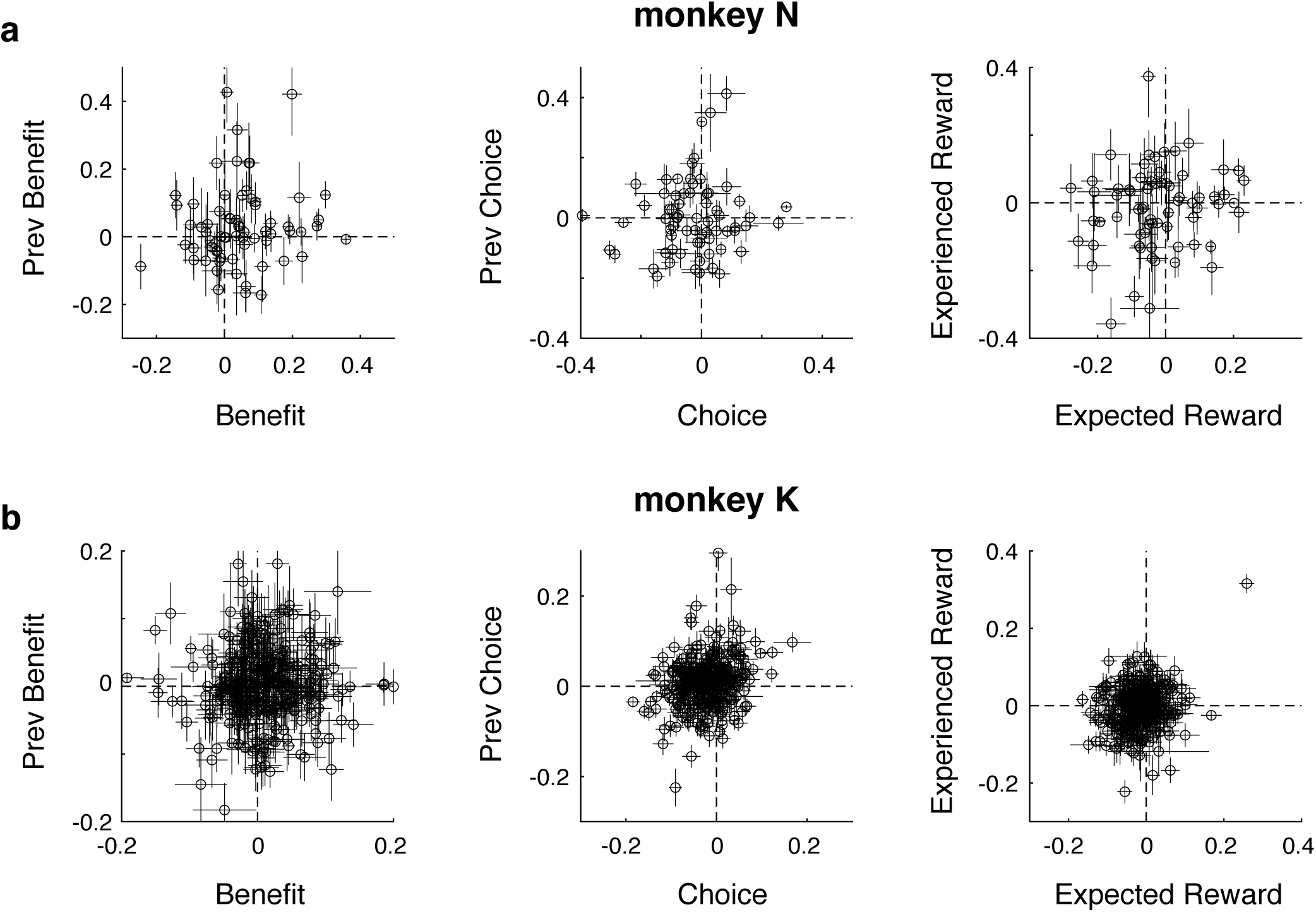
Relationship between present- and previous-trial representations. The current figure examines the relationship between the present- and previous-trial representations of the *same* variable. **(a**,**b)** Regression coefficients for the present- (abscissa) and previous-trial (ordinate) representations (i.e., sRAs) derived from all trials are shown for individual units (open circles) with associated standard deviations (horizontal and vertical error bars) derived from bootstrapped resampling of trials for monkeys N (a) and K (b). The horizontal and vertical meridians (dashed lines) indicate the location of hypothetical units representing the given variable from only the present or previous trial (but not both). Most units did not intersect these meridians, consistent with mixed representations of present- and previous-trial variables at the level of individual units. See Supplementary Table 3 for correlation coefficients and angles between present- and previous-trial sRAs. For present figure, sRAs were derived without orthogonalizing regression axes.

**Supplementary Figure 28.**
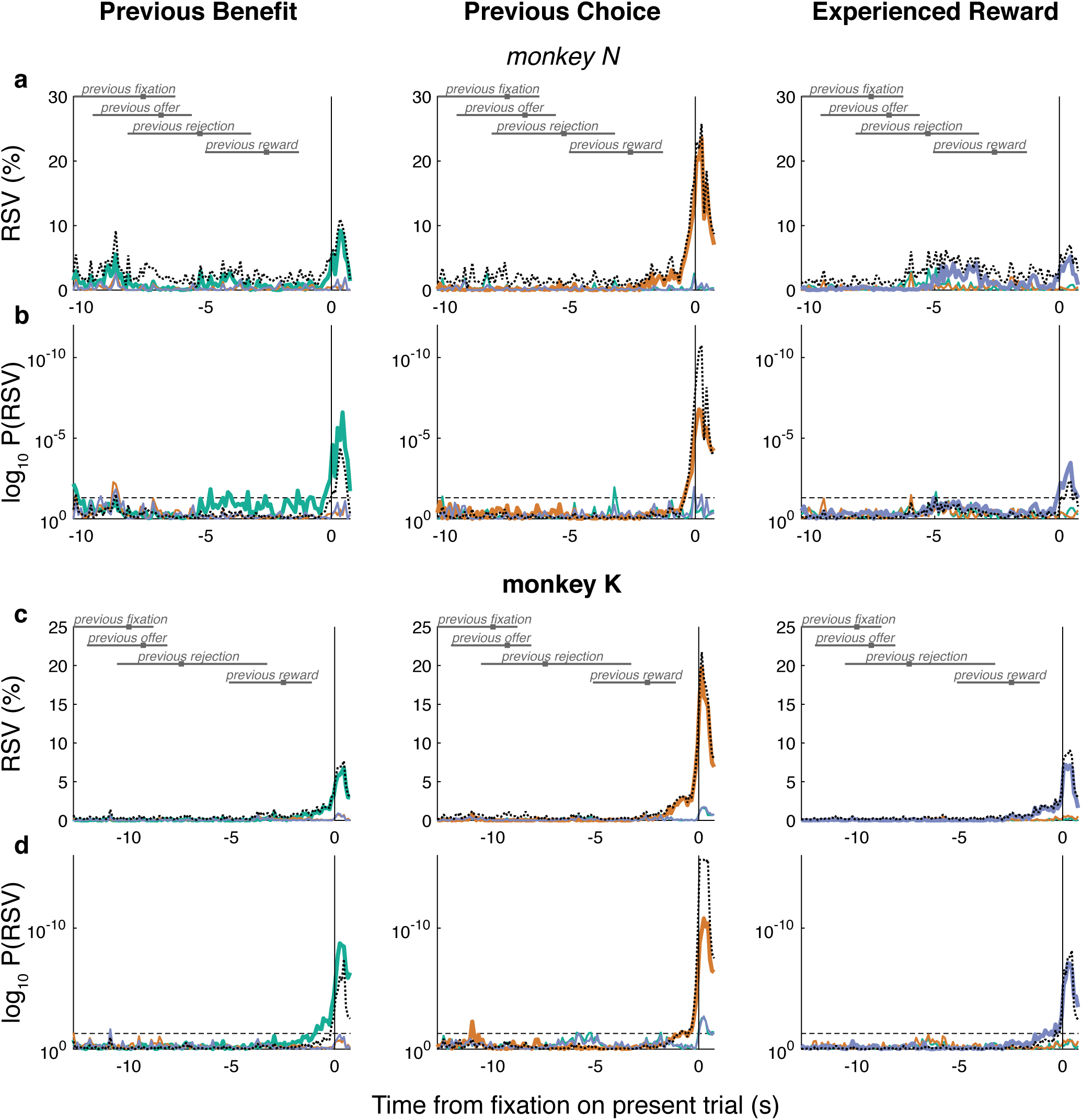
Specificity of previous-trial low-dimensional representations. **(a-d)** For detailed explanation of plotting conventions, refer to the analogous Supplementary Figure 15. Both figures demonstrate the specificity of a set of low-dimensional representations (i.e., sRAs) to their targeted variables of interest. Whereas the prior figure concerned representations of present-trial variables during the present offer and work periods, the current figure concerns representations of variables from the previous trial—previous benefit (green), previous choice (orange), and experienced reward (blue)—during a period aligned to the present fixation and extending retrospectively to the previous fixation. The on-target relevant signal variance (RSV; thick curves), off-target partial RSV (thin curves), and overall variance explained (dotted curves) for the previous-trial sRAs (columns, as labeled) are shown in (a,c) with associated log_10_ probability shown in (b,d) for monkeys N (a,b) and K (c,d). Horizontal gray bars and solid gray squares indicate the 2.5-to-97.5 percentile range and median, respectively, of previous trial events, as labeled.

**Supplementary Figure 29.**
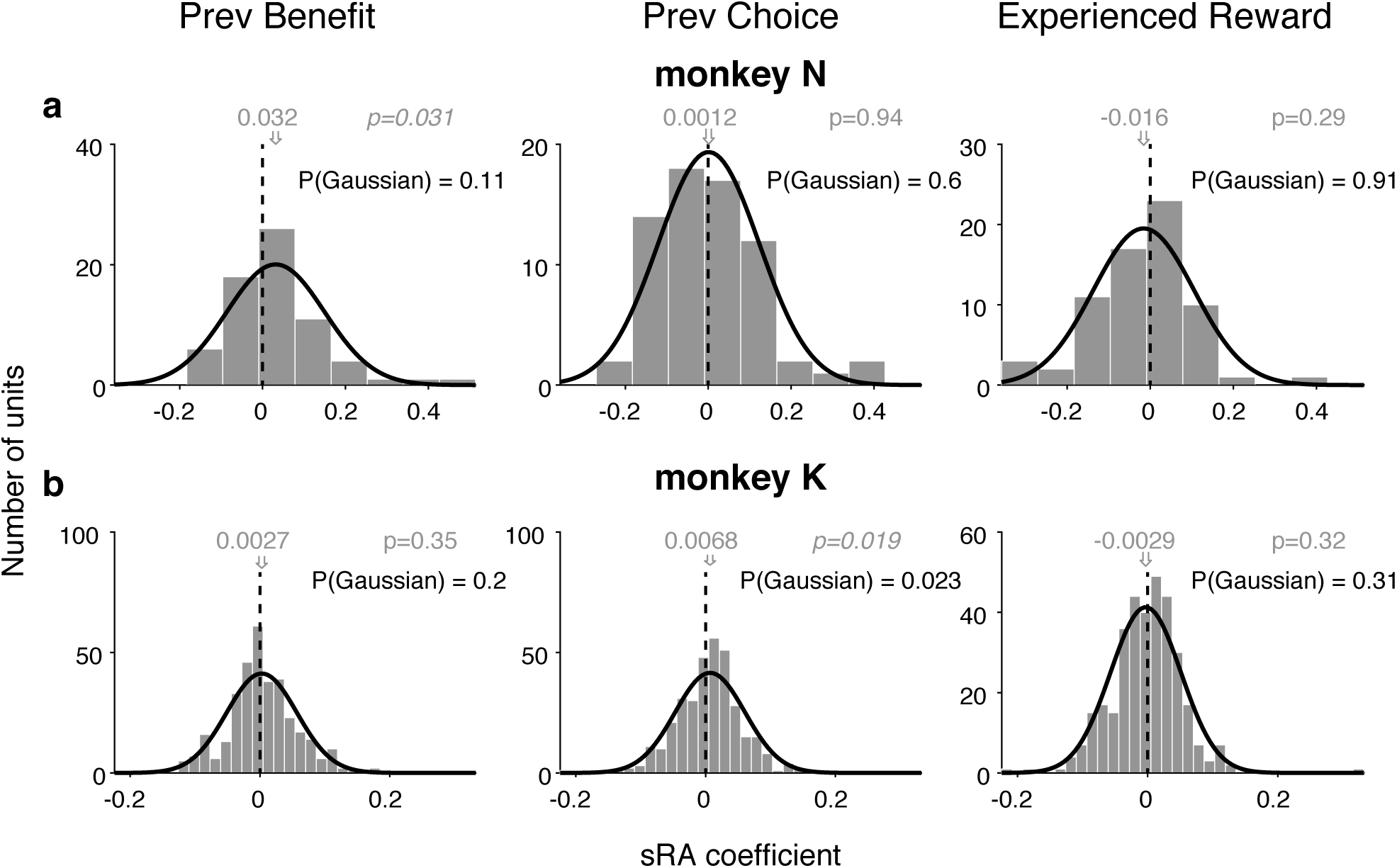
Contribution of individual units to previous-trial representations. **(a**,**b)** Histogram of regression coefficients across units that specified each unit’s contribution to the static low-dimensional representations (sRAs) of previous-trial variables: PREVIOUS BENEFIT (left panels), PREVIOUS CHOICE (middle panels), and EXPERIENCED REWARD (right panels) for monkeys N (a) and K (b). Distribution mean (gray arrow and text) and p-value (gray text; via *t*-test) of null hypothesis that mean = 0 (vertical dashed line) are shown. Gaussian functions were fit to each distribution (black curves). Probability (black text) of falsely rejecting null hypothesis that observed distribution was Gaussian was computed via two-tailed Kolmogorov–Smirnov test. No distribution except one differed significantly from Gaussian. For present figure, sRAs were derived without orthogonalizing regression axes.

**Supplementary Figure 30.**
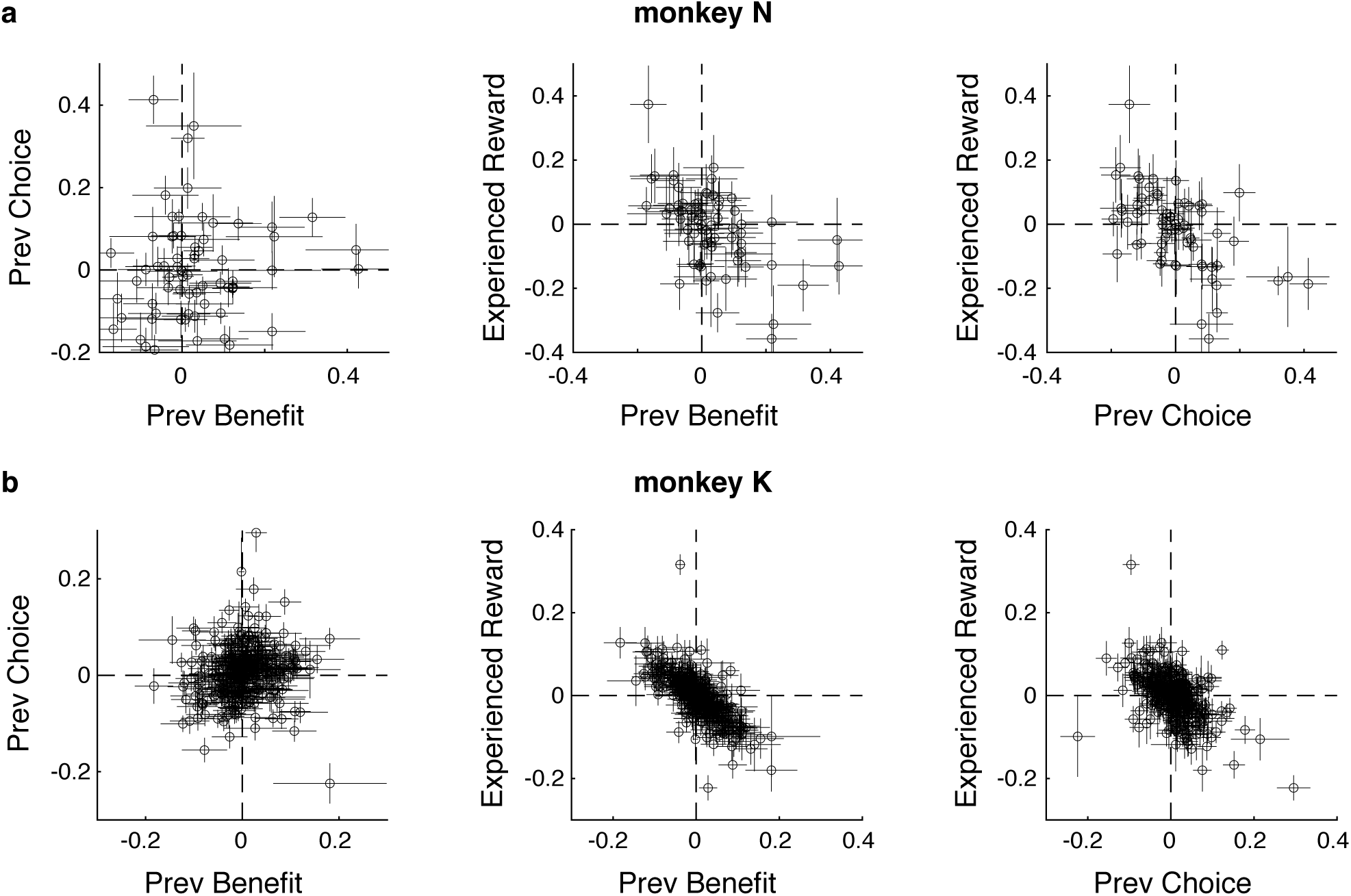
Relationship between previous-trial representations. **(a**,**b)** As for the present-trial representations in Figure 3a,c, the pairwise relationship between regression coefficients for the previous-trial representations (i.e., sRAs; abscissa and ordinate labels) derived from all trials are shown for individual units (open circles) with associated standard deviations (horizontal and vertical error bars) derived from bootstrapped resampling of trials for monkeys N (a) and K (b). The horizontal and vertical meridians (dashed lines) indicate the location of hypothetical units representing a solitary variable. Most units did not intersect these meridians, consistent with mixed representations at the level of individual units. See Supplementary Table 4 for correlation coefficients and angles between previous-trial sRAs. For present figure, sRAs were derived without orthogonalizing regression axes.

**Supplementary Figure 31.**
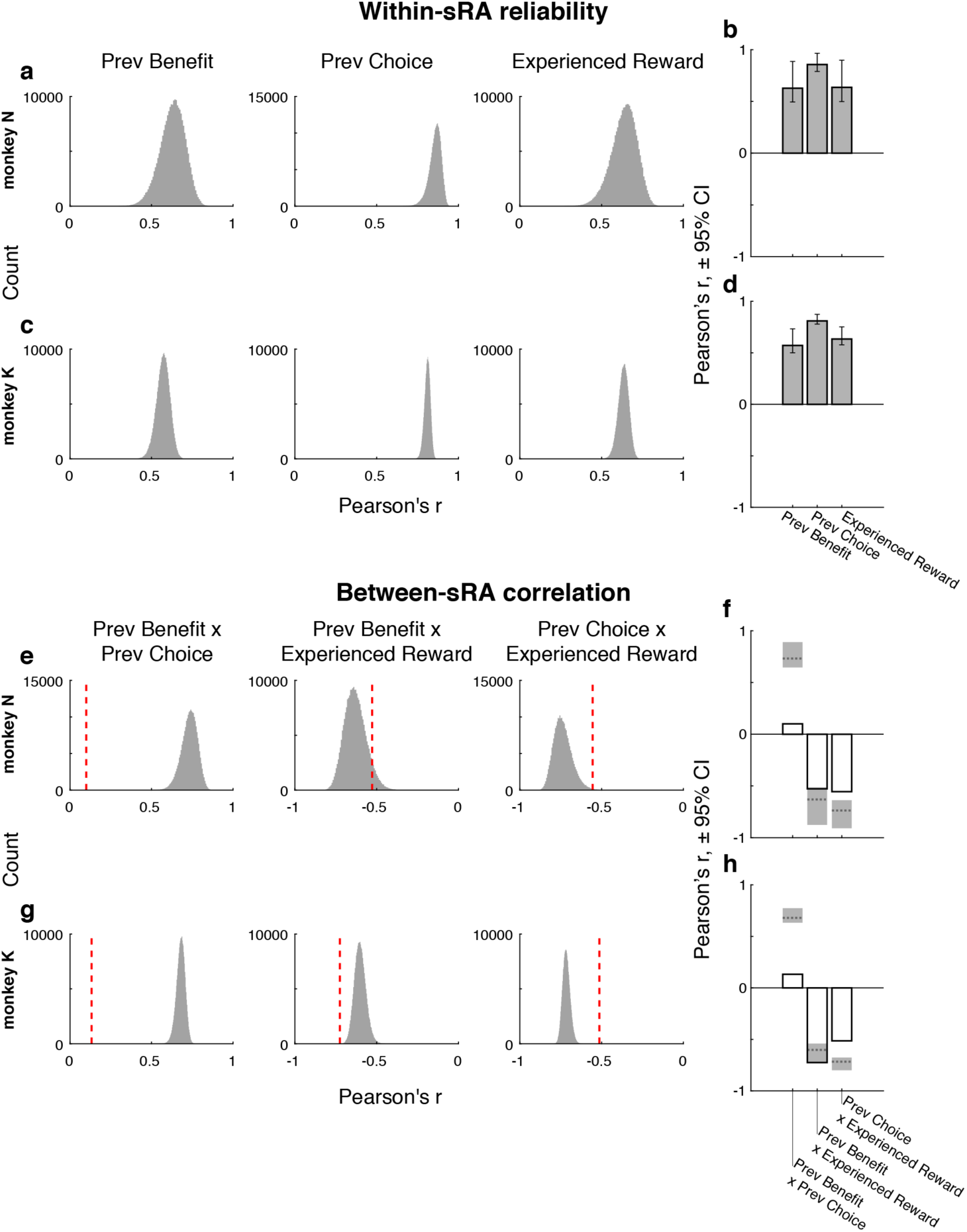
Reliability and separability of previous-trial representations. Figure refers to previous-trial representations (i.e., sRAs). For detailed description of analogous procedures on present-trial representations, see Supplementary Figure 8 for (a,c,e,g) and Figure 3b,d for (b,d,f,h). **(a**,**c)** Histograms of within-variable correlation coefficient 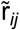 across resampled datasets are shown for monkeys N (a) and K (c), revealing high reliability of the sRAs. **(b**,**d)** The mean and 95% CI of the distributions of 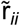 are summarized by bar height and error bars, respectively, for monkeys N (b) and K (d). **(e**,**g)** Histograms of the null model of between-variable correlation coefficients 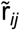 are shown for representations of each pair of variables *i* and *j* and compared to the observed correlation r_*ij*_ (vertical dashed red line) for monkeys N (e) and K (g). Null model refers to the hypothetical correlation between two perfectly correlated representations corrupted by independent noise (i.e., low reliability of the coefficients), as measured across resampled datasets. (f,h) The observed and hypothetical between-variable correlations are summarized for monkeys N (f) and K (h). Bar height corresponds to observed correlation r_*ij*_ (i.e., red vertical dashed line in (e,g)). Gray dashed horizontal line and shading indicate the mean and 95% confidence interval (CI), respectively, of the hypothetical correlation 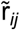 (i.e., histograms in (e,g,)). In all but one case, observed correlations were closer to zero (i.e., less correlated) than 95% of the hypothetical correlations, defining the pairs of representations as separable. See Supplementary Table 4 for within- and between-variable correlation coefficients and associated statistical tests. For present figure, sRAs were derived without orthogonalizing regression axes.

### Computing low-dimensional representations *only* in single units

**Supplementary Figure 32.**
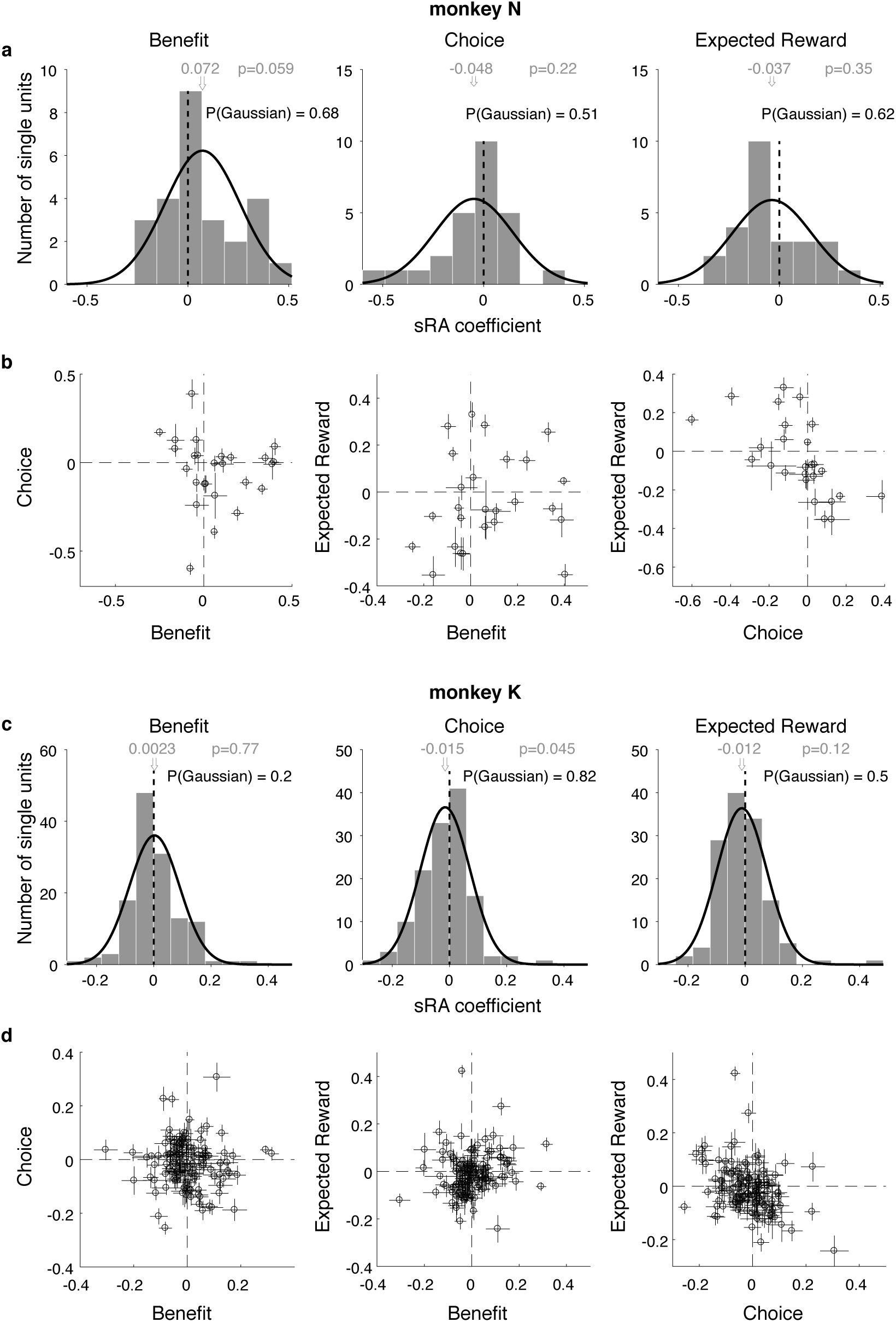
Contribution of single units to low-dimensional representations. We separately discovered the low-dimensional representations (sRAs) in a subset of data limited to single units. **(a**,**c)** As in Supplementary Figure 6a,b, histograms of regression coefficients that determined the contribution each single unit to the sRAs of BENEFIT (left panels), CHOICE (middle panels), and EXPECTED REWARD (right panels) are shown for monkeys N (a) and K (c). Distribution mean (gray arrow and text) and p-value (gray text; via *t*-test) of null hypothesis that mean = 0 (black vertical line) are shown. Probability (black text) of falsely rejecting null hypothesis that observed distribution was Gaussian was computed via two-tailed Kolmogorov–Smirnov test. No distribution differed significantly from Gaussian. **(b**,**d)** As in Figure 3a,c, the pairwise relationship between regression coefficients for task-relevant variables (abscissa and ordinate labels) derived from all trials are shown for single units (open circles) with associated standard deviations (horizontal and vertical error bars) derived from bootstrapped resampling of trials for monkeys N (b) and K (d). The horizontal and vertical meridians (dashed lines) indicate the location of hypothetical units representing a solitary variable. Most units did not intersect these meridians, consistent with mixed representations at the level of single units. See Supplementary Table 2 for correlation coefficients and angles between single-unit sRAs. For present figure, all regression coefficients were found without orthogonalizing the sRAs.

**Supplementary Figure 33.**
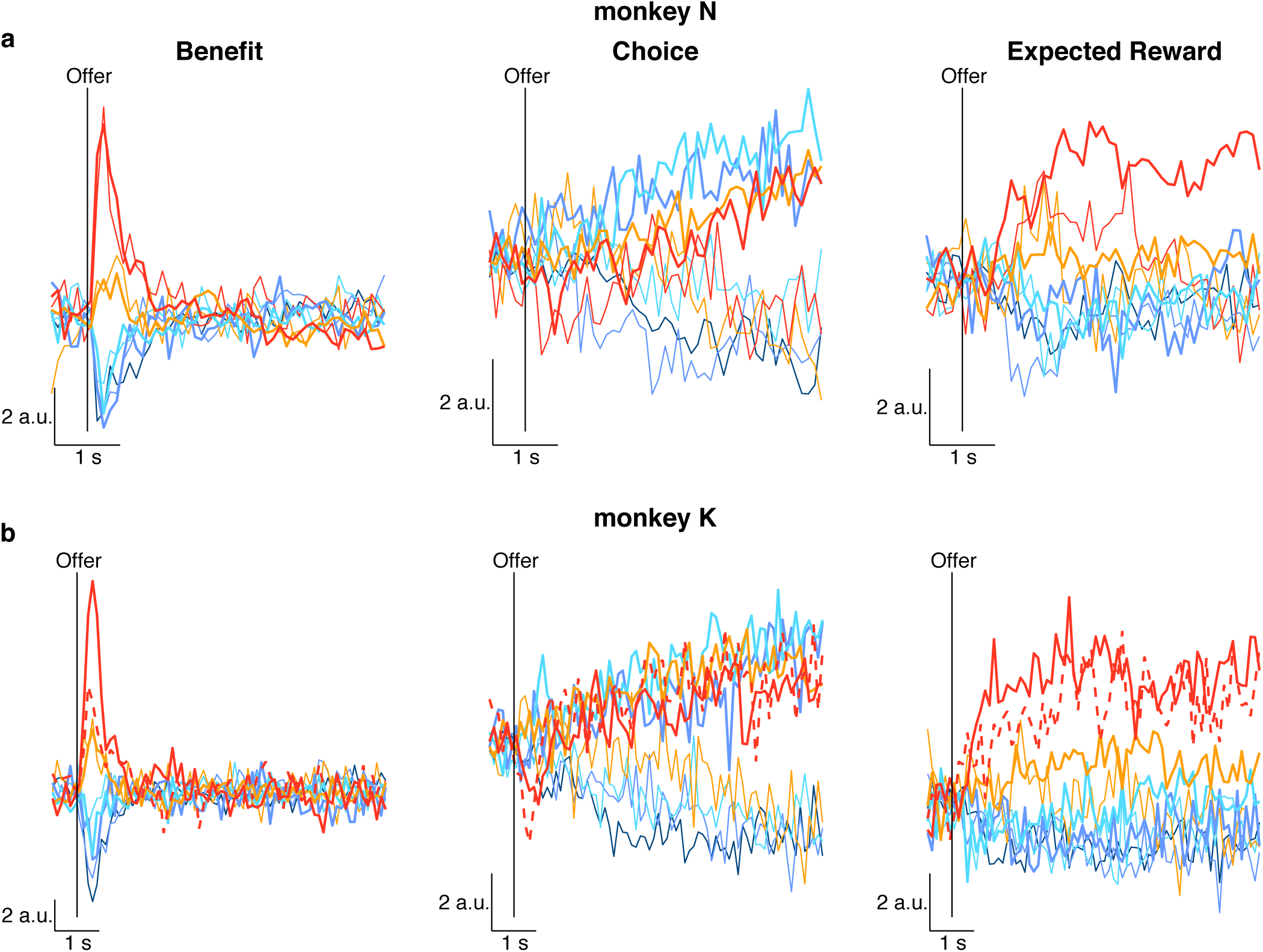
Activity of single unit-based low-dimensional representations. We separately discovered the low-dimensional representations (sRAs) in a subset of data limited to single units and projected the population response onto the sRAs of BENEFIT (left panels), CHOICE (middle panels), and EXPECTED REWARD (left panels) as a function of time from the onset of the offer period (vertical black lines) for monkeys N (a) and K (b). All conventions as in Figure 4a,b. The representation and separation of the task-relevant variables did not differ qualitatively between single units and the entire population. See Supplementary Figure 34 for additional measures.

**Supplementary Figure 34.**
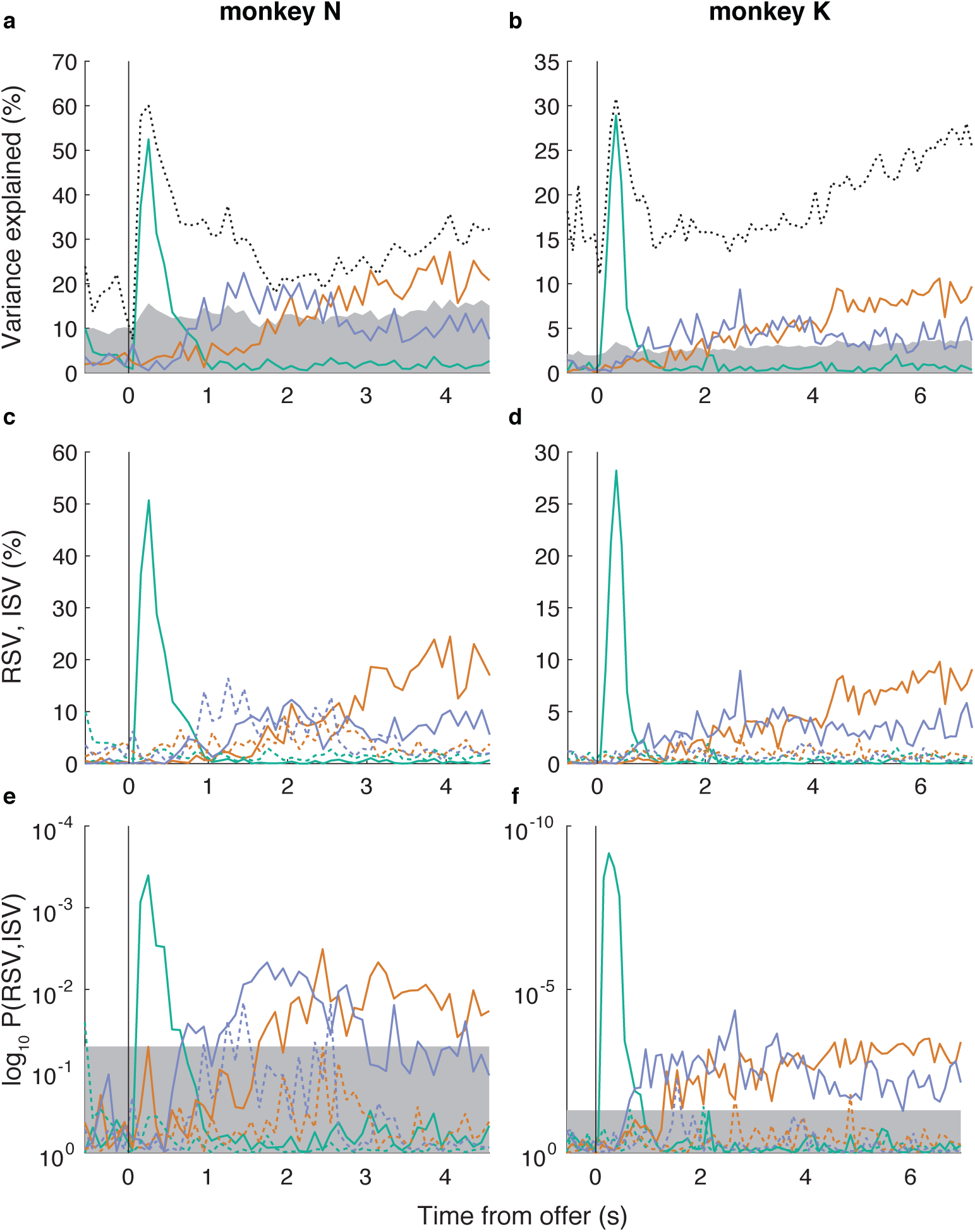
Variance explained by single unit-based low-dimensional representations. We separately discovered the low-dimensional representations (sRAs) in a subset of data limited to single units for monkeys N (a,c,e) and K (b,d,f). **(a**,**b)** As in Supplementary Figure 14a, the variance explained by these single-unit based sRAs of BENEFIT (green curve), CHOICE (orange curve), and EXPECTED REWARD (blue curve) is plotted as a function of time from the onset of the offer period (black vertical line). The area of gray shading represents the variance explained by 95% of random vectors reflecting the dimensionality of the data (redefined here for single-unit responses only). To estimate the upper-bound for variance explained, we performed principal components analysis independently at each time bin and plotted the variance explained by the top component (black dotted curve), which represented the maximal variance explained by any single, time-varying dimension. **(c**,**d)** As in Figure 4c,d, we plotted the relevant and irrelevant signal variance (RSV and ISV, solid and dashed lines, respectively) explained by the sRAs (colors as above) with respect to the targeted variable of interest (e.g., RSV explained by the BENEFIT sRA with respect to the benefit variable). **(e**,**f)** As in Figure 4e,f, we plotted the log_10_ probability of RSV and ISV based on the respective null distribution generated from random vectors. Area of gray shading represents p > 0.05.

## Supplementary Tables

**Supplementary Table 1.** Significant encoding of task-relevant variables in individual units.

| Variable A | Monkey N |  |  | Monkey K |  |  |
| --- | --- | --- | --- | --- | --- | --- |
|  | BENEFIT | CHOICE | EXPECTED REWARD | BENEFIT | CHOICE | EXPECTED REWARD |
| 1. Included* | 67 | 67 | 67 | 340 | 340 | 340 |
| 2. Significant | 45 (67%) | 34 (51%) | 42 (63%) | 137 (40%) | 148 (44%) | 106 (31%) |
| *Single Units* |  |  |  |  |  |  |
| 3. Included* | 25 | 25 | 25 | 129 | 129 | 129 |
| 4. Significant | 17 (68%) | 11 (44%) | 19 (76%) | 137 (40%) | 148 (44%) | 106 (31%) |
| Overlap with Variable B | CHOICE | EXPECTED REWARD | BENEFIT | CHOICE | EXPECTED REWARD | BENEFIT |
| 5. Both sig.<br>obs. / exp. | 25 (37%)<br>22.8 (34%) | 22 (33%)<br>21.3 (32%) | 31 (46%)<br>28.2 (42%) | 68 (20%)<br>59.6 (18%) | 58 (17%)<br>46.1 (14%) | 55 (16%)<br>42.7 (13%) |
| 6. Both non-sig. | 13 (19%)<br>10.8 (16%) | 13 (19%)<br>12.3 (18%) | 11 (16%)<br>8.21 (12%) | 123 (36%)<br>115 (34%) | 144 (42%)<br>132 (39%) | 152 (45%)<br>140 (41%) |
| 7. A sig. only | 20 (30%)<br>22.2 (33%) | 12 (18%)<br>12.7 (19%) | 11 (16%)<br>13.8 (21%) | 69 (20%)<br>77.4 (23%) | 90 (26%)<br>102 (30%) | 51 (15%)<br>63.3 (19%) |
| 8. B sig. only | 9 (13%)<br>11.2 (17%) | 20 (30%)<br>20.7 (31%) | 14 (21%)<br>16.8 (25%) | 80 (24%)<br>88.4 (26%) | 48 (14%)<br>59.9 (18%) | 82 (24%)<br>94.3 (28%) |
| 9. $p(\chi^2, df)$ | 0.26 (1.3, 1) | 0.73 (0.12, 1) | 0.13 (2.3, 1) | 0.062 (3.5, 1) | 0.0051 (7.8, 1) | 0.0034 (8.6, 1) |

**Supplementary Table 2.**
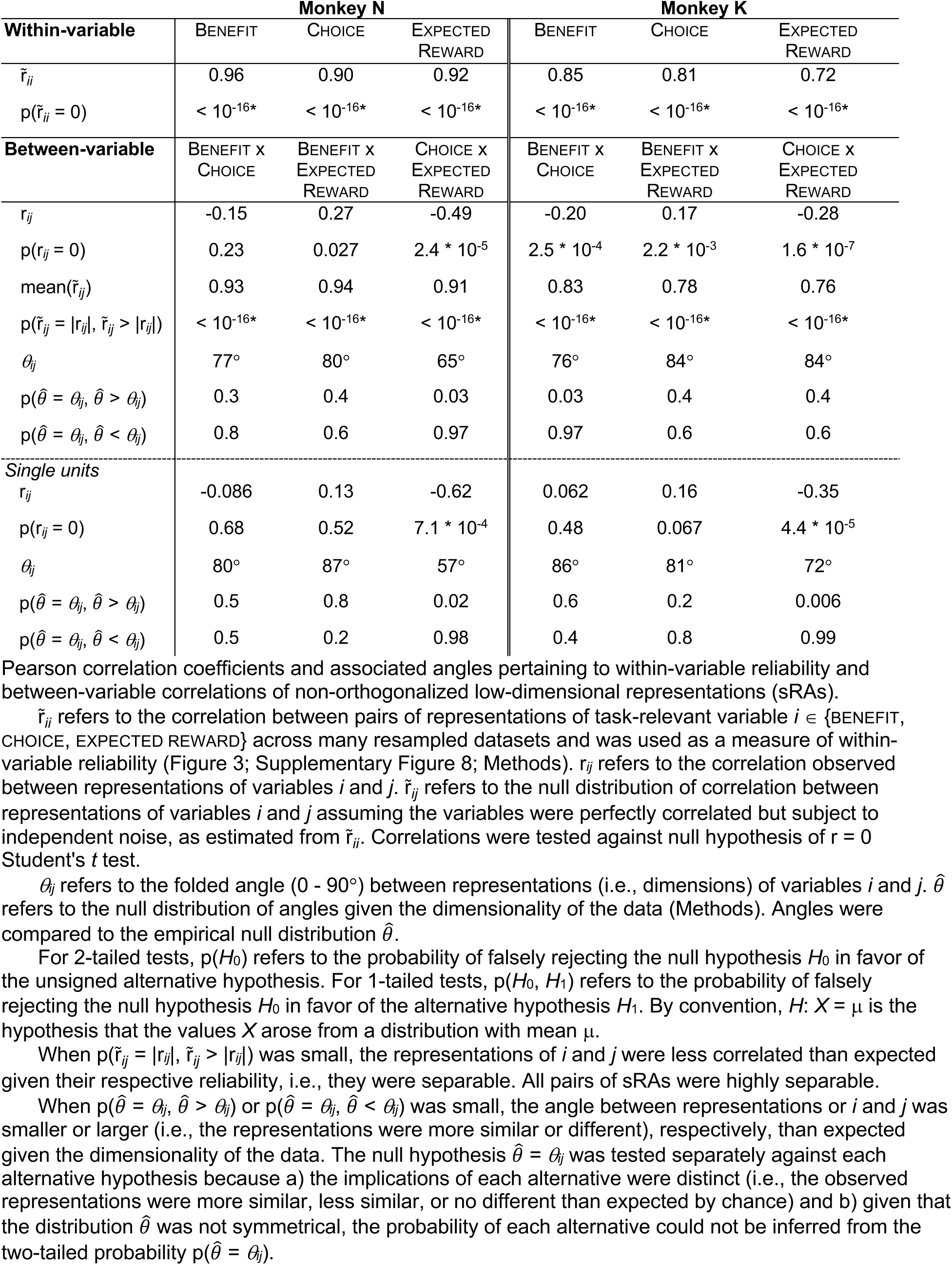
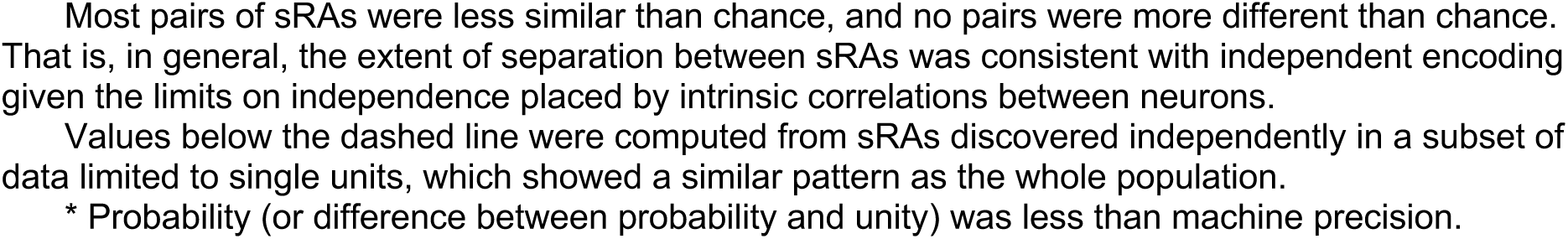
Reliability of and correlation between low-dimensional representations.

**Supplementary Table 3.** Correlation between present- and previous-trial low-dimensional representations.

|  | Monkey N |  |  | Monkey K |  |  |
| --- | --- | --- | --- | --- | --- | --- |
|  | BENEFIT X<br>PREVIOUS<br>BENEFIT | CHOICE X<br>PREVIOUS<br>CHOICE | EXPECTED<br>REWARD X<br>EXPERIENCED<br>REWARD | BENEFIT X<br>PREVIOUS<br>BENEFIT | CHOICE X<br>PREVIOUS<br>CHOICE | EXPECTED<br>REWARD X<br>EXPERIENCED<br>REWARD |
| $r_{ij}$ | 0.12 | 0.097 | 0.15 | 0.021 | 0.23 | 0.17 |
| $p(r_{ij} = 0)$ | 0.35 | 0.43 | 0.24 | 0.70 | $1.5 * 10^{-5}$ | $1.4 * 10^{-3}$ |
| $\theta_{ij}$ | 78° | 85° | 80° | 88° | 81° | 80° |
| $p(\hat{\theta} = \theta_{ij}, \hat{\theta} > \theta_{ij})$ | 0.3 | 0.6 | 0.3 | 0.8 | 0.09 | 0.05 |
| $p(\hat{\theta} = \theta_{ij}, \hat{\theta} < \theta_{ij})$ | 0.8 | 0.4 | 0.7 | 0.3 | 0.9 | 0.95 |

**Supplementary Table 4.**
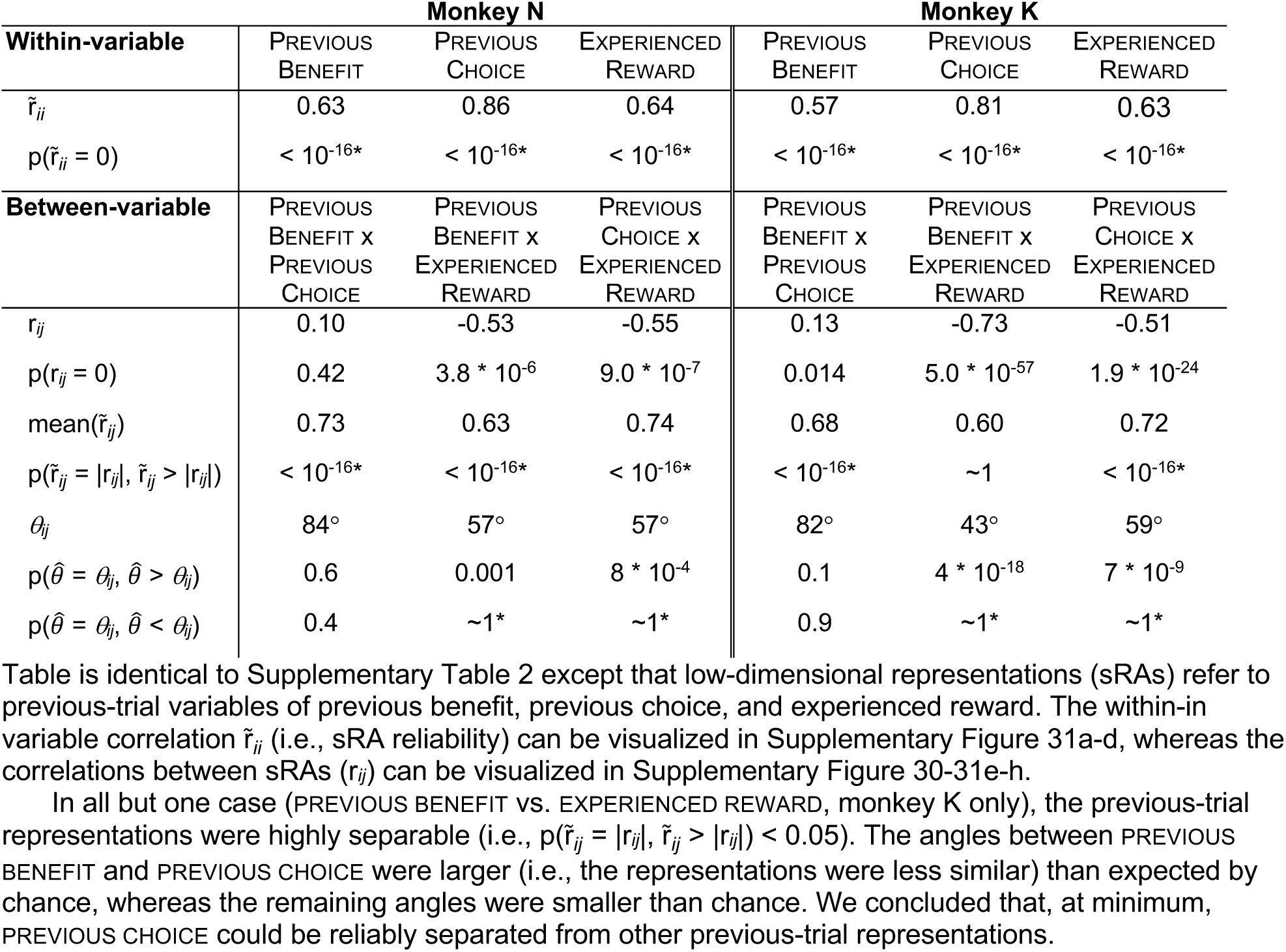
Reliability of and correlation between previous-trial low-dimensional representations.

## Appendix 1

### Solving single-trial problem with trial-average responses

Here, we derive the trial-average least squares problem that is equivalent to the single-trial problem in Eq. 4, thereby allowing us to express the objective function in Eq. 5 in terms of trial-average responses while weighting each response by the number of trials contributing to its average.

Let *R*_*n*_(*r, t*), as defined in Eq. 4, be the firing rate of unit *n* at trial *r* and time *t*. We drop the *t* and *n* notation here as this derivation applies for all times and units. Define an experiment condition c as the unique combination of the *K* predictors **P** ∈ *ℝ*^*K*^ (i.e., unique combination of benefit, choice and expected reward). Let 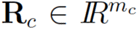 be the vector of firing rates across trials of condition c (*m*_*c*_: is the number of trials of condition c).

The single trial model in Eq. 4 can thus be written as 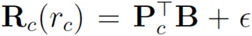, where B ∈ *ℝ*^*K*^ is the vector of model regression coeffiecients (∀*r*_*c*_ ∈ {1,…, *m*_*c*_} and ∀_*c*_ ∈ {1,…, *C*}) and *ϵ* is indepedent Gaussian noise. To solve the best coefficients 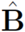, we solve the following linear regression problem:

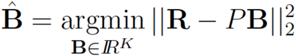

Where 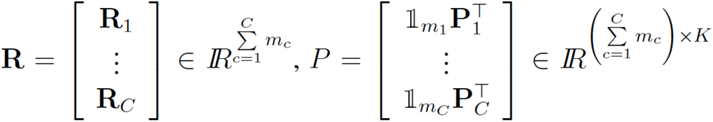 and 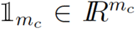 is a vector of length *m*_*c*_ with all elements equal to one, which repeats **P**_*c*_ for the number of trials with that unique set of predictors.

The solution of this problem is the least squares solution:

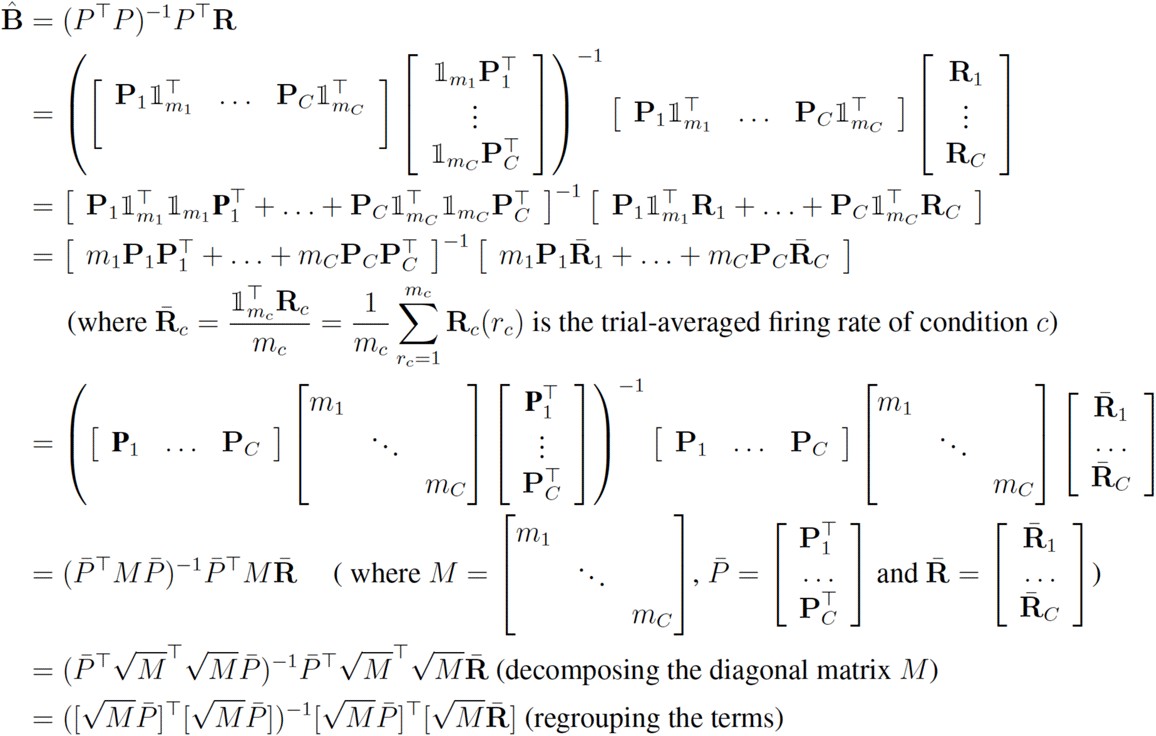

The expression above is the least square solution of the following problem:

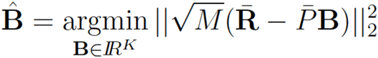

The above problem only includes trial-averaged terms. Thus,

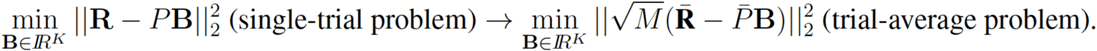

Note that the above expression pertains to a single neuron. To extend to ***N*** neurons, the vectors 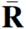 and **B** are extended to ***C*** × ***N*** and ***K*** × ***N*** matrices, respectively, with neurons arranged in columns. For a given neuron, the ***C*** × ***C*** diagonal matrix ***M*** is then collapsed such that the elements on the diagonal form a ***C*** × 1 vector, and then, across neurons, extended to the ***C*** × ***N*** matrix M, with neurons arranged in columns. The difference between the measured and predicted responses (analogous to 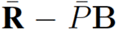 above) is then multiplied element-wise by 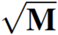. Finally, to arrive at the expression in Eq. 5, matrices pertaining to neural responses should be transposed such that neurons are arranged in rows.

